# Multiple chromosomal inversions contribute to adaptive divergence of a dune sunflower ecotype

**DOI:** 10.1101/829622

**Authors:** Kaichi Huang, Rose L. Andrew, Gregory L. Owens, Kate L. Ostevik, Loren H. Rieseberg

## Abstract

Both models and case studies suggest that chromosomal inversions can facilitate adaptation and speciation in the presence of gene flow by suppressing recombination between locally adapted alleles. Until recently, however, it has been laborious and time-consuming to identify and genotype inversions in natural populations. Here we apply RAD sequencing data and newly developed population genomic approaches to identify putative inversions that differentiate a sand dune ecotype of the prairie sunflower (*Helianthus petiolaris*) from populations found on the adjacent sand sheet. We detected seven large genomic regions that exhibit a different population structure than the rest of the genome and that vary in frequency between dune and non-dune populations. These regions also show high linkage disequilibrium and high heterozygosity between, but not within haplotypes, consistent with the behavior of large inversions, an inference subsequently validated in part by comparative genetic mapping. Genome-environment association analyses show that key environmental variables, including vegetation cover and soil nitrogen, are significantly associated with inversions. The inversions co-locate with previously described “islands of differentiation,” and appear to play an important role in adaptive divergence and incipient speciation within *H. petiolaris*.

## INTRODUCTION

Genetic differentiation between differently adapted populations can be highly variable across the genome. During the process of adaptive divergence, genomic regions under selection will display strong differentiation, while ongoing gene flow between populations will homogenize other regions, generating heterogeneous patterns of genomic divergence (Wu, 2001; Nosil, Funk, & Ortiz-Barrientos, 2009). Large islands of differentiation, namely “genomic islands of divergence”, are commonly seen in recently diverging populations, ecotypes, and species, including well-known examples in *Rhagoletis* (Feder, Chilcote, & Bush, 1988), *Anopheles* (Turner, Hahn, & Nuzhdin, 2005), *Heliconius* (Nadeau et al., 2012) and *Helianthus* (Andrew & Rieseberg, 2013). The causes of these large islands are not fully understood (although see McGaugh & Noor, 2012; Berg et al., 2017). It has been proposed that divergence hitchhiking, in which gene exchange is reduced adjacent to a locus under strong divergent selection, could generate large regions of differentiation, but the conditions under which it occurs are limited (Via, 2012; Feder & Nosil, 2010). Chromosomal inversions represent another possible explanation for such islands because they can suppress recombination and impede gene flow across large genomic regions (Butlin, 2005; Hoffman & Rieseberg, 2008).

Inversions have long been viewed as important in local adaptation and speciation (Wellenreuther & Bernatchez, 2018; Dobzhansky & Sturtevant, 1938). One primary reason is that, by suppressing recombination, inversions can establish and maintain favorable combinations of locally adapted alleles, despite gene flow with non-adapted populations (Rieseberg, 2001; Kirkpatrick & Barton, 2006). The critical importance of inversions in local adaptation has been revealed by emerging studies that document the association of inversions with adaptive traits within species (Feder, Roethele, Filchak, Niedbalski, & Romero-Severson, 2003; Lowry & Willis, 2010; Kirubakaran et al., 2016; Wellenreuther & Bernatchez, 2018 for review). Beyond their role in adaptation, inversions can preserve alleles that cause intrinsic genetic incompatibilities in hybrids, and facilitate the accumulation of new incompatibilities, thereby aiding species’ persistence in the face of gene flow (Noor, Grams, Bertucci, & Reiland, 2001; Navarro & Barton, 2003). Finally, inversions can establish linkage between locally adapted alleles and those causing assortative mating, which is typically required in models of speciation with gene flow (Felsenstein, 1981; Trickett & Butlin, 1994; Servedio, 2009).

Much of what we know about inversions (at least until very recently), comes from studies of Dipteran flies, whose very large larval salivary gland chromosomes permit detection of inversions from chromosome banding patterns (Krimbas & Powell, 1992). However, in most other organisms, more time-consuming and/or expensive methods have been required, such as analyses of meiotic configurations (Heslop-Harrison, 2013), comparative genetic mapping (Kirubakaran et al., 2016), Hi-C sequencing (Dixon et al., 2018), optical mapping (Tang, Lyons, & Town, 2015), paired-end mapping (Lamichhaney et al., 2016), or long-read sequencing. The laboriousness and/or expense of these methods have hindered our understanding of the frequency and importance of inversions in natural populations. Recently, population genomic approaches have been applied to detect potential inverted regions, including methods based on linkage disequilibrium (LD) (Faria et al., 2019; Arostegui, Quinn, Seeb, Seeb, & McKinney, 2019) and local population structure (Li & Ralph, 2019). The LD approach takes advantage of the expectation that inversions will create high LD between (but not within) inversion haplotypes. The local population structure approach assumes that the lack of gene flow between inversion haplotypes will lead to systematic differences in patterns of genetic relatedness between inverted and collinear regions. Such differences can be detected by conducting windowed analyses of population structure across the genome (Li & Ralph, 2019). Both methods offer an efficient means for identifying putative inversions and estimating their frequency in natural populations.

In this study, we focus on the genetic architecture of adaptation in a dune-adapted ecotype of the prairie sunflower *Helianthus petiolaris* Nutt. This widespread annual sunflower inhabits sandy soils in the Central and Southwestern USA. However, in the Great Sand Dunes National Park and Preserve (GSD), Colorado, an ecotype of this species occurs in active sand dunes. This dune ecotype differs from conspecific populations, which are abundant on the sand sheet below the dunes, for a number of ecologically relevant phenotypic traits, including seed size, branching, and root architecture (Andrew, Ostevik, Ebert, & Rieseberg, 2012). Despite its origin less than 10,000 years ago (Andrew, Kane, Baute, Grassa, & Rieseberg, 2013), multiple reproductive barriers isolate the two ecotypes, including strong extrinsic selection against immigrants and hybrids, conspecific pollen precedence, as well as a weak crossability barrier (Ostevik, Andrew, Otto, & Rieseberg, 2016). Nonetheless, substantial and asymmetric gene flow have been reported between dune and non-dune populations (Andrew et al., 2012), as predicted by models of isolation with gene flow. Moreover, genetic differentiation between the ecotypes is largely restricted to several large genomic regions while background divergence is extremely low (Andrew & Rieseberg, 2013), making it a good system to study the evolution of genomic islands of divergence. The underlying mechanism for these large regions of high divergence was not previously determined, but chromosomal inversions represent a leading hypothesis given their ability to impede introgression, as well as the high rates of chromosomal evolution reported for *Helianthus* (Burke et al., 2004; Ostevik, Samuk, & Rieseberg, 2019).

Our analyses complement a recently submitted study from our group on the genetic architecture of local adaptation across three sunflower species (Todesco et al., 2019). In that study, we used whole genome shotgun sequence (WGS) data to sample genetic variation across the ranges of three sunflower species, including *H. petiolaris*. The study detected numerous large haplotypes in all three species that co-varied with ecologically relevant phenotypic, climate, and soil variation. Further analyses show that many of the haplotypes (but not all), were associated with structural variation, including inversions. One population from GSD (ten individuals) was included in this study, and it appeared to be enriched for structural variants. Thus, we also wished to validate this observation with more extensive sampling from GSD and surrounding regions, as well as to exploit a comprehensive data set on local variation in soil fertility and plant cover on the dune and surrounding sand sheet to better assess the role of inversions in divergent adaptation with gene flow.

Specifically, we employ RAD sequence data previously generated for this system (Andrew et al., 2013) and apply a local population structure approach to detect and genotype putative inversions in this system. We also conduct additional population genomic analyses (including LD analyses) and develop two genetic maps (one for each ecotype) to further validate these inferences. Lastly, we search for associations between the genotypic data and key environmental factors, including soil nutrient availability and vegetation coverage. We address four main questions: 1) Can structural variants such as inversions be detected with RAD sequencing data? 2) If so, are they enriched in the dune habitat at GSD as previously suggested? 3) Likewise, do they correspond closely to the genomic islands of differentiation (i.e., high *F*_ST_ regions) previously reported between dune and non-dune sunflowers? and 4) Lastly, is there evidence that inversions contribute importantly to adaptive divergence in this system?

## MATERIALS AND METHODS

### Plant materials and RAD sequencing

Our study employs the plant materials and RAD sequencing (Baird et al., 2008) data set previously reported by Andrew & Rieseberg (2013) and Andrew et al. (2013). Twenty populations from dune, non-dune and intermediate habitats in the GSD were sampled (Andrew et al., 2013, Supporting Information Table S1), and five unrelated individuals from each of the 20 populations were subjected to RAD sequencing by Floragenex (Portland, OR) using the restriction enzyme *Pst*I. All samples were barcoded and sequenced with at least 60 bp reads, with a subset sequenced with 80 bp reads. The first 5 bp covering the restriction site and relatively low-quality 20 bp at 3’ end of the 80 bp reads were trimmed with PRINSEQ v0.20.4 (Schmieder & Edwards, 2011), yielding reads with equal length of 55 bp, to avoid biases in alignment due to sequences of different lengths.

### SNP calling

We re-called SNPs from the RAD sequencing data since much better reference genomes are now available for cultivated sunflower (*Helianthus annuus*), a close relative of *H. petiolaris*. Briefly, RAD sequences were aligned to reference genome Ha412HOv2.0 with bwa mem v0.7.17 (Li, 2013) using the default settings. Variant calling was performed with the Genome Analysis Tool Kit v4.0.8.1 (GATK; DePristo et al., 2011). Sample alignments were processed with the GATK HaplotypeCaller and samples were jointly genotyped using GATK’s GenotypeGVCFs chromosome by chromosome. Variants of all chromosomes were later merged with MergeVcfs in Picard tools (http://broadinstitute.github.io/picard/). Only bi-allelic SNPs were selected for downstream analyses. SNPs were filtered with GATK VariantFiltration with filter expression “QD < 4.0 || FS > 20.0 || MQ < 40.0 || MQRankSum < −5.0” and individual genotypes with depth less than 30 were set as missing. Loci that were non-variant or varied only due to singletons after filtering, as well as those with > 40% missing data, were excluded from the data set. Finally, SNPs with excess heterozygosity were filtered with GATK’s ‘VariantFiltration’ filter expression “ExcessHet < 20.0” to avoid misalignment on paralogous regions.

Because the new reference genome provides physical locations of the SNPs and has much more complete chromosome coverage compared to the one used by Andrew and Rieseberg (2013), we re-calculated Weir and Cockerham’s *F*_ST_ (Weir, 1996) between dune and non-dune ecotypes with VCFtools (Danecek et al., 2011) to examine genetic divergence across the new reference genome and re-localize regions of divergence.

### Local population structure analysis

We analyzed patterns of population structure across the genome using the R package “lostruct” (Li & Ralph, 2019), in order to detect regions of abnormal population structure that might be generated by chromosomal inversions. The genome was divided into non-overlapping windows with size of 50 SNPs and principle component analysis (PCA) was calculated for each window to reflect local population structure. To measure the similarity of patterns of relatedness between windows, Euclidean distances between matrices were calculated for the first two principle components (PCs) and then mapped using multidimensional scaling (MDS) into 40-dimensional space. Different window sizes were tested to reach the best balance between signal and noise. The SNP data set was converted to BCF format with BCFtools v1.9 (Li, 2011) before input to lostruct.

To identify localized genomic regions with extreme MDS values, we first defined outlier windows as those with absolute values greater than 4 standard deviations from the mean across all windows for each of the 40 MDS coordinates. We then tested whether outlier windows were chromosomally clustered with 1,000 permutations of windows over chromosomes to evaluate differences from random expectation where outliers are randomly distributed among chromosomes. For each MDS coordinate with more than 4 outlier windows, we selected the first chromosome with a significant excess of outliers (p < 0.01) for further examination. For each coordinate, outlier windows that deviated in different directions were examined separately. Adjacent outliers with less than 4 windows between them were kept as a cluster. In cases where the same chromosome had outlier clusters across multiple MDS coordinates, we calculated Pearson’s product moment correlation coefficient between the MDS coordinates using sample genotype matrices and collapsed the ones with correlation > 0.8 by selecting the coordinate with the larger number of outliers. The coordinates of the putative inversions were defined by the start position of the first outlier window to the end position of the last outlier window.

While inversions are a major driver of MDS outliers detected by lostruct (Li & Ralph, 2019), MDS outliers can be generated by other processes as well, such as linked selection. Therefore, we performed a series of additional analyses to look for additional population genomic signatures of inversions. Due to suppressed recombination, haplotype blocks with different orientations should evolve largely independently, resulting in distinct nucleotide differences between them. Therefore, for an inversion segregating in a population, a PCA of population structure should divide the samples into three distinct groups representing the two inversion haplotypes, with heterozygotes between the haplotypes forming an intermediate cluster. To test this, we calculated PCAs with SNPrelate (Zheng et al., 2012) using all SNPs from each putative inversion. To identify the composition of groups of genotypes, we used the R function “kmeans” with the method developed by Hartigan and Wong (1979) to perform clustering on the first PC, using the maximum, minimum and middle of the range of PC scores as the initial cluster centers. The discreteness of the clustering was evaluated by the proportion of the between-cluster sum of squares over the total. The K-means cluster assignment was used as the genotype of the sample.

If the groups detected in the PCA represent homozygotes and heterozygote for the orientations, we expect the central group to have high heterozygosity relative to the other two groups. For each region identified, we extracted all variable sites across the outlier windows and calculated the proportion of heterozygous sites over the total as heterozygosity for each individual in each group identified by k-means clustering.

To examine the effect of recombination suppression of the putative inversions, intrachromosomal linkage disequilibrium (LD) was calculated among all SNPs with minor allele frequencies > 5%. Pairwise LD (R^2^) values were calculated using PLINK v1.9 (Chang et al., 2015; Purcell et al., 2007) for each chromosome with all samples. Values of SNPs were grouped into 1 Mb windows and the second largest R^2^ value was plotted using ggplot2 (Wickham, 2016). For chromosomes with MDS outlier regions, R^2^ was also calculated with individuals homozygous for the more common orientation only.

Only the regions displaying clustering of three distinct groups in the PCA with higher heterozygosity in the middle group and high LD were kept as putative inversions in downstream analyses. For each region, allele frequency differences between ecotypes were estimated using “prop.test” in R and the genotype frequency for each population was plotted onto a map of land cover classification downloaded from Multi-Resolution Land Characteristics Consortium (https://www.mrlc.gov/) at 30-m resolution.

### Genetic map construction

Genetic maps of dune and non-dune ecotypes were generated using F1 testcross mapping to validate our inversion detection approach. Pollen from a single dune plant (seed collected from population 1300) and a single non-dune plant (seed collected from a new population at Latitude 37.724, Longitude −105.718) from GSD, was used to fertilize individuals of the male sterile *H. annuus* HA89cms cultivar, which is highly homozygous. For each cross, the HA89cms individuals that bore the most seeds (100-150 seeds) were selected to produce the F1 mapping populations. Loci that are heterozygous in a wild parent are expected to segregate 1:1 in the corresponding F1 population, permitting the generation of a genetic map. DNA was extracted from germinated F1 seeds or, when germination failed, directly from seeds. Barcoded genotyping-by-sequencing (Poland, Brown, Sorrells, & Jannink, 2012) libraries were prepared using the restriction enzymes *Pst*I and *Msp*I. A depletion step with Duplex-Specific Nuclease (DSN; Evrogen, Moscow, Russia) was conducted on the libraries to reduce the proportion of repetitive sequences, including plastid DNA (Todesco et al., in preparation). The libraries were sequenced on an Illumina Hiseq 4000 instrument to produce paired end, 100 bp reads (Illumina, San Diego, CA, USA). Samples were demultiplexed using a custom Perl script that also removed barcode sequences. FASTQ files were examined for quality but not trimmed. Raw reads were aligned to the Ha412HOv2.0 reference genome using NextGenMap v0.5.2 (Sedlazeck, Rescheneder, & Von Haeseler, 2013) and variants were called using GATK v4.0.8.1 as described above for the RAD sequences. Only SNPs were kept and filtered with the expression “QD < 15.0 || FS > 20.0 || MQ < 40.0 || MQRankSum < −5.0”, and individual genotypes with depth less than 30 were set as missing. Loci that were invariant after filtering and had a genotype missing rate > 50% were excluded.

Genetic maps were built using R/qtl (Broman, Wu, Sen, & Churchill, 2003) and R/ASMap (Taylor & Butler, 2017). Individuals with fewer than 50% markers genotyped were excluded, as were duplicate markers, markers with less than 50% of individuals scored, and markers with extreme segregation patterns (genotype frequency <0.3 or >0.7). The “mstmap.cross” function was used to construct linkage groups (LGs) with the remaining markers using a *p*-value of 10^-15^, which was chosen to minimize false linkages. Because marker phase was unknown prior to mapping, mirror image LGs were generated initially, and the function “switchAlleles” was used to reverse genotype scores for such LGs. Markers with segregation distortion *P*-value < 0.05 and missing rate < 0.1 were pulled aside from the map, and those with more than three double crossovers and markers with extreme (> 2 standard deviation) segregation distortion within a 21-marker window were removed using custom functions. LGs with less than two markers were discarded. Some less extreme markers that were originally placed aside were then pushed back into the map and the markers were filtered again with the same criteria. This step was done twice to reintroduce markers with segregation distortion *P*-value < 0.01, missing rate < 0.3 and the ones with segregation distortion *P*-value < 0.001 and missing rate < 0.5. The function “calc.errorlod” was also used to filter genotyping errors. Finally, very small (1-5 markers) LGs were discarded, leaving 17 LGs for each ecotype.

Due to sparse marker density on the LG that corresponded to chromosome 5 after filtering, markers that mapped to chromosome 5 on the *H. annuus* reference genome were extracted and genetic mapping was repeated using less stringent parameters. Markers that were located at the far end of LG 5, and those that disturbed synteny, were removed because they might represent misaligned markers from other chromosomes. This remapping was conducted for both dune and non-dune mapping populations and the new LGs were included in downstream genetic map comparisons.

To compare marker orders, we took advantage of the fact that SNP markers were called against the Ha412HOv2.0 reference genome. Homologous reference chromosomes for each linkage groups were identified based on physical positions of markers and prior knowledge of the location of translocations between *H. petiolaris* and *H. annuus* (Ostevik et al., 2019). For each putative inversion, we asked whether markers from that region differed in order or genetic distance with respect to the reference genome and/or between ecotypes.

### Genome-environment association analysis

To further assess the role of putative inversions in dune adaptation in the ecotype, as well as to identify the environmental variables that might be driving divergent selection pressures, we used data on soil nutrient availability and vegetation coverage for each population to conduct genome-environment association (GEA) analysis.

The collection and estimation of these measurements have been described in detail in a previous study (Andrew et al., 2012). Additional composite variables of soil or cover data were generated by PCA and the first three PCs (soil PC1-3 and cover PC1-3) were used in the analyses. The vegetation coverage data were arcsine-square-root-transformed and all measurements were standardized prior to PCA.

The GEA analysis was performed using BayPass v2.1, which explicitly accounts for the covariance structure among the population allele frequencies resulting from population demography (Gautier, 2015). We further filtered the SNPs by missing rate < 10% and minimum allele frequency > 10% and generated a data set of SNP frequencies for all populations. Population structure was estimated by running BayPass under the core model mode with all filtered SNPs. The covariance matrix from this analysis was then used as a control for population structure to evaluate associations of SNPs with each environmental variable. For each SNP, a Bayes factor (BF) was computed under the standard covariate model using the default importance sampling estimator approach. Scaling was performed for each environmental variable using the “-scalecov” option. Due to missing soil data in population 970, the analysis was run separately for soil variables and coverage variables.

To further examine the associations between the putative inversions and environmental variables, we also performed a GEA analysis in which putative inversions were treated as single bi-allelic loci. A SNP data set excluding SNPs from within the putative inversions was used to estimate the covariance matrix to control for the effects of the MDS outlier regions on population structure. Bayes factors were calculated using the same core model mode in BayPass as described above.

To calculate a significance threshold, we simulated pseudo-observed data (POD) with 1,000 SNPs using the “simulate.baypass” function implemented in BayPass with the covariance matrix generated under the core model, and analyzed the newly created POD for each environmental variable as described above. The top 1% quantile of the POD BFs was computed as the threshold for significance.

## RESULTS

### SNP Calling

Using a high quality reference genome for cultivated *H. annuus*, 87.0% of RAD sequences were aligned on average, and after variant calling with GATK, a total of 260,478 variable sites were scored. Filtering produced a data set of 37,930 high-quality bi-allelic SNPs across 17 chromosomes of the reference, which corresponds to approximately 12 sites per Mbp. This compares favorably to the 11,727 SNPs that could be positioned on chromosomes in our previous analyses (Andrew & Rieseberg, 2013).

Analysis of patterns of genetic divergence between the dune and non-dune ecotypes yielded similar results to the previous study (Andrew & Rieseberg, 2013): low overall *F*_ST_ and high heterogeneity among sites with the largest clusters of outliers found on chromosomes 5,9 and 11 (Figure 1). However, highly divergent regions are more distinct and contiguous in the present study due to the larger number of SNPs and better genome assembly. In addition, a distinctive island can now be seen on the end of chromosome 7, which was not detected in the previous analysis

**FIGURE 1.**
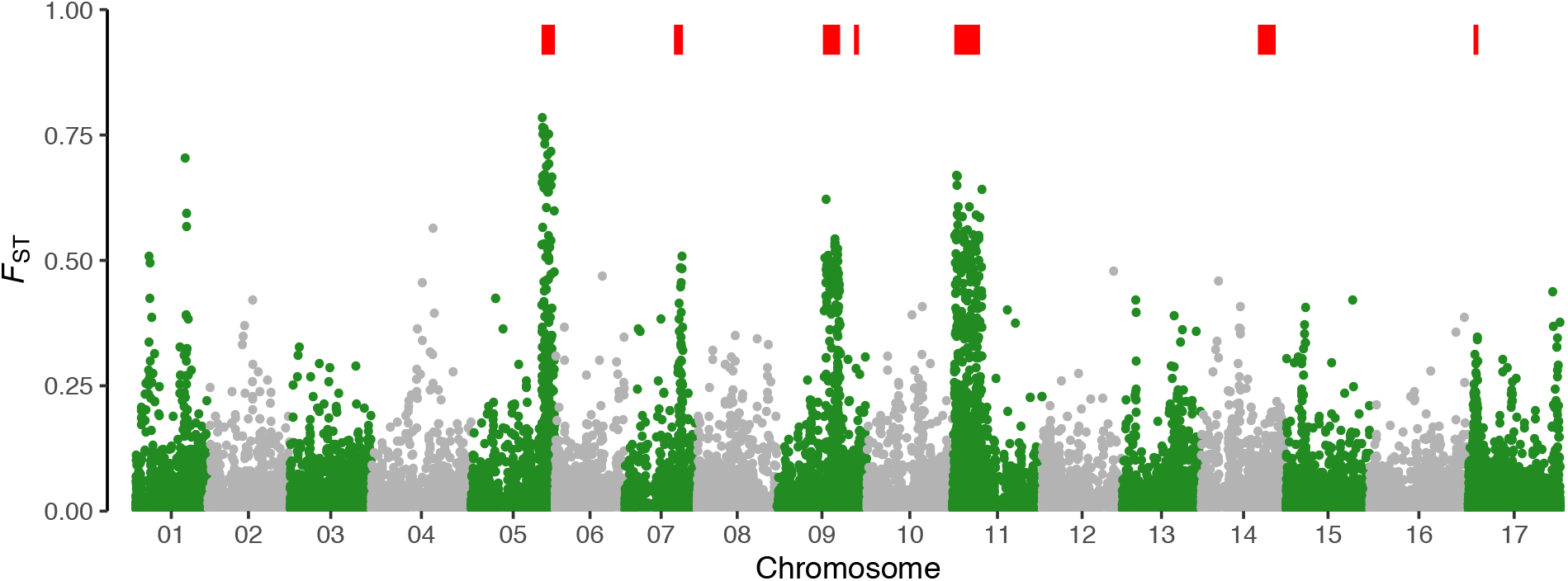
Weir and Cockerham’s *F*_ST_ between dune and non-dune ecotypes, as well as location of putative inversions (indicated by red bars on top)

### Detection of putative chromosomal inversions

Using a window-based local population structure analysis implemented in “lostruct,” and our outlier discovery approach, we identified a total of 9 clusters of MDS outliers with our RAD SNPs (Table 1, Figure 2).

**TABLE 1.**
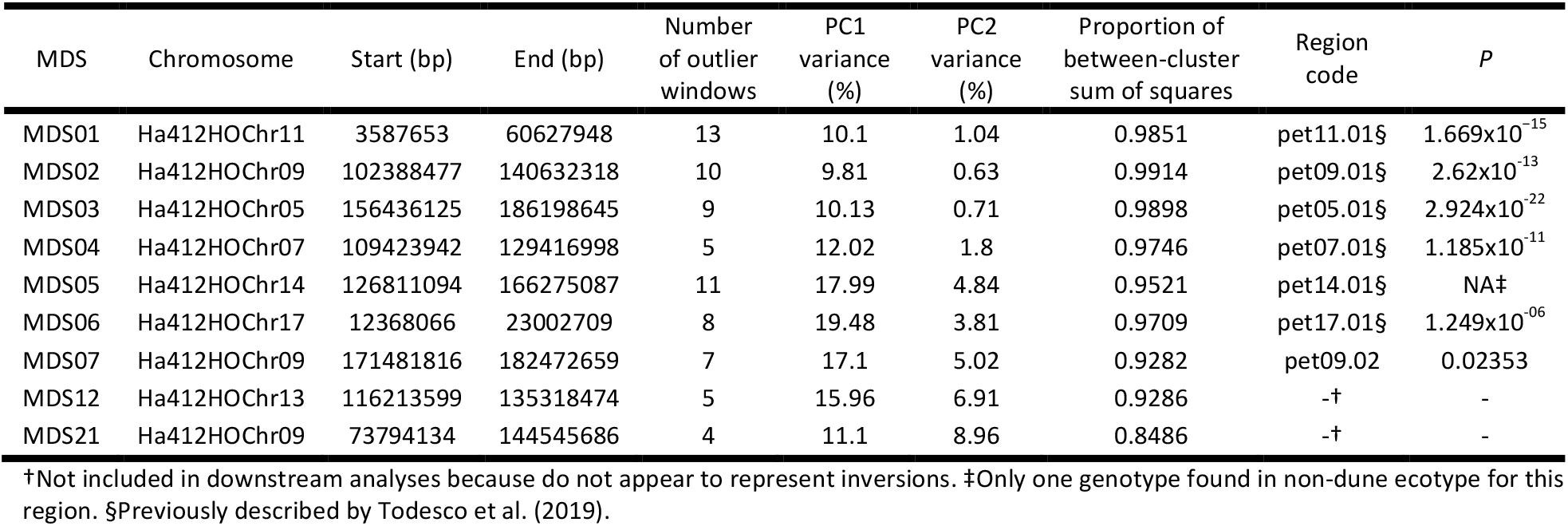
Clusters of MDS outliers obtained with “lostruct”. MDS coordinates for which the outlier regions were identified, reference chromosomes with start and end positions of MDS outlier clusters, numbers of MDS outlier windows, variance explained by PC1 and PC2 in PCA of outlier regions, proportions of between-cluster sum of squares in k-means clustering, codes used in main text for putative inversions, as well as *P*-values of the “prop.test” for haplotype frequency differences between ecotypes are shown

**FIGURE 2.**
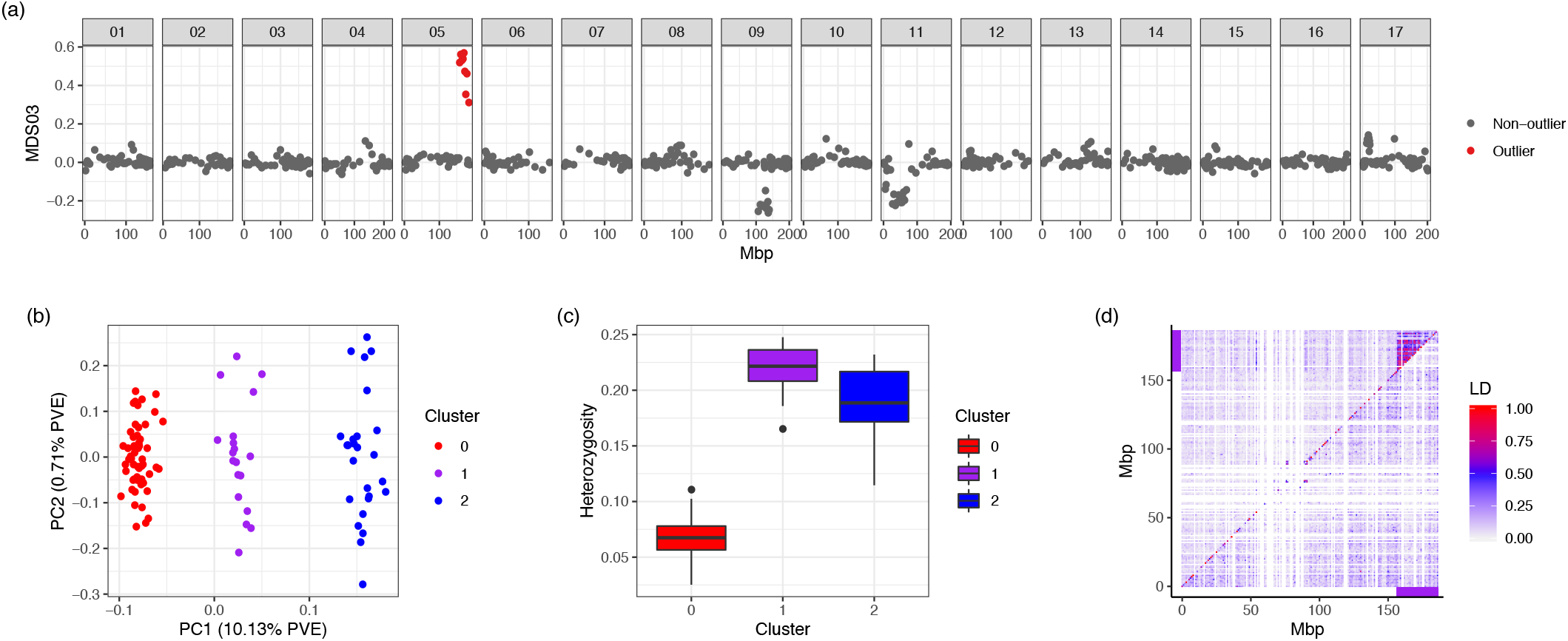
Characterization of the MDS outlier region on chromosome 5 (pet05.01). (a) Genome plot of corresponding MDS values across 17 reference chromosomes. Each dot represents a window of 50 SNPs, and outlier windows are highlighted in red. (b) PCA based on SNPs from outlier region. Three clusters identified using k-means clustering correspond to two homozygote groups (blue and red) and a heterozygote group (purple). (c) Heterozygosity for each of the groups identified in PCA. (d) LD plot for chromosome 5. Upper triangle with all individuals and lower triangle with only individuals homozygous for the more common orientation. SNPs were summarized and the second highest R^2^ values were presented in 1 Mbp windows. Purple bars represent the location of the inversion

In PCAs of most outlier regions, individuals were aggregated into three discrete groups on the first PC, which explained much more variation than the second PC (Table 1, Figure 2b, Supporting Information Figure S1). The discreteness was supported by the high (>0.9) proportion of the between-cluster sum of squares over the total in k-means clustering (Table 1). Moreover, in most regions, heterozygosity of the middle group was significantly higher than within the other two groups (Figure 2c, Supporting Information Figure S1). These patterns are consistent with the presence of two clusters of individuals that are homozygous for alternative inversion haplotypes and an intermediate cluster of individuals that are heterozygous for the inversion haplotypes with no or very little recombination between them. Two exceptions were found, including one on chromosome 13 for MDS12, where samples formed only two groups in the PCA and the expected pattern of heterozygosity was not observed. Likewise, samples did not form distinct clusters for outlier region MDS21 on chromosome 9 (Table 1, Supporting Information Figure S1). Note that outlier region for MDS21 encompasses that of MDS02, which does act like a legitimate inversion, as well as an upstream region of the chromosome that generally does not.

Almost all outlier clusters were also characterized by high LD, and almost all large regions of high LD across the genome were identified as outlier regions in our analyses. For the two outlier clusters that did not form three distinct groups in the PCA, the MDS12 outlier region on chromosome 13 was characterized by high LD. Thus, we cannot rule out the possibility that this is an inversion, but that heterozygotes are rare and genotypes mis-classified. MDS21 includes a large high LD region, which represents the MDS02 outlier region, as well as a smaller high LD region at the start. Possibly the latter represents a small inversion that is in partial LD with the MDS02 outlier region. There also were a handful of very small high LD regions (e.g., on Chromosome 15 from 119-123 Mbp) that might represent inversions, but they did not pass our stringent criteria for MDS outliers. Lastly, while high LD was detected for the outliers when compared across all samples, recombination was not restricted within the homozygous group (Figure 2d, Supporting Information Figure S1, S2), except for MDS21. These results are consistent with the role of inversions in altering recombination in heterozygotes while recombination in homozygotes remains unaffected.

Overall, seven of the outlier clusters showed clustering of three distinct groups in PCA, higher heterozygosity in the middle group and high LD across the outlier region, and were kept as putative inversions for downstream analyses (Table 1). All the putative inversions, except one on chromosome 9 (pet09.02), overlapped substantially with large haplotypes identified in *H. petiolaris* using WGS data over its entire geographic distribution (Todesco et al. 2019; Table 1). These 7 putative inversions occurred on 6 chromosomes. A majority of them were located near the end of chromosomes, while the putative inversion on chromosome 7 (pet07.01) and the larger one on chromosome 9 (pet09.01) resided in the middle sections of the chromosomes (Figure 1). Each of the putative inversions contained at least 5 MDS outlier windows (i.e. 250 SNPs) and their sizes varied between 11 and 57 Mbp (Table 1).

All of the putative inversions displayed significant allele frequency differences between dune and non-dune ecotypes (*P* ranges from 0.024 for pet09.02 to 2.92×10^-22^ for pet05.01, Table 1), but the distributions of the genotypes for each inversion were variable. For several putative inversions, the sand dunes are enriched with samples homozygous for one of the orientations (cluster 0 or cluster 2 identified by k-means clustering) (e.g., pet11.01 and pet05.01), while others showed more heterozygotes in the dunes (e.g., pet09.01) (Figure 3, Supporting Information Figure S3). For pet14.01, the “dune” orientation was not found in the non-dune habitat, although this orientation has a low frequency among samples, with only one individual identified as homozygous (Figure 3).

**FIGURE 3.**
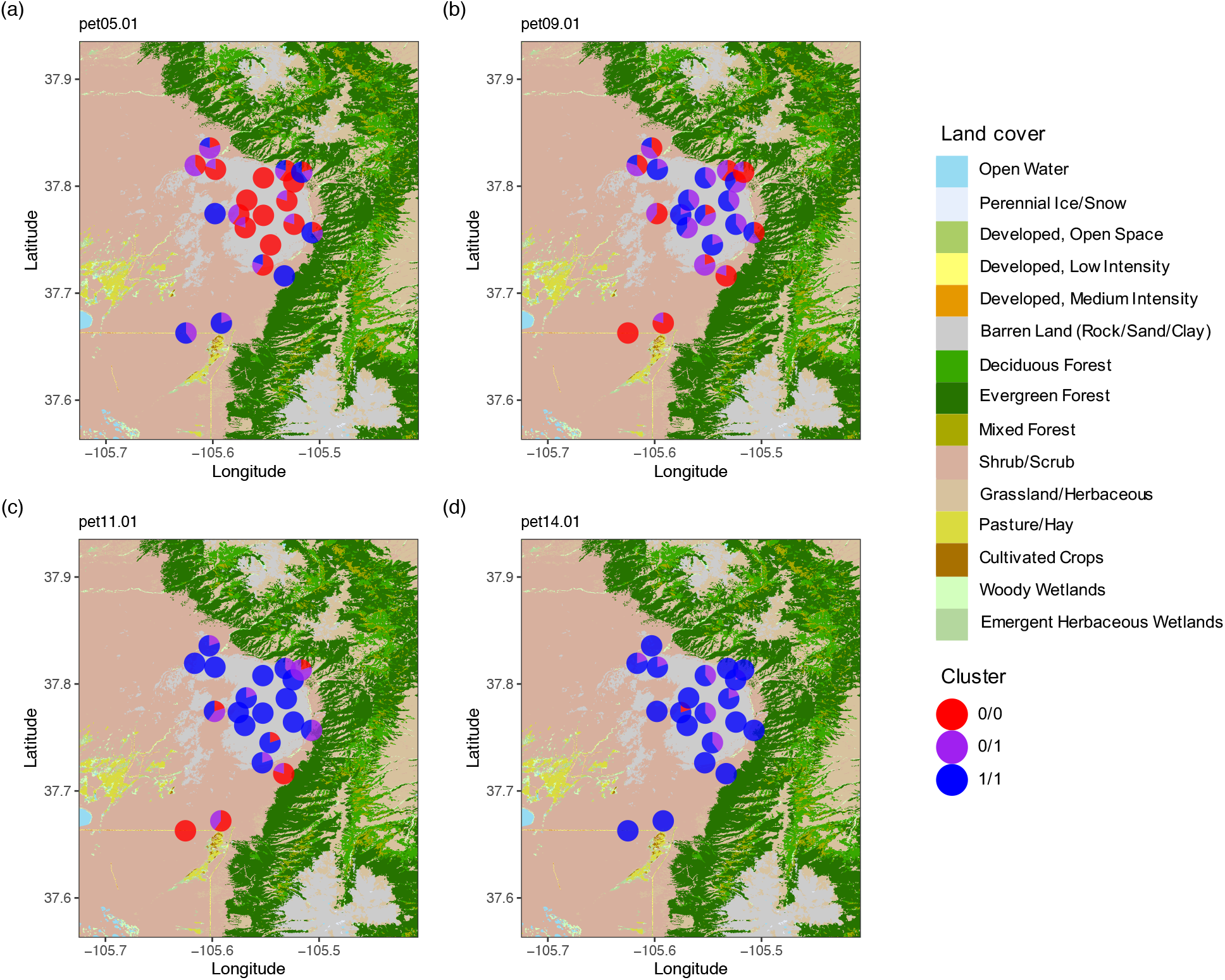
Map of Great Sand Dune National Park showing genotype distributions of (a) pet05.01, (b) pet09.01, (c) pet11.01 and (d) pet14.01. Genotypes are based on k-means cluster assignment in PCA. One of the haplotypes (inversion orientations) is more commonly found in dunes, which are represented by barren land surrounded by shrubby habitat in the map. Land cover classification downloaded from Multi-Resolution Land Characteristics Consortium (https://www.mrlc.gov/) at 30-m resolution

Most of the putative inversions were associated with regions of high *F*_ST_ between dune and non-dune ecotypes (Figure 1), especially in pet05.01, pet07.01, pet09.01 and pet11.01, where the largest divergence between ecotypes was found. Two exceptions were pet14.01 and pet09.02, for which the frequency of the “dune” orientation was relatively low.

### Genetic maps

After SNP filtering, a total of 117 individuals and 9,926 markers from non-dune mapping population, and 128 individuals and 11,748 markers from dune mapping population, entered the map construction process. The final map for non-dune ecotype is made up of 2,559 markers at 801 unique positions with 98.5% of the map having a marker at least every 10 cM and 89.7% having a marker every 5 cM. Similarly, the map for the dune ecotype is made up of 3,077 markers at 571 unique positions with 96.8% of the map having a marker every 10 cM and 87.4% of it having a marker every 5 cM. Both of the final genetic maps correspond well with the expected 17 chromosomes and translocations found previously between *H. petiolaris* and *H. annuus* (Burke et al., 2004; Ostevik et al., 2019). The LGs are longer than the map reported by Burke et al. (2004), which is probably due to greater coverage of the genome. However, we cannot rule out the possible that a low level of genotyping error from our GBS mapping approach may have contributed as well, although note that our maps are comparable in length with maps for the two subspecies *of H. petiolaris* recently reported by Ostevik et al. (2019). Two LGs in the dune map were unexpectedly short (D_LG2 and D_LG5; Supporting Information Figure S4, S5) due to few markers from the middle of the corresponding reference chromosomes, which caused the LGs to split after stringent filtering. After reconstruction with less stringent parameters, LG5s in both maps were of similar size and had enough coverage for map comparisons.

In map comparisons of the putative inversions, pet05.01 exhibited the expected pattern of reverse marker orders between the two maps. In the map for non-dune ecotype, markers were largely syntenic with the reference genome, while in the map for dune ecotype, there was a continuous block of markers with inverted order relative to the reference (Figure 4a). However, for pet07.01, pet09.01, pet09.02 and pet14.01, marker orders did not differ between the maps. But, for pet09.01, the many markers that mapped to this region formed tight clusters in both maps, indicating very low recombination in the wild non-dune and dune plants used to make these maps (Figure 4, Supporting Information Figures S6). This implies that both plants are heterozygous for the pet09.01 inversion, which would account for the recombination suppression observed. A similar pattern of reduced recombination was seen for pet11.01 and pet17.01 in the non-dune maps, but not in the map made from dune plant, in which markers from the region were in reverse order compared to the reference. Interestingly, markers with reverse order only covered part of the region for pet11.01, which implies the presence of an adjacent low recombination region or sequential inversions (Supporting Information Figures S6).

**FIGURE 4.**
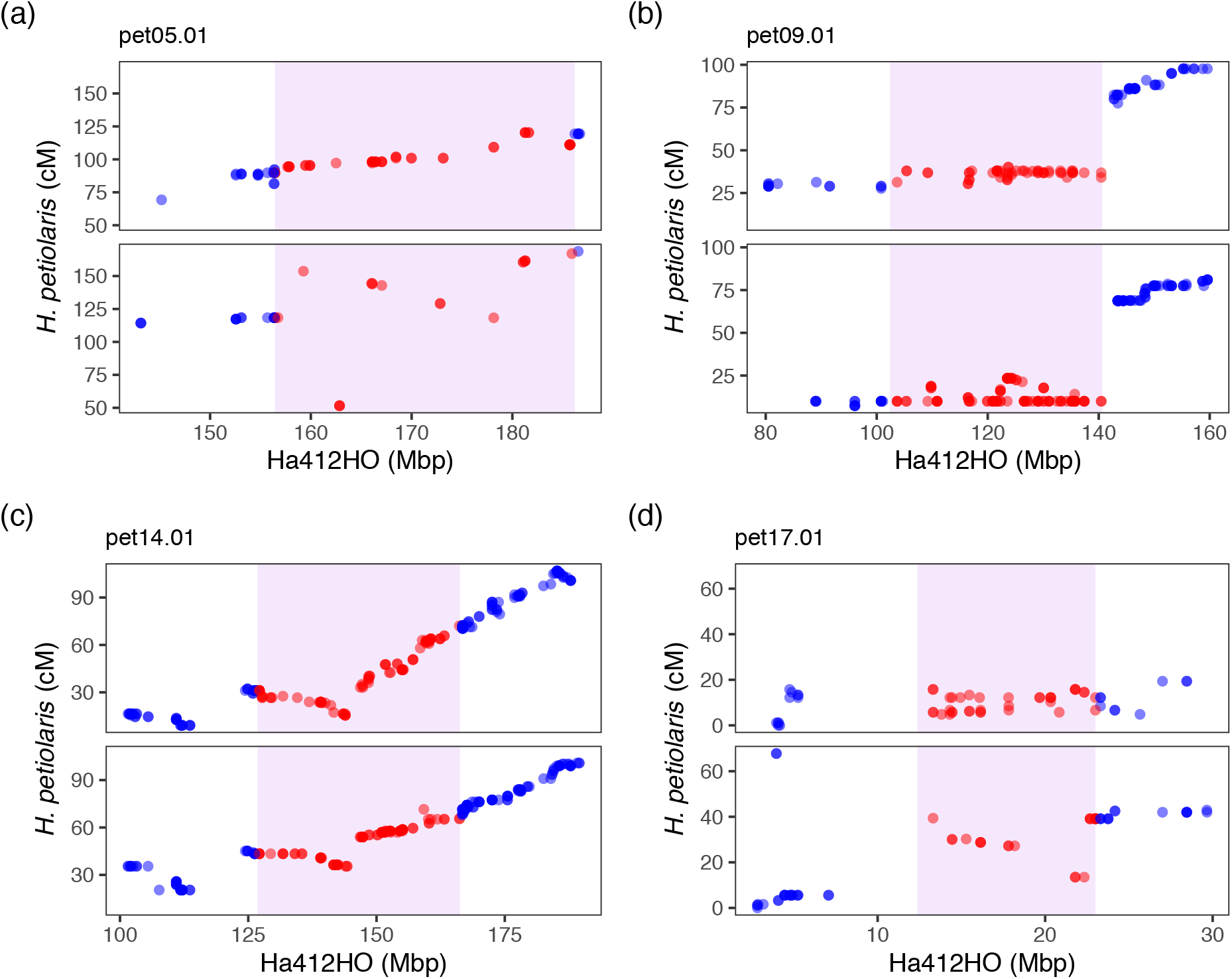
Genetic map comparisons for (a) pet05.01, (b) pet09.01, (c) pet11.01 and (d) pet17.01. Maps for non-dune (top panels) and dune (bottom panels) are plotted relative to the HA412HOv2 reference genome. Regions identified by lostruct and the markers that fall within them are highlighted in violet. Different patterns of marker orders are shown: reverse ordering between ecotypes for pet05.01 (a); recombination suppression in both maps for pet09.01 (b); similar forward ordering for pet14.01 (c); as well as recombination suppression in one map and reverse ordering in another for pet17.01(d)

Genotyping of the inversions in the parental plants using GBS confirms our interpretations. The dune and non-dune parental plants were homozygous for different haplotypes of pet05.01 and heterozygous for both haplotypes at pet09.01. For pet11.01 and pet17.01, the dune plant was homozygous while the non-dune was heterozygous for the inversion, which explains the clustering of markers in the non-dune maps.

### Genome-environment association analysis

After stringent filtration, 8,383 SNPs were retained for GEA analysis. In GEA, we found several large genomic regions with consistently high BF values, most of which overlapped nearly perfectly with the putative inversions. When treated as single loci, the putative inversions typically exhibited associations that were similar in strength to the peaks seen for the genome-wide SNPs (Figure 5, Supporting Information Figures S7, S8).

**FIGURE 5.**
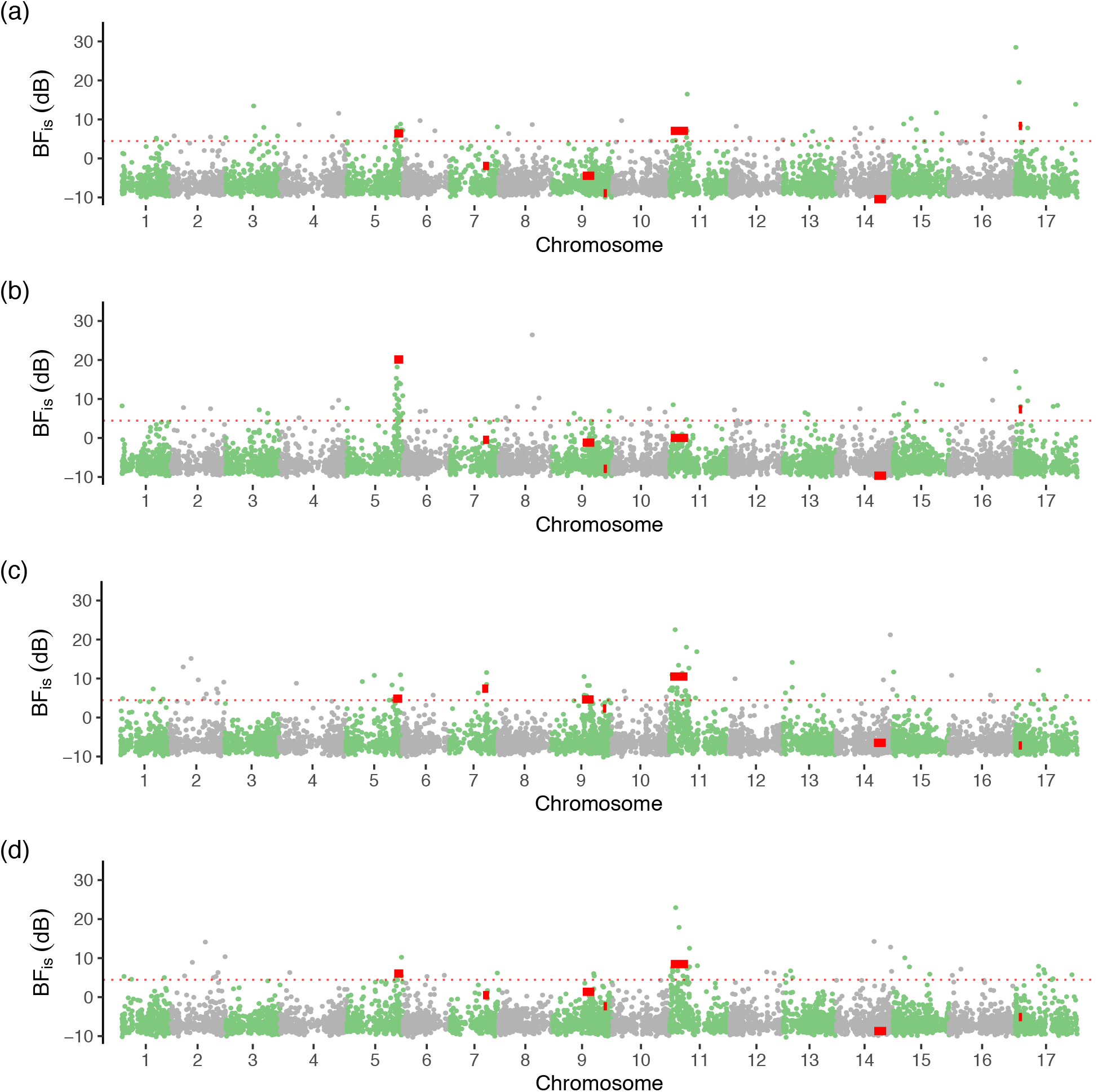
Genome-environment association for (a) % grasses, (b) coverage PC1, (c) soil NO_3_ nitrogen and (d) soil PC2. Bayes factors (BF_is_, in deciban unit) was estimated using the importance sampling estimator approach in BayPass. SNPs on different reference chromosomes are represented in alternate colors. The locations and BF_is_ values of 7 putative inversions are indicated by red solid bars. Red horizontal dashed lines represent 1% significance thresholds computed from simulated samples

The BF thresholds computed with POD ranged from 1.42 to 5.46 decibans (dB) depending on environmental variables. Several putative inversions displayed significant associations with environmental variables. The strongest signal of association was found for variables describing vegetation cover (e.g., % forbs, % grasses, and % debris), with the most striking one being pet05.01 with PC1 of coverage variables (Table 2). pet17.01 was also found to be associated with coverage variables, especially total cover. For soil characteristics, the strongest association was found for pet11.01 with NO_3_ nitrogen. pet11.01 also displayed a significant association with PC2 of the soil variables but it was not as strong. Pet07.01 displayed significant associations with a number of soil variables but not with any of the three soil PCs. In contrast, pet05.01 was marginally associated with soil PC2, but not with any of the individual soil variables. Interestingly, % grasses is strongly associated with both pet05.01 and pet11.01, whereas % forbs is only associated with the former. This pattern might be related to nitrogen availability, since nitrogen (also associated with pet11.01) is often limiting for grasses, but not for legumes, which are the most frequent forbs on the dunes.

**TABLE 2.** Bayes factors of genome-environment association analyses with coverage and soil data for putative inversions treated as single loci. Asterisks indicate Bayes factors above significance thresholds computed with simulated POD samples

| Variable | pet05.01 | pet07.01 | pet09.01 | pet09.02 | pet11.01 | pet14.01 | pet17.01 |
| --- | --- | --- | --- | --- | --- | --- | --- |
| Grass | 6.423* | -1.917 | -4.443 | -8.937 | 7.064* | -10.441 | 8.333* |
| Forb | 13.942* | -5.516 | -3.309 | -8.141 | -7.581 | -7.601 | -4.286 |
| Debris | 14.286* | -4.332 | -4.853 | -9.645 | -4.219 | -10.423 | 0.572 |
| Cover | 17.758* | -0.64 | -2.146 | -8.735 | 1.473 | -9.978 | 11.316* |
| Cover PC1 | 20.125* | -0.471 | -1.188 | -7.905 | 0.016 | -9.672 | 7.337* |
| Total N | 4.216 | 6.822* | 5.223* | 2.62 | 9.635* | -6.687 | -7.267 |
| NO3-N | 4.85 | 7.413* | 4.64 | 2.319 | 10.517* | -6.482 | -7.171 |
| Ca | 0.13 | 8.1* | -5.029 | -6.64 | -4.792 | -6.115 | -7.023 |
| P | -1.725 | 4.957* | -3.374 | -2.521 | 0.819 | -8.546 | -8.808 |
| S | -3.023 | 4.078* | -4.504 | -5.88 | 4.324* | -9.177 | -8.886 |
| Soil PC2 | 6.048* | 0.5 | 1.364 | -2.347 | 8.469* | -8.699 | -5.121 |

Only the environmental variables with a significant association with at least one putative inversion are shown.

## DISCUSSION

Genomic islands of differentiation often arise between diverging populations connected by gene flow (Feder & Nosil, 2009). While regions with higher than average differentiation can be created by divergence hitchhiking (Via, 2012), such regions are unlikely to be large or to have the sharp boundaries often reported for islands of divergence. Inversions represent a more likely explanation for large and discrete islands since recombination is reduced across the entire inverted region. Also, unlike other recombination modifiers, inversions reduce recombination between haplotypes, but not within them, which facilitates adaptive divergence. Theory indicates that inversions will be favored if they prevent recombination between locally adapted alleles when challenged by migration of non-adapted alleles (Kirkpatrick and Barton 2006). Inversions can also facilitate speciation by preventing recombination between locally adapted alleles and those contributing to assortative mating (Ortiz-Barrientos, Engelstädter, & Rieseberg, 2016).

Despite the clear importance of inversions in adaptation and speciation, it remains difficult to identify and genotype them, especially in non-model systems. Using a population genomic approach with RAD sequencing data, we detected seven putative chromosomal inversions that separate dune and non-dune *H. petiolaris* in GSD, which we validated by a combination of population genetic and comparative genetic mapping approaches. Also, we demonstrated that inversions account for the genomic islands of high divergence between the ecotypes and contribute to ecological divergence in this system.

### Identification of inversions

Employing the methods implemented in lostruct, which makes use of the effect that inversions have on population structure, we found clusters of windows with outlier MDS values, i.e. genomic regions with extreme population structure compared to the rest of the genome, and we provided multiple lines of evidence showing that the majority of these signals are left by inversions.

There are other processes that can generate a pattern of contiguous outlier MDS, such as selection coupled with gene flow, low recombination, or introgression. Linked selection can generate heterogeneous population structure across the genome (Li & Ralph, 2019), especially when selection is strong and acts in the face of gene flow, and may also generate long LD blocks. However, the regions that we identified are typically > 10Mb. It is unlikely that the effect of selection would span a region of several to tens of Mbp on the genome in the absence of structural variation. Moreover, such regions under selection are expected to generate a continuous pattern of population structure in a PCA as opposed to the three discrete clusters with higher heterozygosity in the middle cluster reported here. Lastly, the finding of high LD across putative inversions when tested across all samples, but not within putative homozygous groups, distinguishes inverted regions from other regions of reduced recombination (e.g. centromeres), because other mechanisms of recombination suppression are expected to restrict recombination in all groups of individuals. Other small, blurred-edged regions of low recombination were also found in our LD analysis (e.g., on chromosome 8 from 85-100 Mbp and chromosome 17 from 185-205 Mbp; Supporting Information Figure S2), but they displayed symmetric patterns of LD in different sample sets and were often associated with low sequence coverage, suggestive of centromeres or other heterochromatic regions. Introgression from another species can also form two distinct haplotype blocks and generate patterns similar to those of an inversion. However, gene flow and recombination will erode such patterns unless the introgression is recent.

Using genetic maps, we were able to validate one of the inversions (pet05.01) identified with population genetic data and provide additional support for three more based on suppressed recombination in putative inversion heterozygotes (pet09.01, pet11.01 and pet17.01). However, because the wild parents might have the same haplotype for pet07.01, pet09.02 and pet14.01, we were unable to corroborate them. This demonstrates one of the weaknesses of the genetic mapping approach – mapping will only detect a subset of segregating inversions. In contrast, approaches based on population genetic data provide a fine-grained and comprehensive way to search for potential inversions, and our methods appear to be robust.

Using RAD sequence data, we detected six structural variants identified from WGS data (Todesco et al., 2019) and one additional new putative inversion (pet09.02). We demonstrated that reduced representation sequencing data have the same power to detect inversions with SNP densities as low as 12 per Mbp. Moreover, with more extensive sampling across the habitat transition than that used by Todesco et al., we were able to better estimate population allele frequencies, as well as genetic divergence between ecotypes. We further demonstrated that these inversions are enriched in dune environment and that they correspond closely to genomic islands of differentiation at GSD (see below).

However, there are limitations to our approach for detecting inversions. First, while a population genomic approach such as that employed here can provide initial clues regarding the existence of chromosomal inversions, additional independent evidence, such as comparative genetic mapping in this study or Hi-C sequencing analysis by Todesco et al. (2019), is needed to confirm the inversions for further investigation. Second, pinpointing the positions of breakpoints is not feasible given the low density of RAD markers. This can be challenging even with high-depth whole genome sequencing because of the abundance (typically) of repetitive sequences near breakpoints (Tang et al., 2015). Third, the limited genomic coverage of RAD sequence data, together with the dependence on deviations in population structure, biases detection towards large inversions with high sequence divergence. Therefore, it is not suitable for estimating the rate of origin and size distribution of chromosomal variants. However, it offers a convenient way to explore the evolutionary role of inversions because large and highly divergent inversions are also the ones that are most likely to play an important role in local adaptation and speciation. Lastly, we expect that the approach we described here could be further improved by better tuning of window size and outlier thresholds to match population sizes and SNP densities. Despite these limitations, our workflow provides a feasible and economical way of examining inversion frequencies and their evolutionary role in natural populations.

### Inversions contribute to adaptive divergence

Previous work identified several large regions of differentiation that displayed signatures of divergent adaptation between dune and non-dune ecotypes in this system (Andrew & Rieseberg, 2013). Our analyses showed that recombination is suppressed in these highly divergent genomic regions due to chromosomal inversions. Increasing evidence suggests that such islands of differentiation may be prevalent in incipient species (Turner et al., 2005; Michel et al., 2010), and inversions have been shown to play an important role in maintaining ecological and genetic divergence in the face of gene flow (Rieseberg, 2001; Noor et al., 2001; Feder et al., 2003; Lowry & Willis, 2010). Our findings add to the growing body of case studies on how structural chromosomal changes interact with local adaptation and gene flow to shape the genomic landscape of divergence in early stages of speciation.

Analyses of inversion haplotype frequencies based on genotypes inferred from k-means showed that all of the inversions are significantly enriched on the dunes (Table 1), suggesting that they may be under selection, although for some inversions “non-dune” alleles are often found as heterozygotes on the dunes. This could be due to differences in the kinds and strength of selection on the inversions, but could also result from our sampling scheme. The individuals used in the study were collected as seeds from mature plants, and thus reflected post-mating population frequencies rather than that of living plants. If the inversions contribute to seedling survival in dunes, then we likely are under-estimating frequency differences between ecotypes. This is not implausible given that selection against immigrants is known to contribute strongly to reproductive isolation in this system (Ostevik et al., 2016).

Additional evidence that the inversions contribute to local adaptation comes from the observation that four of the inversions (pet05.01, pet09.01, pet11.01 and pet14.01) co-localize with seed size QTLs identified in other work (Ostevik, 2016; Todesco et al., 2019). Large seeds help plants survive burial in actively moving sand dunes (Donovan, Rosenthal, Sanchez-Velenosi, Rieseberg, & Ludwig, 2010; Ostevik et al., 2016), and seed size is the most divergent phenotypic trait between the ecotypes. These observations are further reinforced by the strong association of pet05.01 with vegetation cover, which is negatively correlated with dune stability. Among the inversion haplotypes associated with increased seed size, pet14.01 was in relatively low frequency. However, this inversion underlies ecotype differentiation in another dune ecotype of *H. petiolaris* (Todesco et al., 2019). Possibly, pet14.01 was only recently introduced to GSD, so it will be interesting to monitor its frequency over the next 1-2 decades. Several inversions were also found to be associated with soil variables in our GEA analyses. Sand dunes are characterized by low nutrient availability, and a QTL for leaf N content maps to inversion pet11.01 (Todesco et al., 2019), which we have shown to be associated with soil N in this study, suggestive of a role in tolerance to low nutrients. Future mapping studies of related physiological traits would help reveal the mechanistic basis by which inversions, especially pet11.01, aid adaptation to low nutrient soils.

In the study by Todesco et al. (2019), multiple traits and soil characteristics were constantly found associated with the same inversions in *H. petiolaris*. These signals could be caused by the low number of samples in the dunes and the resulting selection-driven linkage of the inversions among those samples. With denser sampling across the landscape, we were able to break the linkage of dune inversions and disentangle the effects in GEA. We show that various sets of inversions are responsible for different aspects of dune adaptation in this system.

The observation that inversions are associated with different traits and environmental factors in the dune habitat implies that the inversions are likely favored because they maintain combinations of locally advantageous alleles despite ongoing gene flow with non-adapted populations (Kirkpatrick & Barton, 2006). Models of parapatric and sympatric speciation have emphasized the importance of linkage between genes underlying local adaptation and those involved in reproductive isolation (Ortiz-Barrientos et al., 2016; Servedio, 2009; Noor et al., 2001). A key assortative mating barrier between the ecotypes is conspecific pollen precedence (Ostevik et al. 2016). Thus, a hypothesis going forward is that loci causing conspecific pollen precedence will also be located within one or more of these inversions

## CONCLUSION

Using RAD sequencing data and a population genomic approach, we were able to detect multiple inversions *de novo* at low cost, determine their frequencies in natural populations, and assess their role in adaptation through GEA analyses. Localized heterogeneity of population structure caused by inversions has been detected in other systems using whole genome sequencing data (Li & Ralph, 2019). We show that inversions can also be detected with reduced representation sequencing data with low SNP densities. Given the ever-expanding population sequencing data available for non-model systems, we anticipate an explosion of inversion reports across the plant and animal kingdoms, especially in systems where divergence appears to have occurred in the face of gene flow.

## ACKNOWLEDGEMENTS

We thank Qin Li for help in map plotting with R, and Shaghayegh Soudi for suggestions on BayPass. This work was supported by a China Scholarship Council scholarship (no. 201506380099) to K.H., a Killam Postdoctoral Fellowship to R.L.A., and NSERC grant (327475) to L.H.R.

## DATA ACCESSIBILITY

RAD sequencing data published previously: Dryad doi:10.5061/dryad.j2448. Environmental data published previously: Dryad doi: 10.5061/dryad.158pb518. Scripts and SNP data for genetic map construction are available upon request and will be set to GitHub and Dryad before publication, respectively.

## AUTHOR CONTRIBUTIONS

K.H. and L.H.R. conceived the study; R.L.A. contributed genetic and environmental data; K.H. performed all the analyses; G.L.O. helped with the local structure analysis; K.L.O contributed to genetic map construction and synteny analysis; K.H. and L.H.R. wrote the paper; and all authors approved the final manuscript.

