## Supporting Information Figure S1 for "Multiple chromosomal inversions contribute to adaptive divergence of a dune sunflower ecotype"

**FIGURE S1** Characterization of MDS outlier regions. (a) Genome plot of corresponding MDS values across 17 reference chromosomes. Each dot represents a window of 50 SNPs, and outlier windows are highlighted in red. (b) PCA with SNPs in the region. Three clusters identified using k-means clustering correspond to two homozygote groups (blue and red) and a heterozygote group (purple). (c) Heterozygosity for each of the groups identified in PCA. (d) LD plot for corresponding chromosome. Upper triangle with all individuals and lower triangle with only individuals homozygous for the more common orientation. SNPs were summarized and the second highest $R^2$ values were presented in 1 Mbp windows. Purple bars represent the range of the inversion

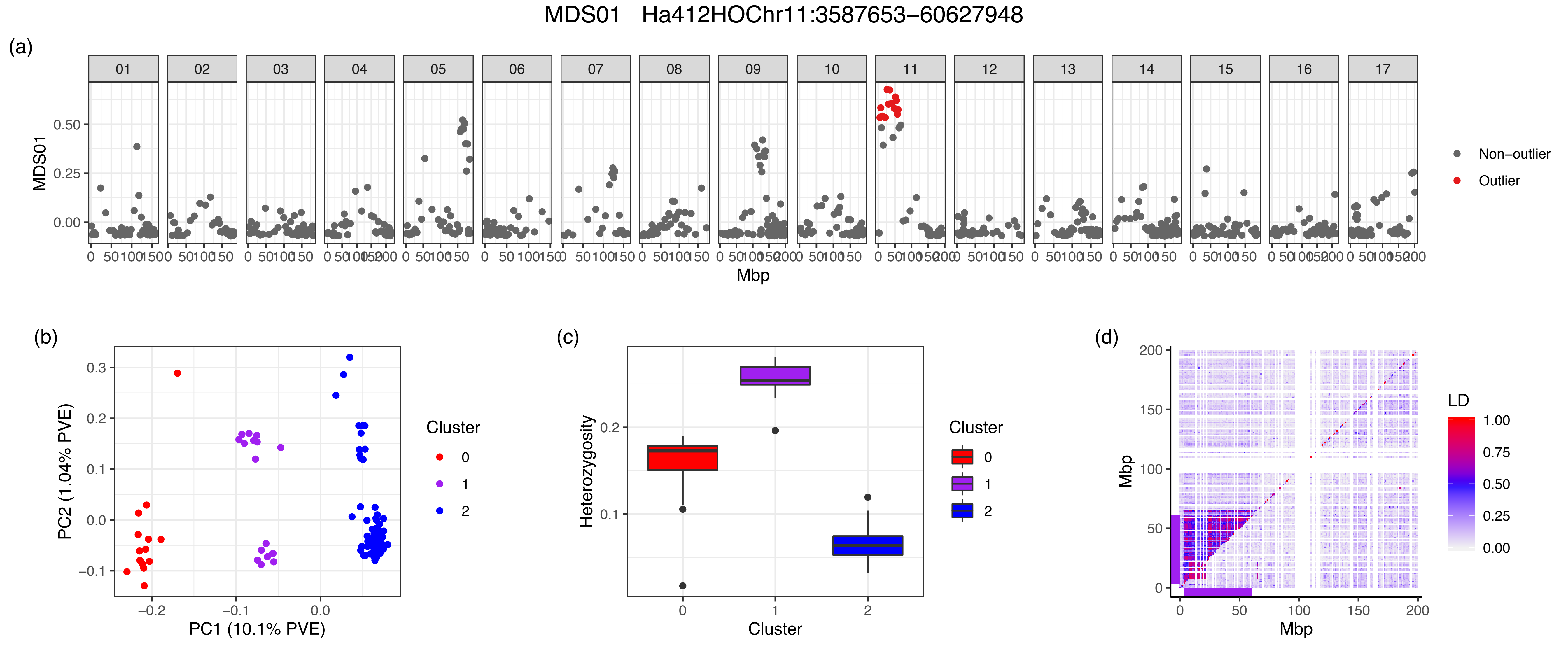
MDS01 Ha412HOChr11:3587653–60627948
(a)
MDS01
Mbp
Non–outlier
Outlier
(b)
PC2 (1.04% PVE)
PC1 (10.1% PVE)
Cluster
0
1
2
(c)
Heterozygosity
Cluster
(d)
Mbp
LD
1.00
0.75
0.50
0.25
0.00

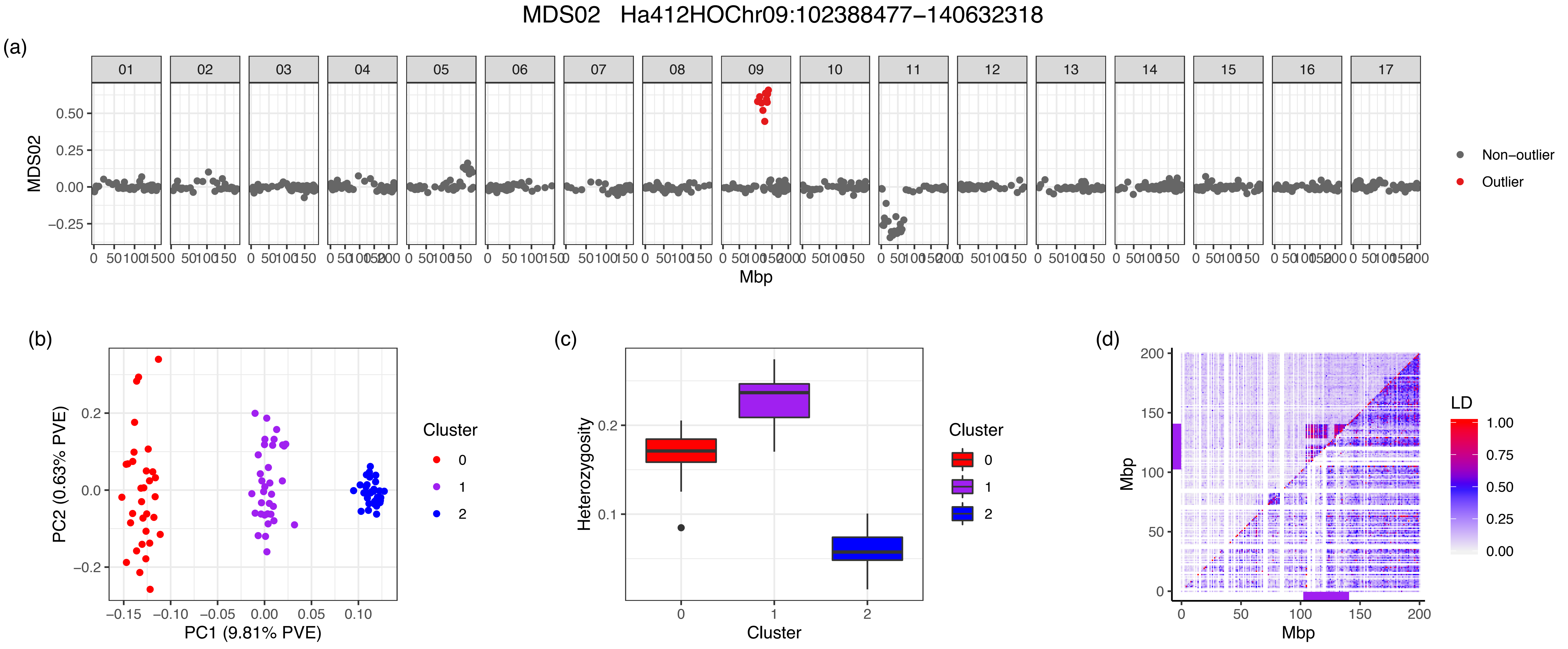
MDS02 Ha412HOChr09:102388477−140632318
(a)
MDS02
0.50
0.25
0.00
−0.25
Mbp
Non−outlier
Outlier
(b)
PC2 (0.63% PVE)
PC1 (9.81% PVE)
−0.15
−0.10
−0.05
0.00
0.05
0.10
0.2
0.0
−0.2
Cluster
0
1
2
(c)
Heterozygosity
0.2
0.1
Cluster
0
1
2
Cluster
0
1
2
(d)
Mbp
200
150
100
50
0
0
50
100
150
200
Mbp
LD
1.00
0.75
0.50
0.25
0.00

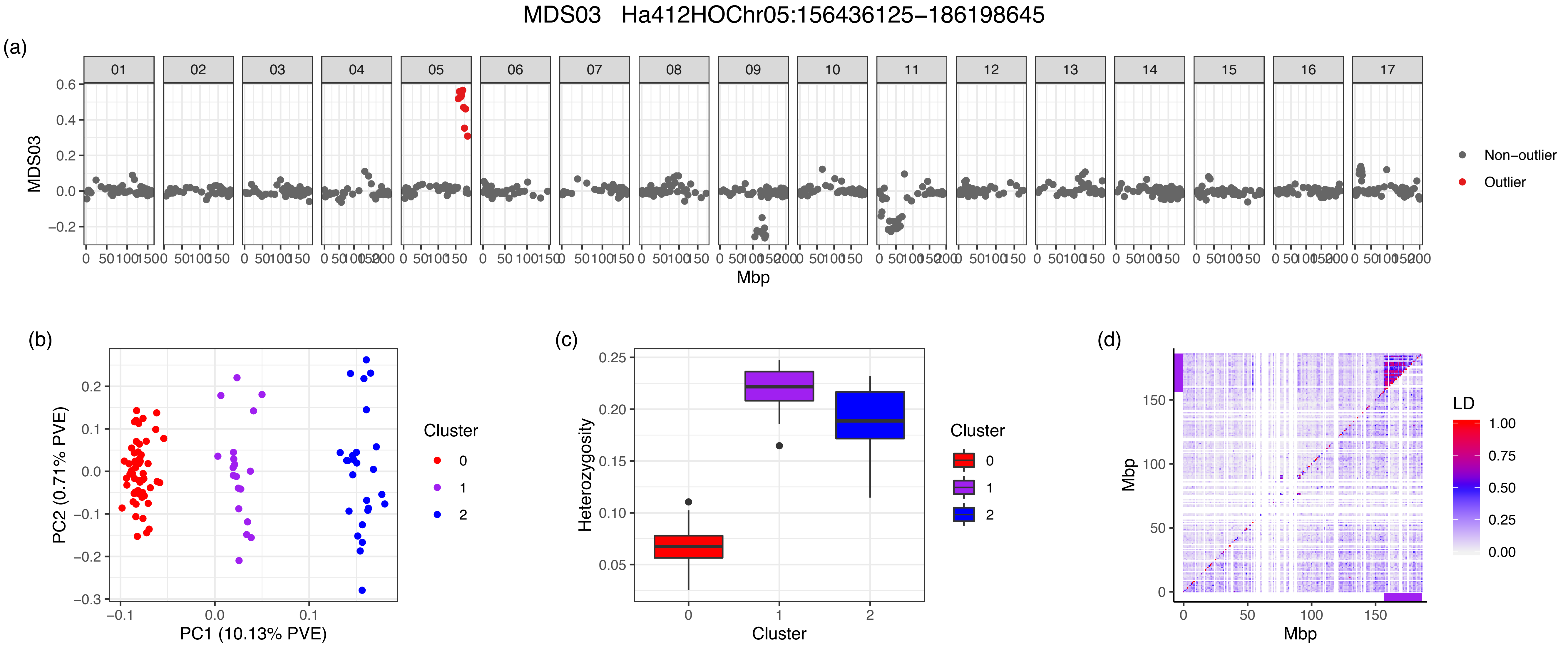
MDS03 Ha412HOChr05:156436125−186198645
(a)
MDS03
Mbp
Non−outlier
Outlier
(b)
PC1 (10.13% PVE)
PC2 (0.71% PVE)
Cluster
0
1
2
(c)
Heterozygosity
Cluster
(d)
Mbp
LD
1.00
0.75
0.50
0.25
0.00

MDS04 Ha412HOChr07:109423942−129416998

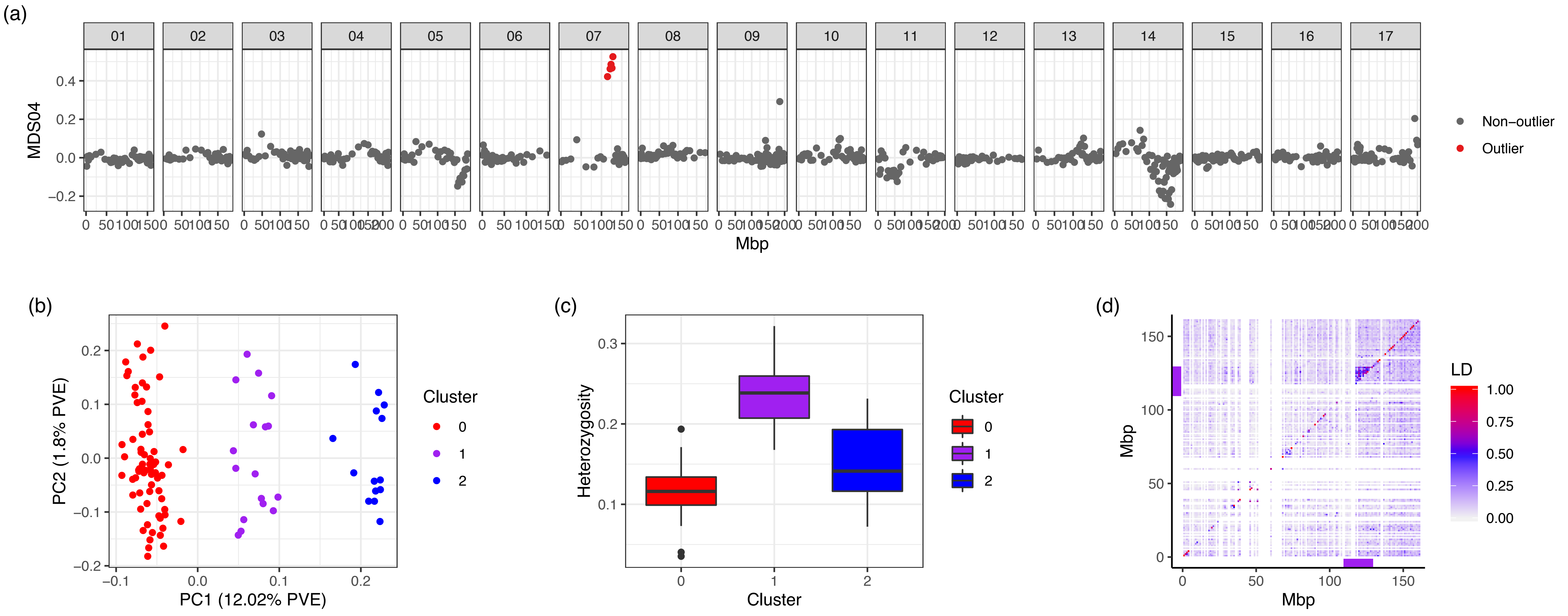

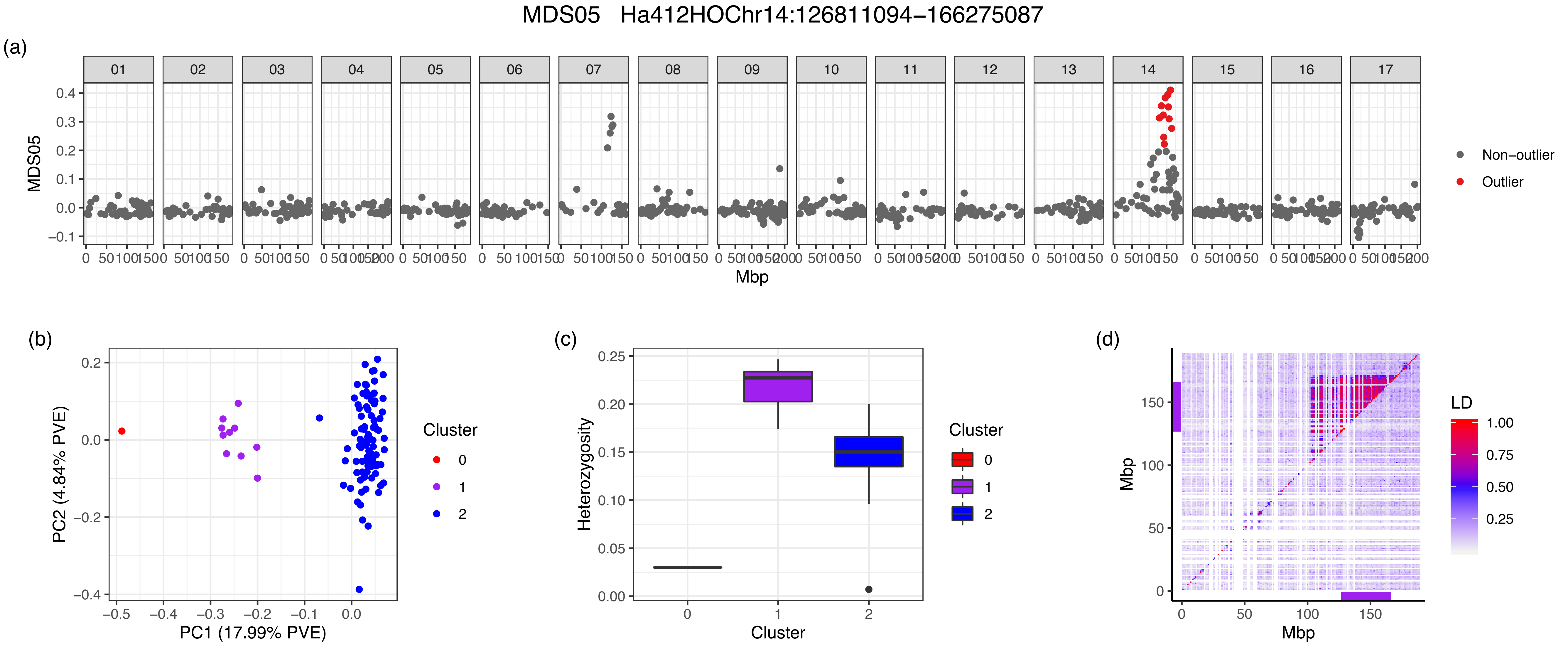

MDS05 Ha412HOChr14:126811094−166275087
(a)
MDS05
Mbp
Non−outlier
Outlier
(b)
PC1 (17.99% PVE)
PC2 (4.84% PVE)
Cluster
0
1
2
(c)
Heterozygosity
Cluster
(d)
Mbp
LD
1.00
0.75
0.50
0.25

MDS06 Ha412HOChr17:12368066–23002709

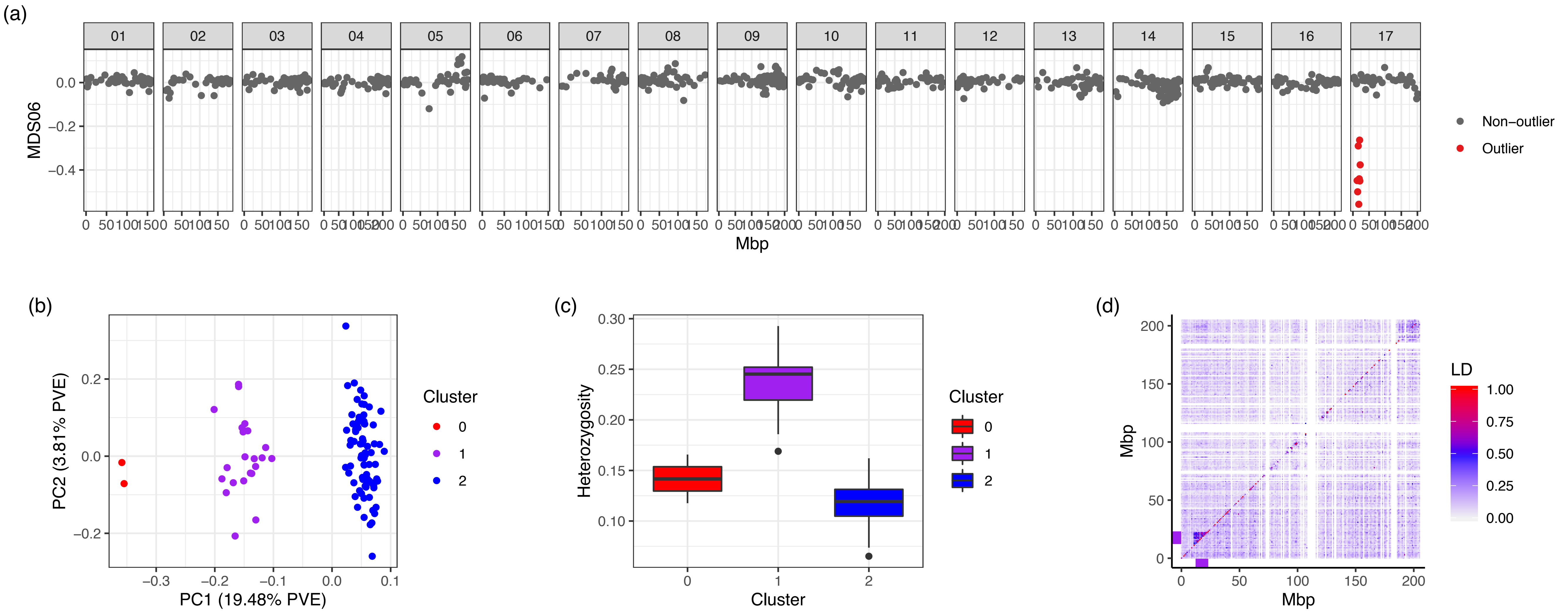

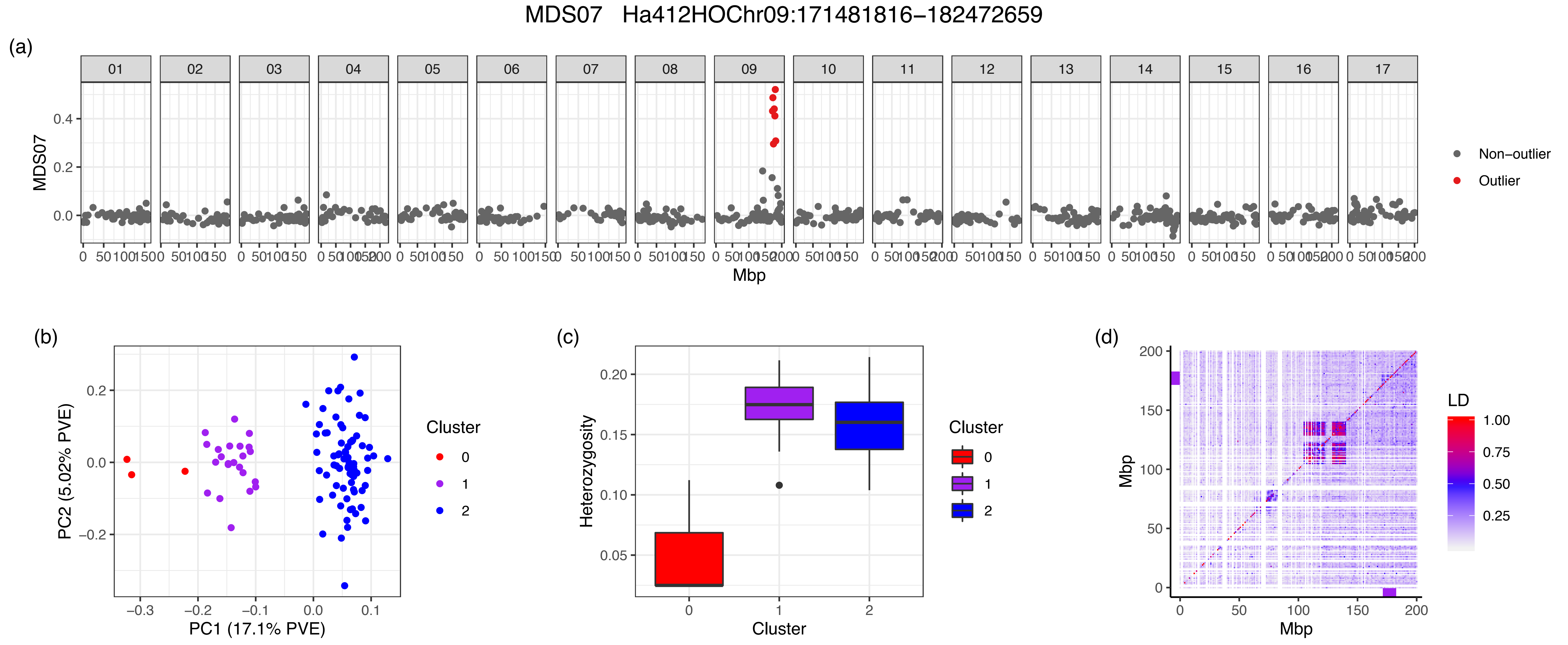
MDS07 Ha412HOChr09:171481816−182472659
(a)
MDS07
Mbp
Non−outlier
Outlier
(b)
PC2 (5.02% PVE)
PC1 (17.1% PVE)
Cluster
0
1
2
(c)
Heterozygosity
Cluster
(d)
Mbp
LD
1.00
0.75
0.50
0.25

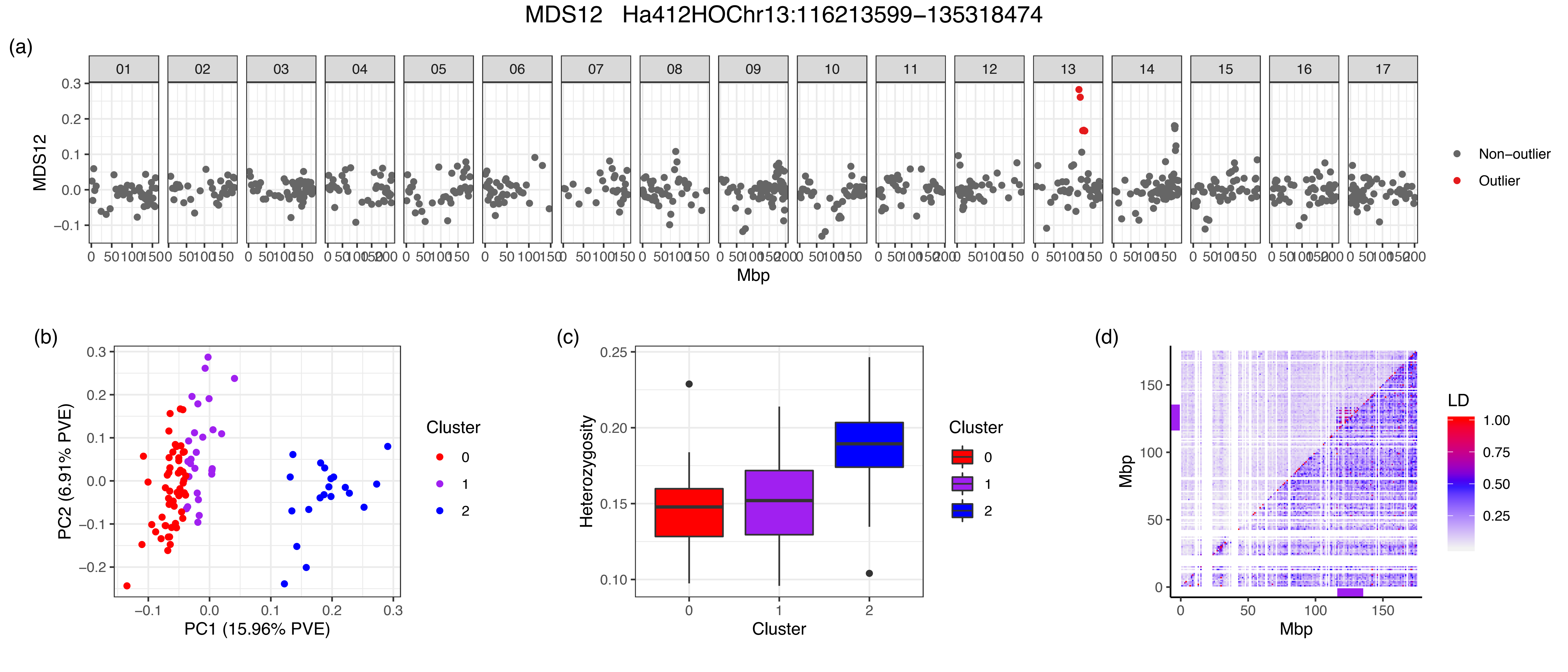
MDS12 Ha412HOChr13:116213599−135318474
(a)
01 02 03 04 05 06 07 08 09 10 11 12 13 14 15 16 17
MDS12
Mbp
Non−outlier
Outlier
(b)
PC1 (15.96% PVE)
PC2 (6.91% PVE)
Cluster
0
1
2
(c)
Heterozygosity
Cluster
0
1
2
(d)
Mbp
Mbp
LD
1.00
0.75
0.50
0.25

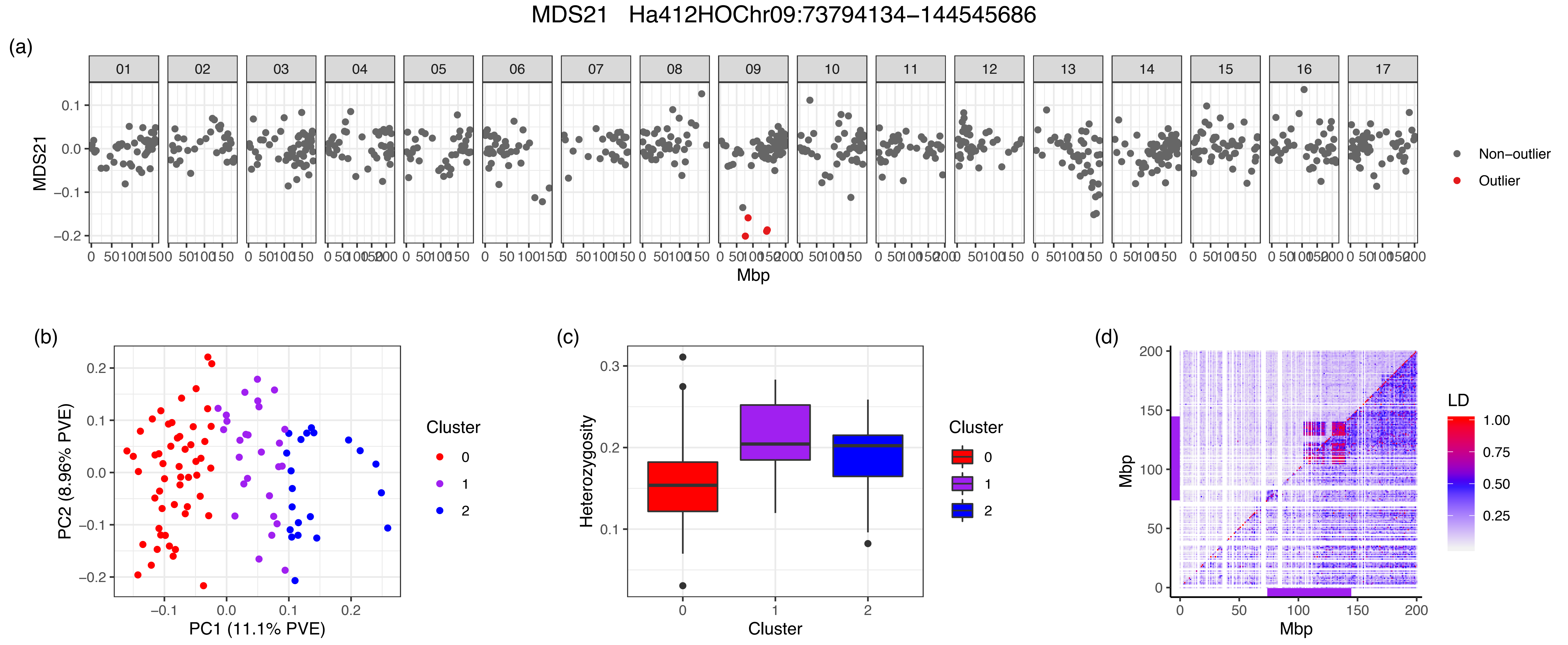
MDS21 Ha412HOChr09:73794134–144545686
(a)
MDS21
Mbp
Non–outlier
Outlier
(b)
PC1 (11.1% PVE)
PC2 (8.96% PVE)
Cluster
0
1
2
(c)
Heterozygosity
Cluster
(d)
Mbp
LD
1.00
0.75
0.50
0.25
