## Supporting Information Figure S2 for "Multiple chromosomal inversions contribute to adaptive divergence of a dune sunflower ecotype"

**FIGURE S2** LD plots for all chromosomes. Result based on all individuals is shown in upper triangle. For the chromosomes with regions of MDS outliers (named by the MDS coordinate and direction where the region was identified), calculation with only individuals homozygous for the more common orientation is shown in lower triangle. SNPs were summarized and the second highest $R^2$ values were presented in 1 Mbp windows. Purple bars represent the location of the outlier region

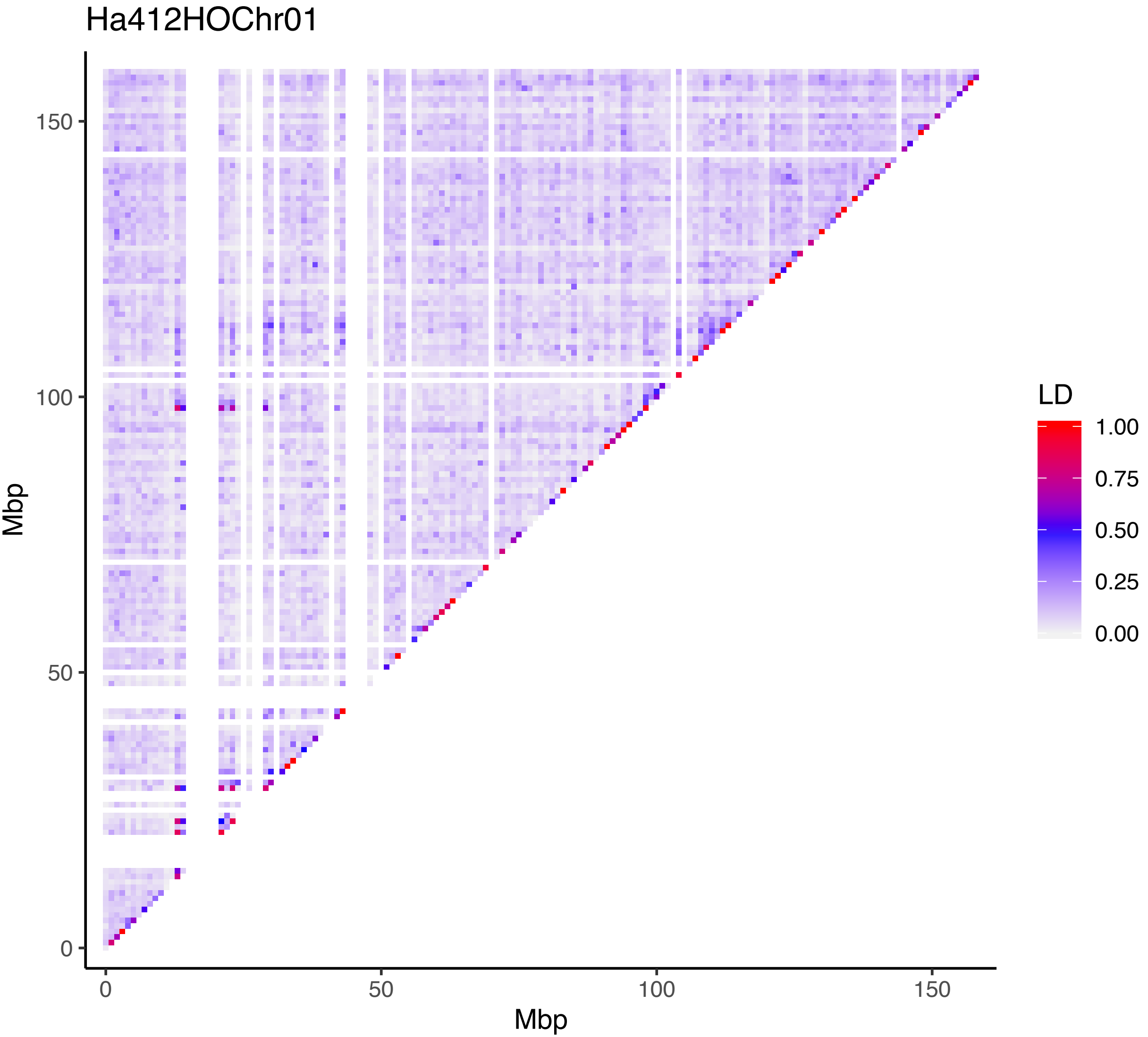

Ha412HOChr01
Mbp
150
100
50
0
0
50
100
150
Mbp
LD
1.00
0.75
0.50
0.25
0.00

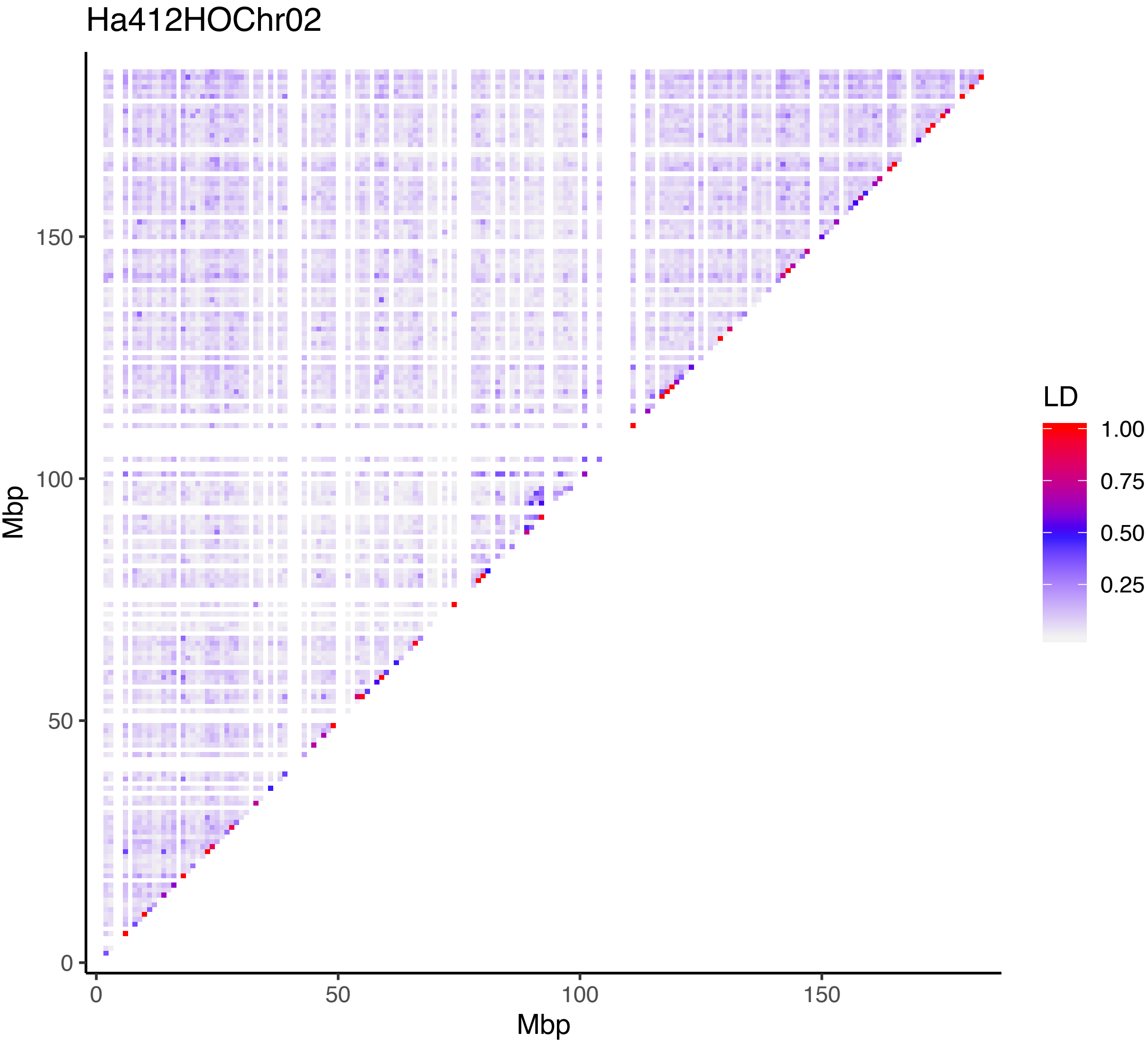
Ha412HOChr02
Mbp
150
100
50
0
0
50
100
150
Mbp
LD
1.00
0.75
0.50
0.25

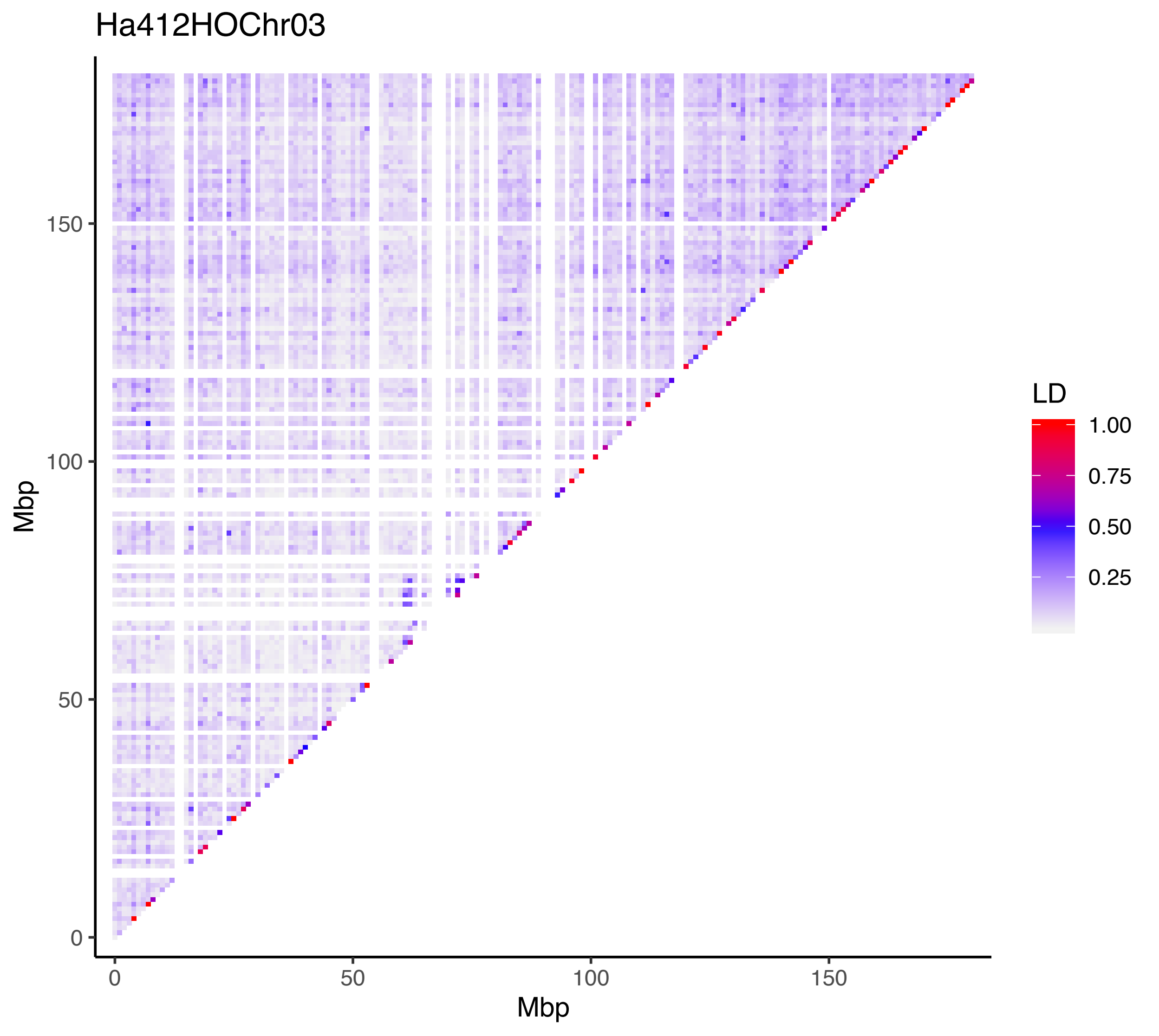
Ha412HOChr03
Mbp
Mbp
0
50
100
150
LD
1.00
0.75
0.50
0.25

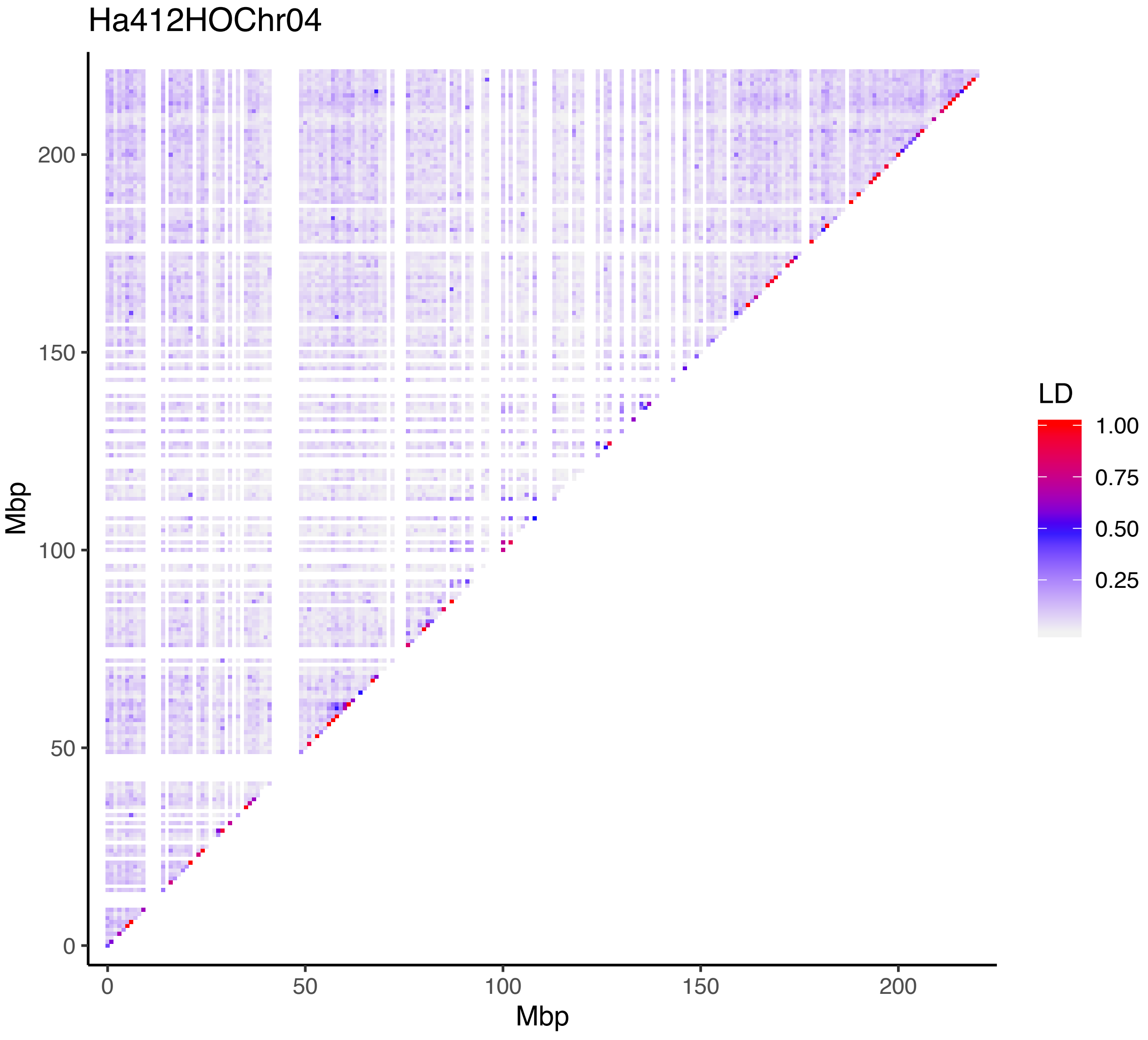
Ha412HOChr04
Mbp
0
50
100
150
200
Mbp
0
50
100
150
200
LD
1.00
0.75
0.50
0.25

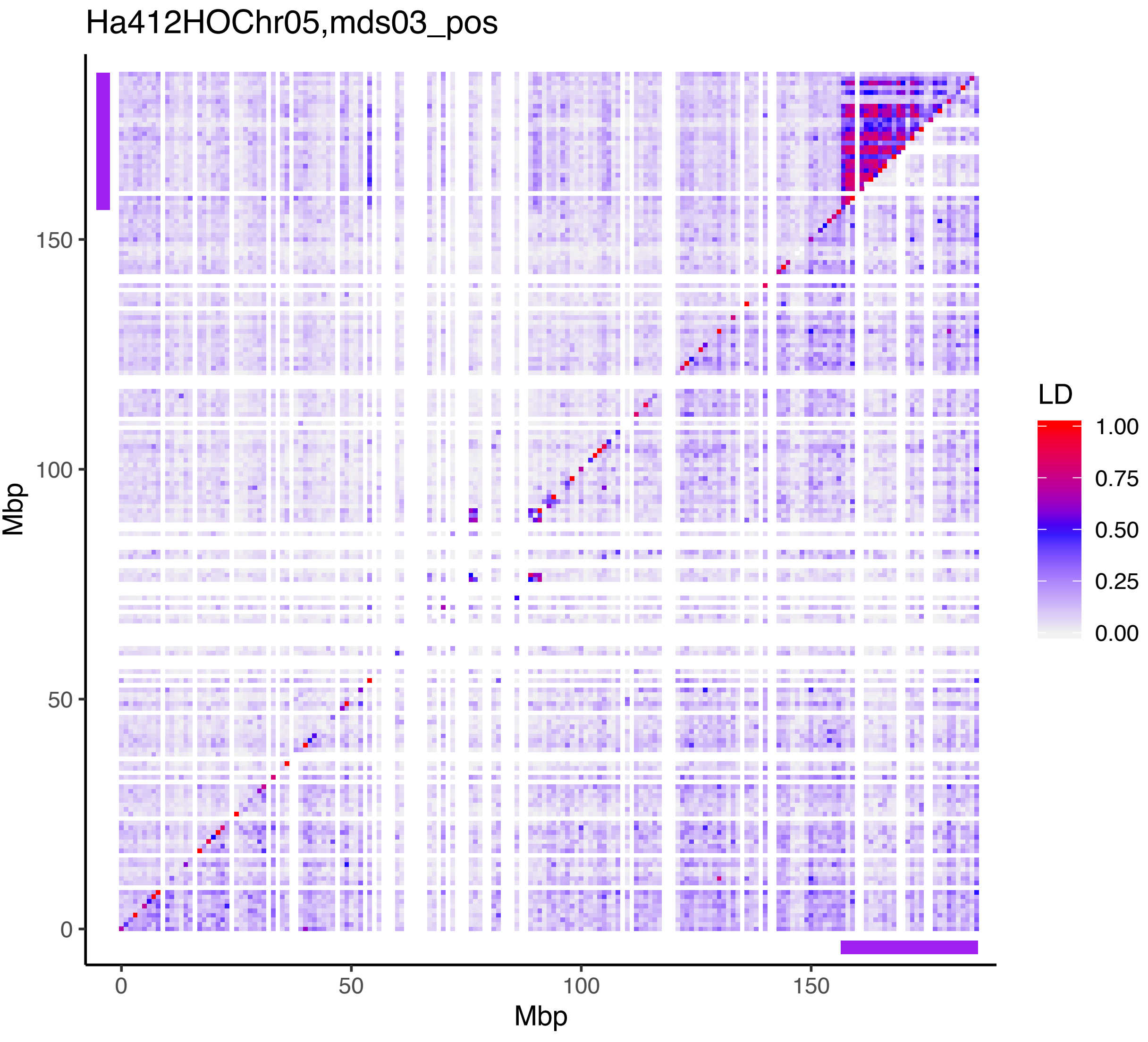

Ha412HOChr05,mds03_pos
Mbp
0
50
100
150
Mbp
0
50
100
150
LD
1.00
0.75
0.50
0.25
0.00

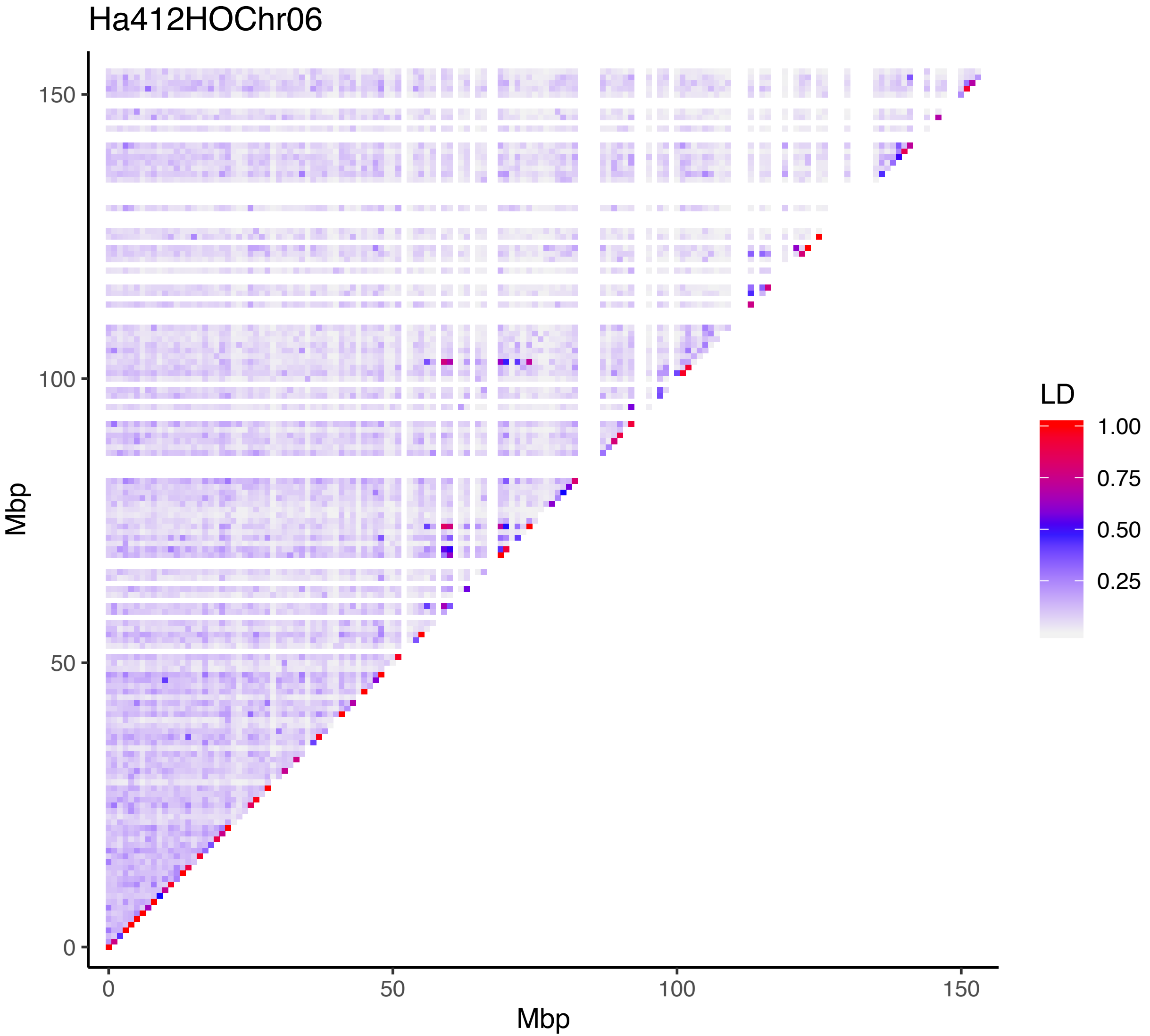

Ha412HOChr06
Mbp
150
100
50
0
0
50
100
150
Mbp
LD
1.00
0.75
0.50
0.25

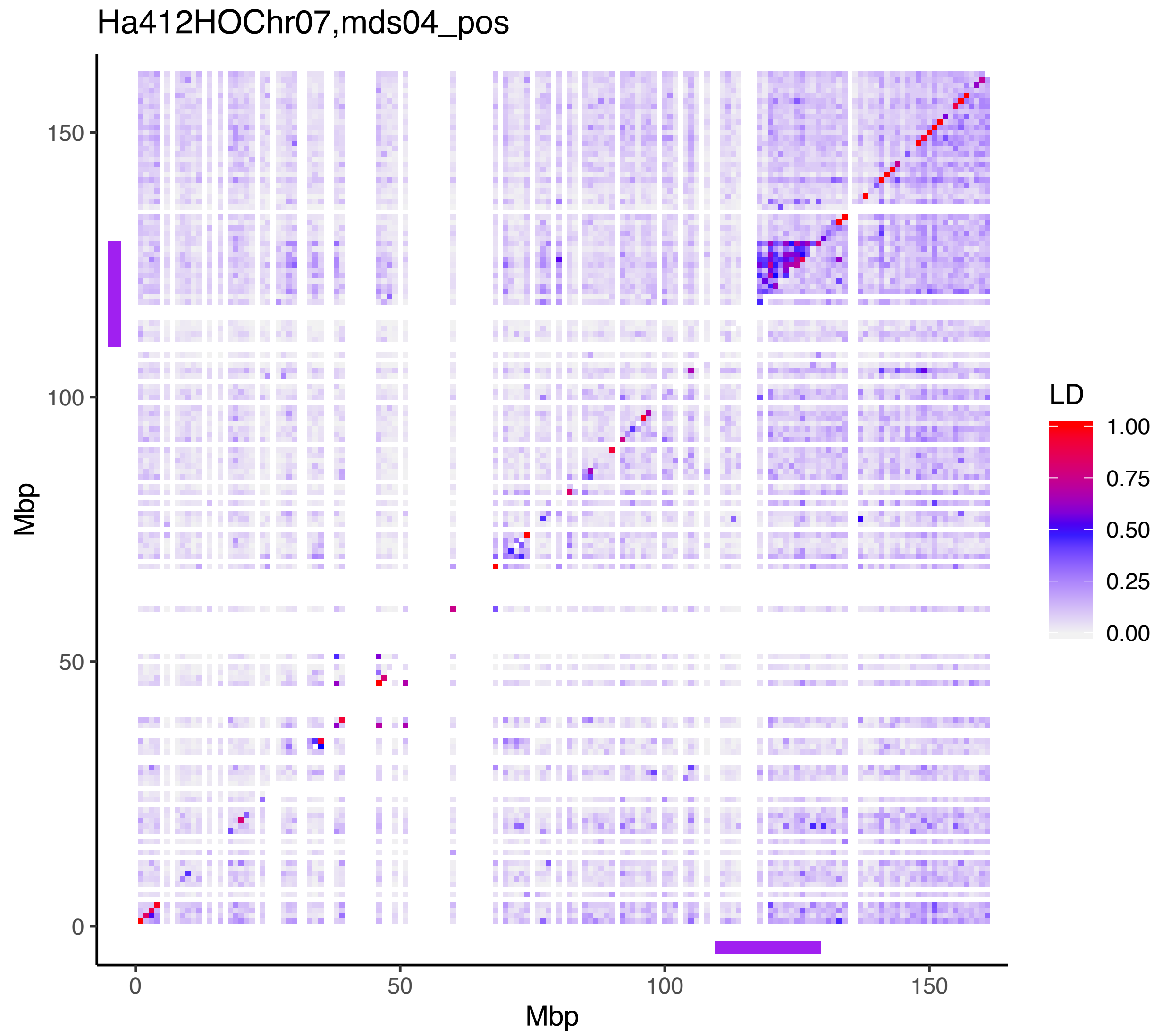

Ha412HOChr07,mds04_pos
Mbp
Mbp
LD
1.00
0.75
0.50
0.25
0.00
0
50
100
150

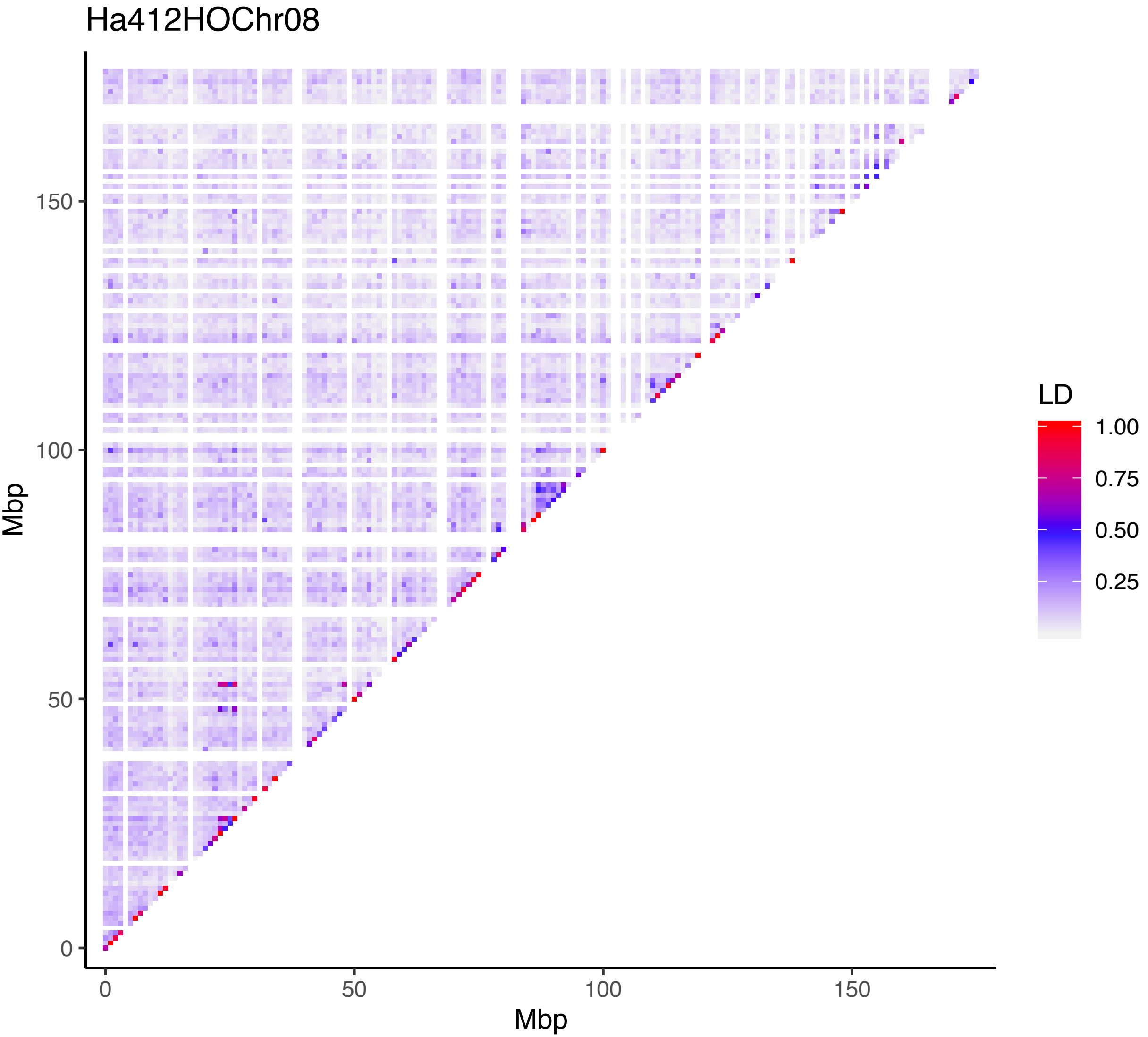
Ha412HOChr08
Mbp
150
100
50
0
0
50
100
150
Mbp
LD
1.00
0.75
0.50
0.25

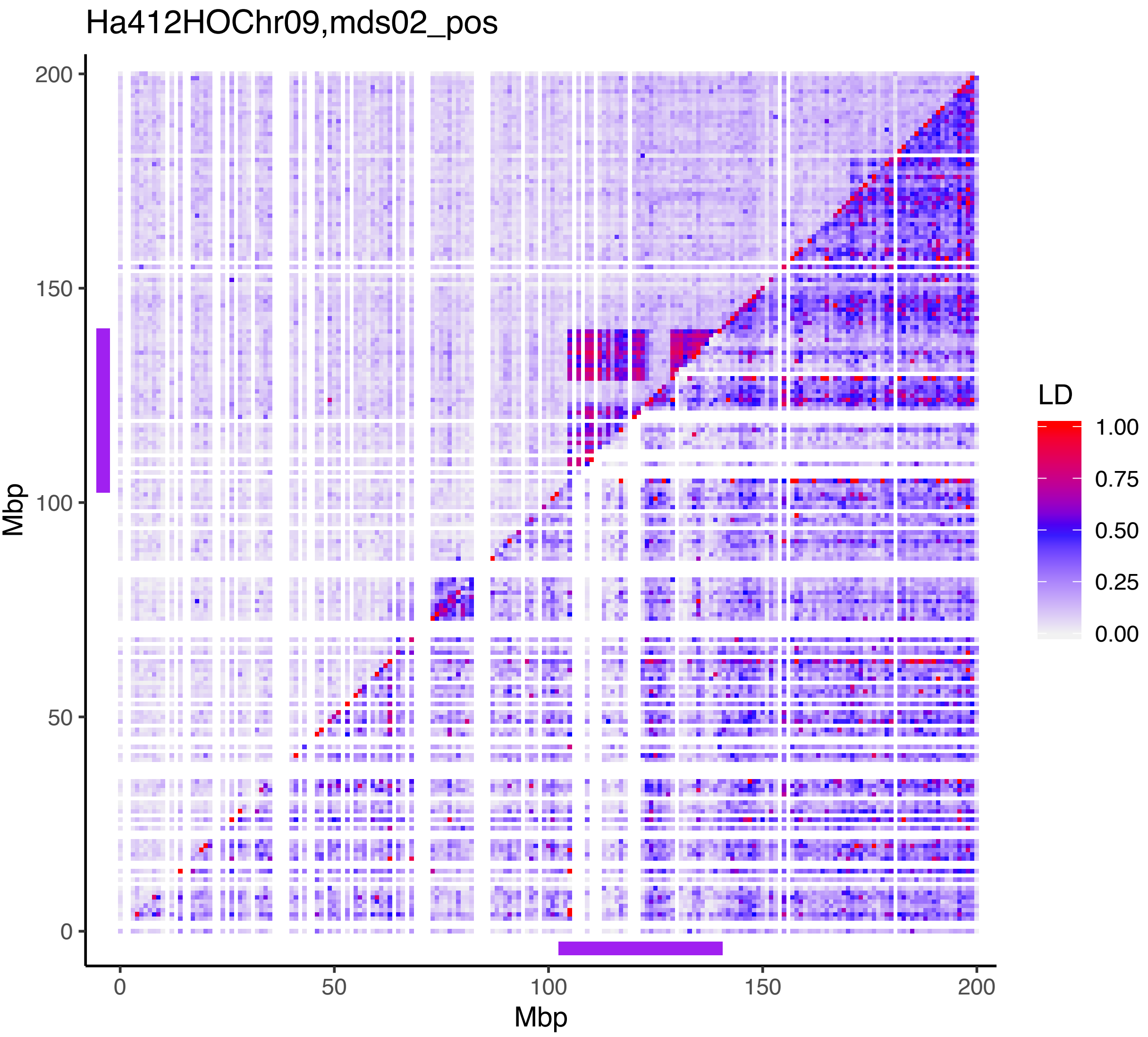

Ha412HOChr09,mds02_pos
Mbp
200
150
100
50
0
0
50
100
150
200
Mbp
LD
1.00
0.75
0.50
0.25
0.00

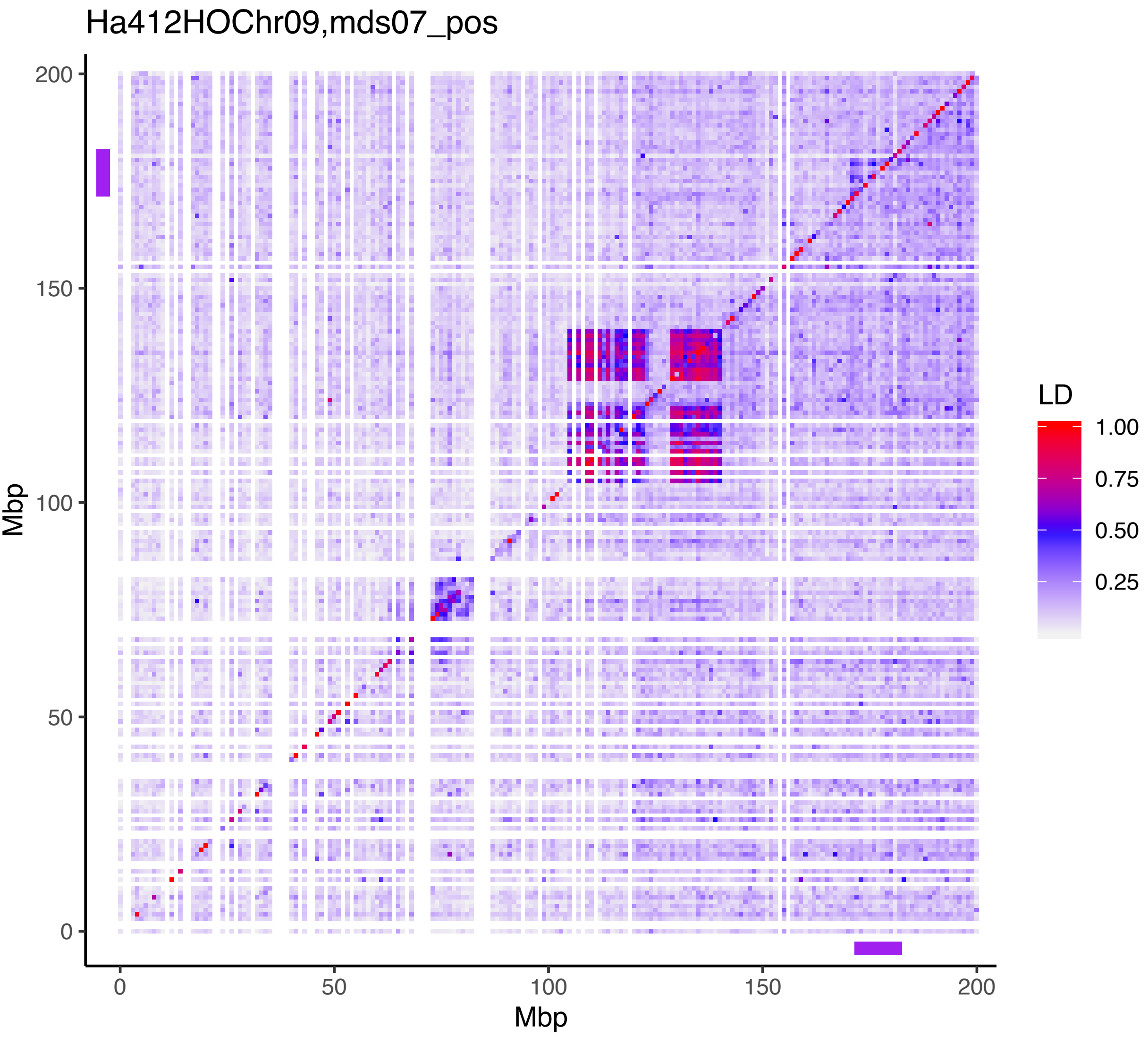

Ha412HOChr09,mds07_pos
Mbp
Mbp
LD
1.00
0.75
0.50
0.25
0
50
100
150
200

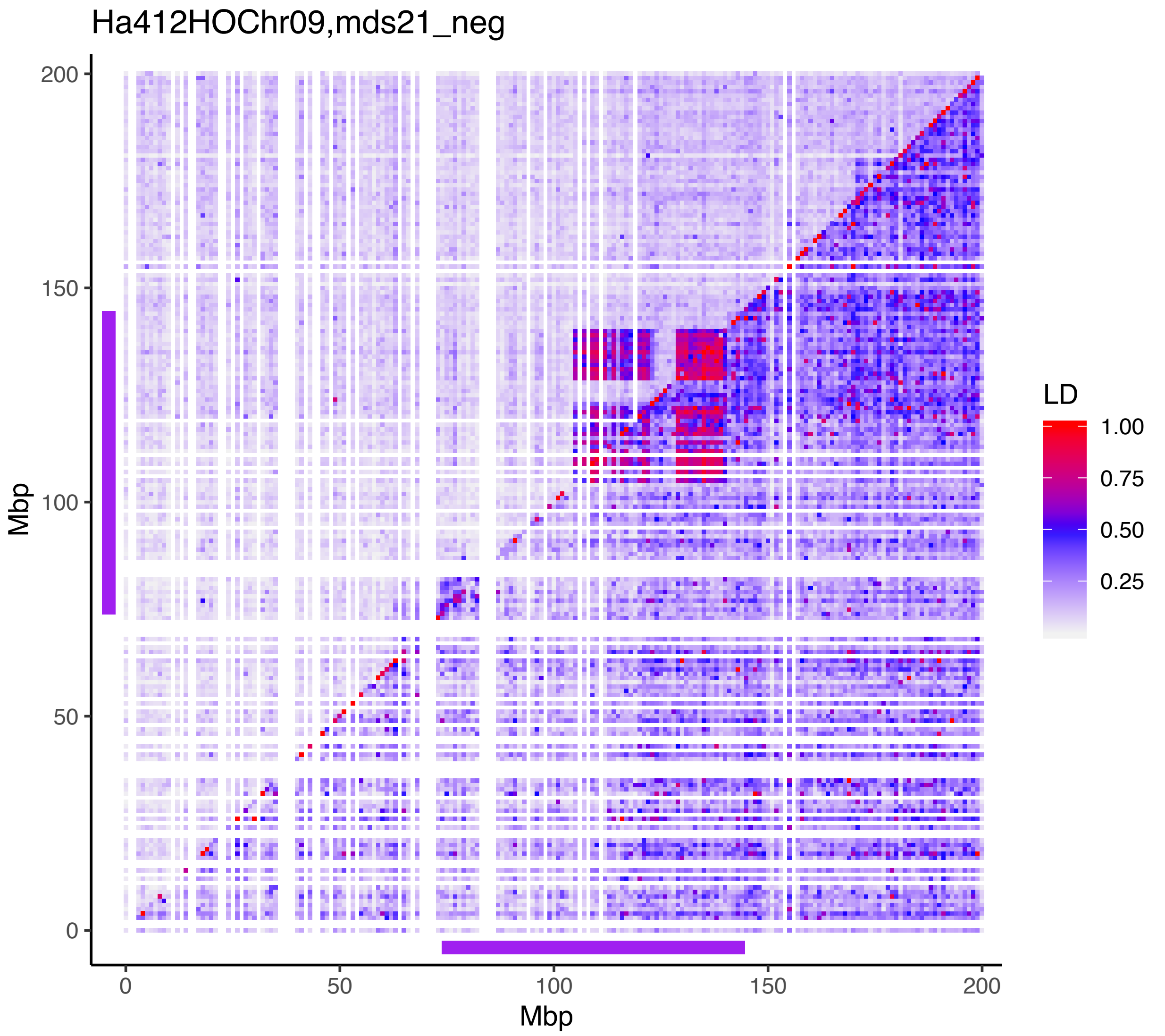

Ha412HOChr09,mds21_neg
Mbp
Mbp
LD
1.00
0.75
0.50
0.25
0
50
100
150
200

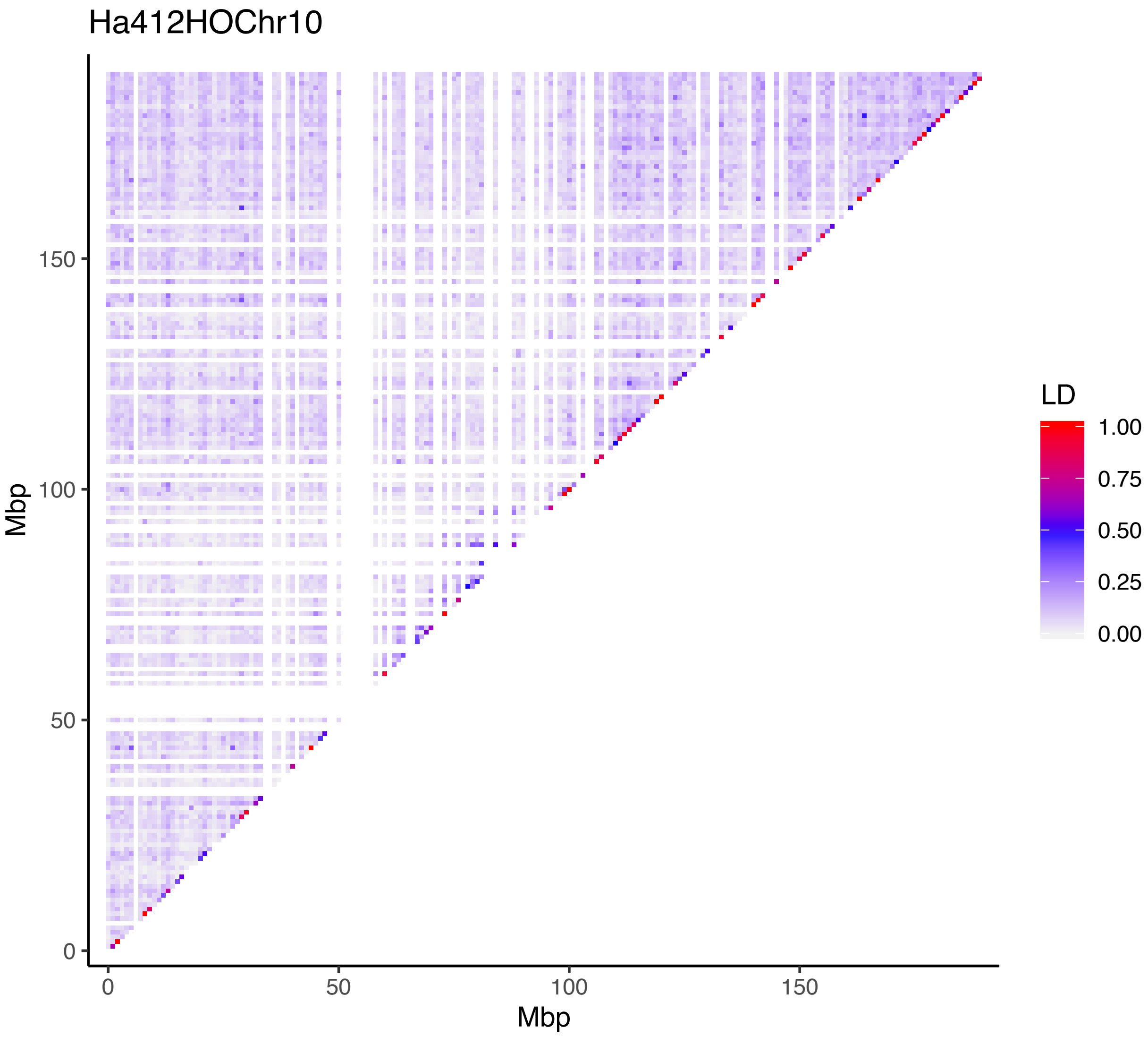
Ha412HOChr10
Mbp
0
50
100
150
Mbp
0
50
100
150
LD
1.00
0.75
0.50
0.25
0.00

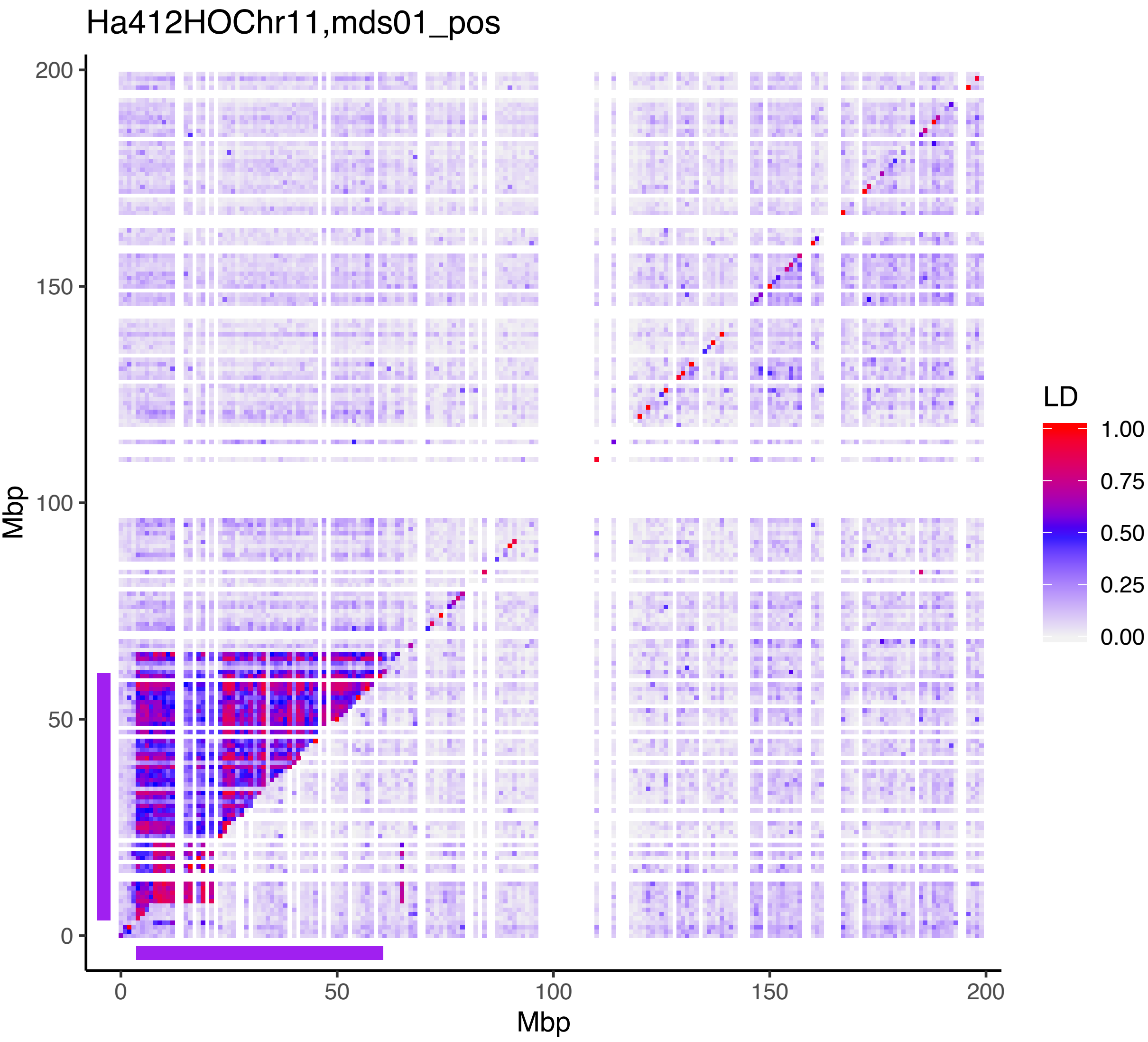

Ha412HOChr11,mds01_pos
Mbp
0
50
100
150
200
LD
1.00
0.75
0.50
0.25
0.00

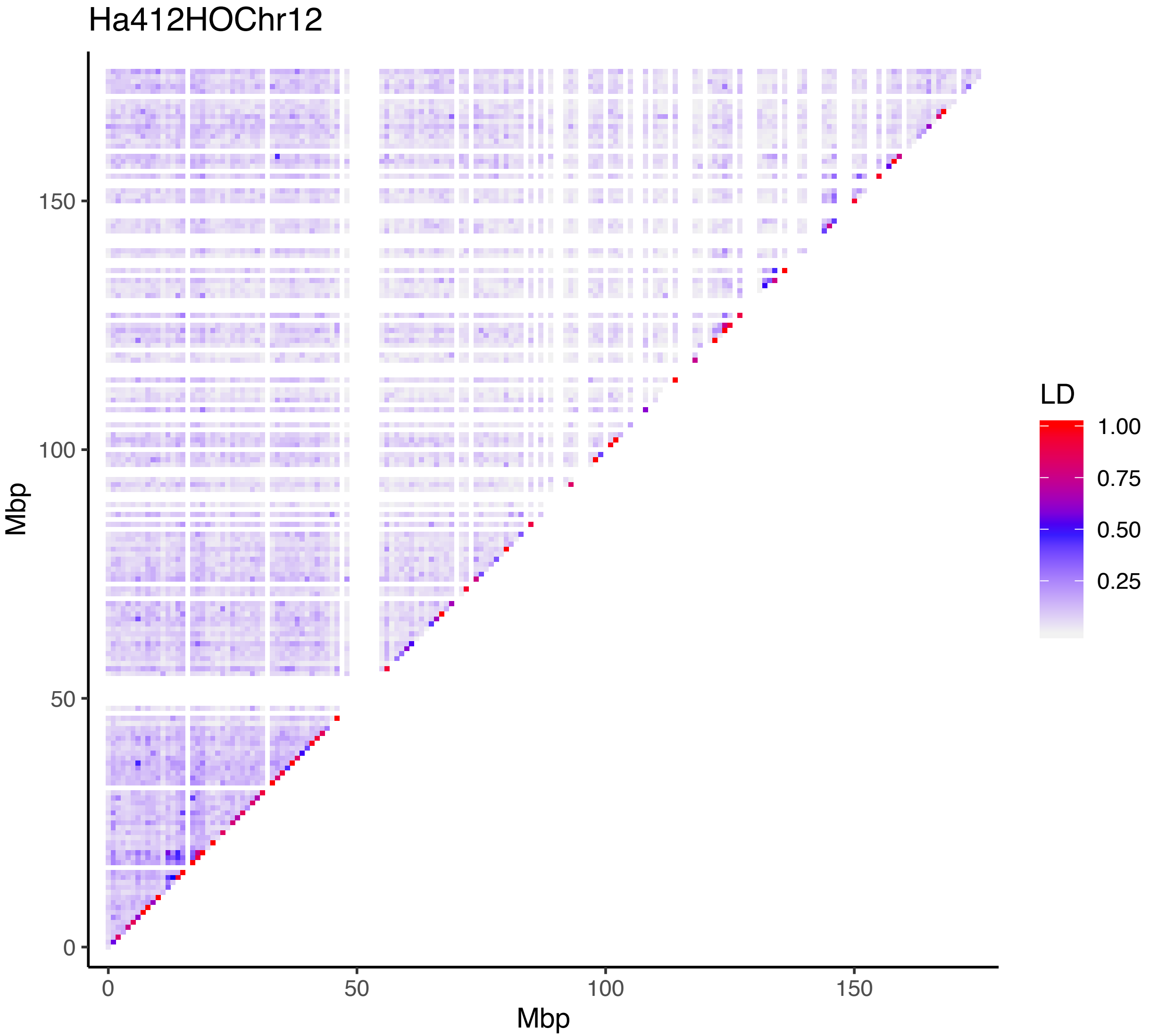
Ha412HOChr12
Mbp
Mbp
0
50
100
150
0
50
100
150
LD
1.00
0.75
0.50
0.25

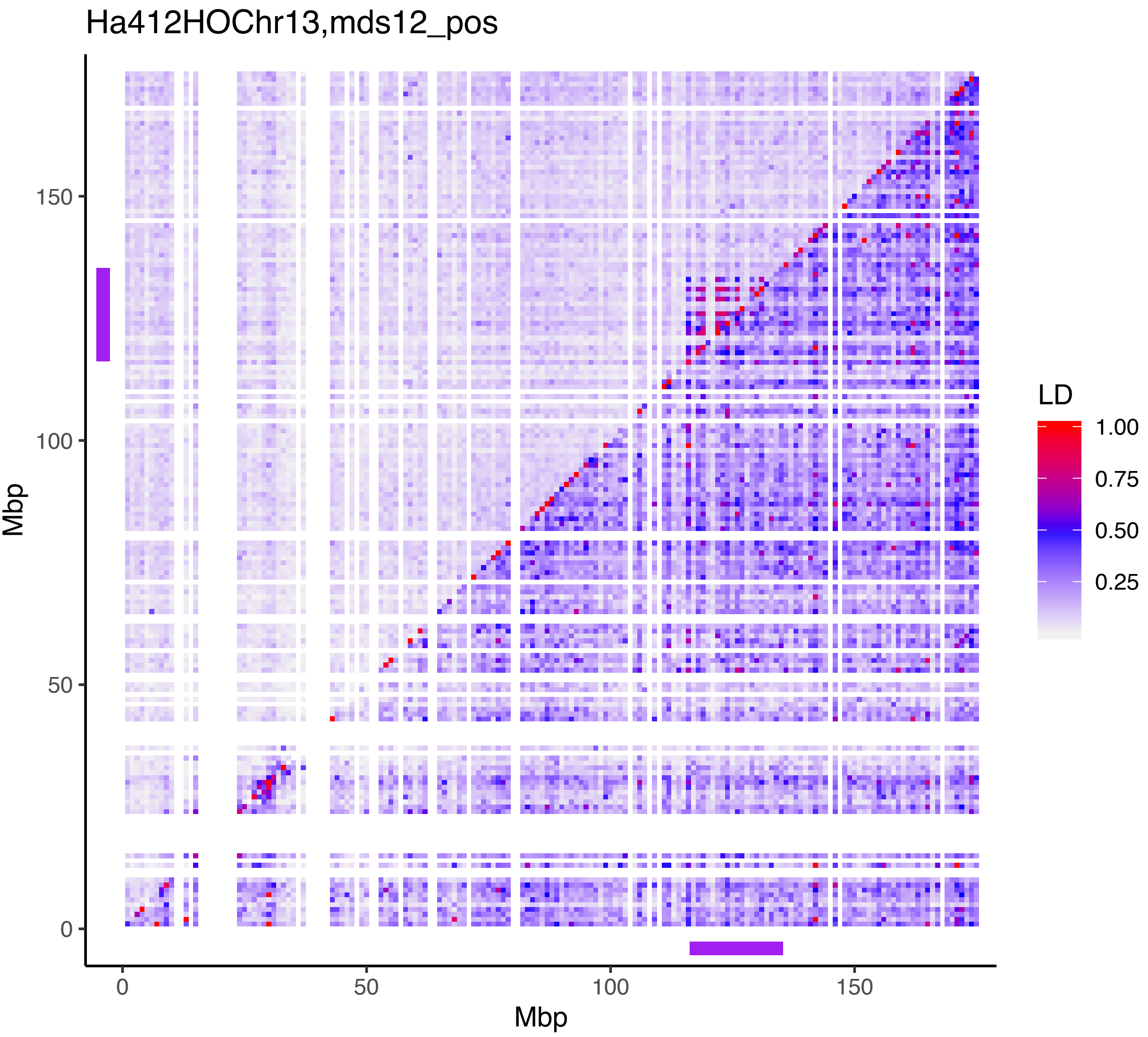

Ha412HOChr13,mds12_pos
Mbp
150
100
50
0
0
50
100
150
Mbp
LD
1.00
0.75
0.50
0.25

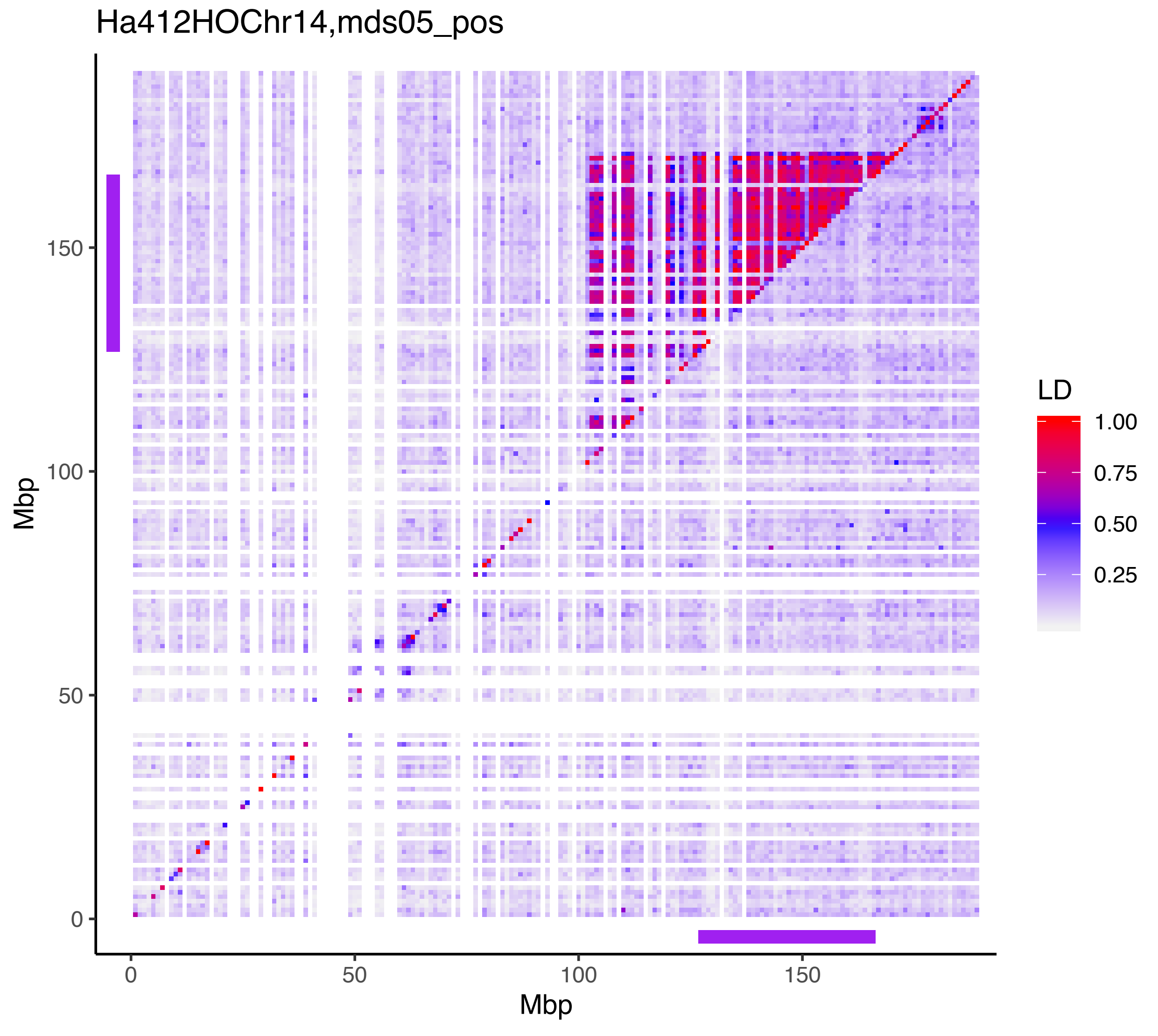
Ha412HOChr14,mds05_pos
Mbp
0
50
100
150
Mbp
0
50
100
150
LD
1.00
0.75
0.50
0.25

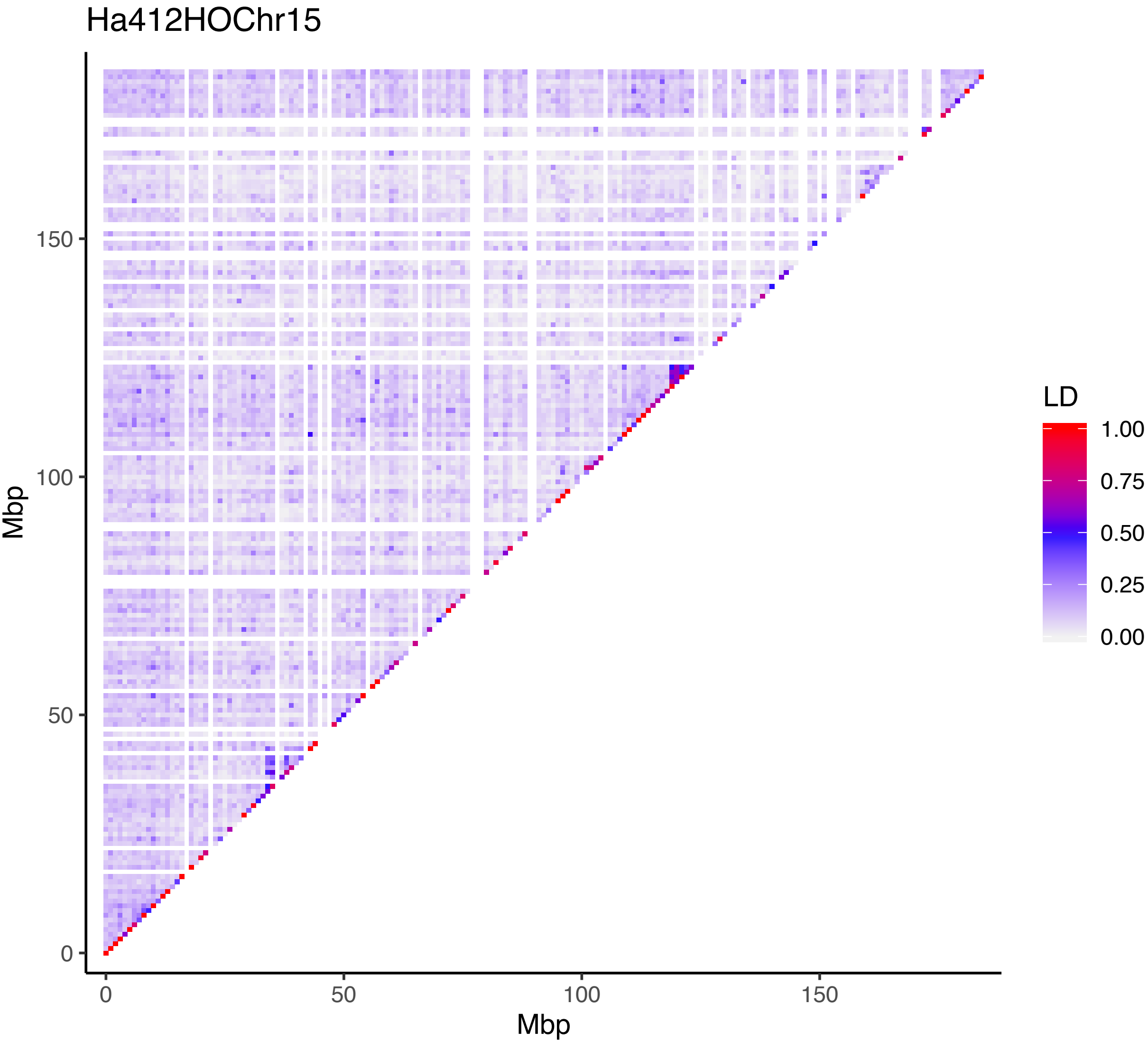
Ha412HOChr15
LD
1.00
0.75
0.50
0.25
0.00
Mbp
150
100
50
0
0
50
100
150
Mbp

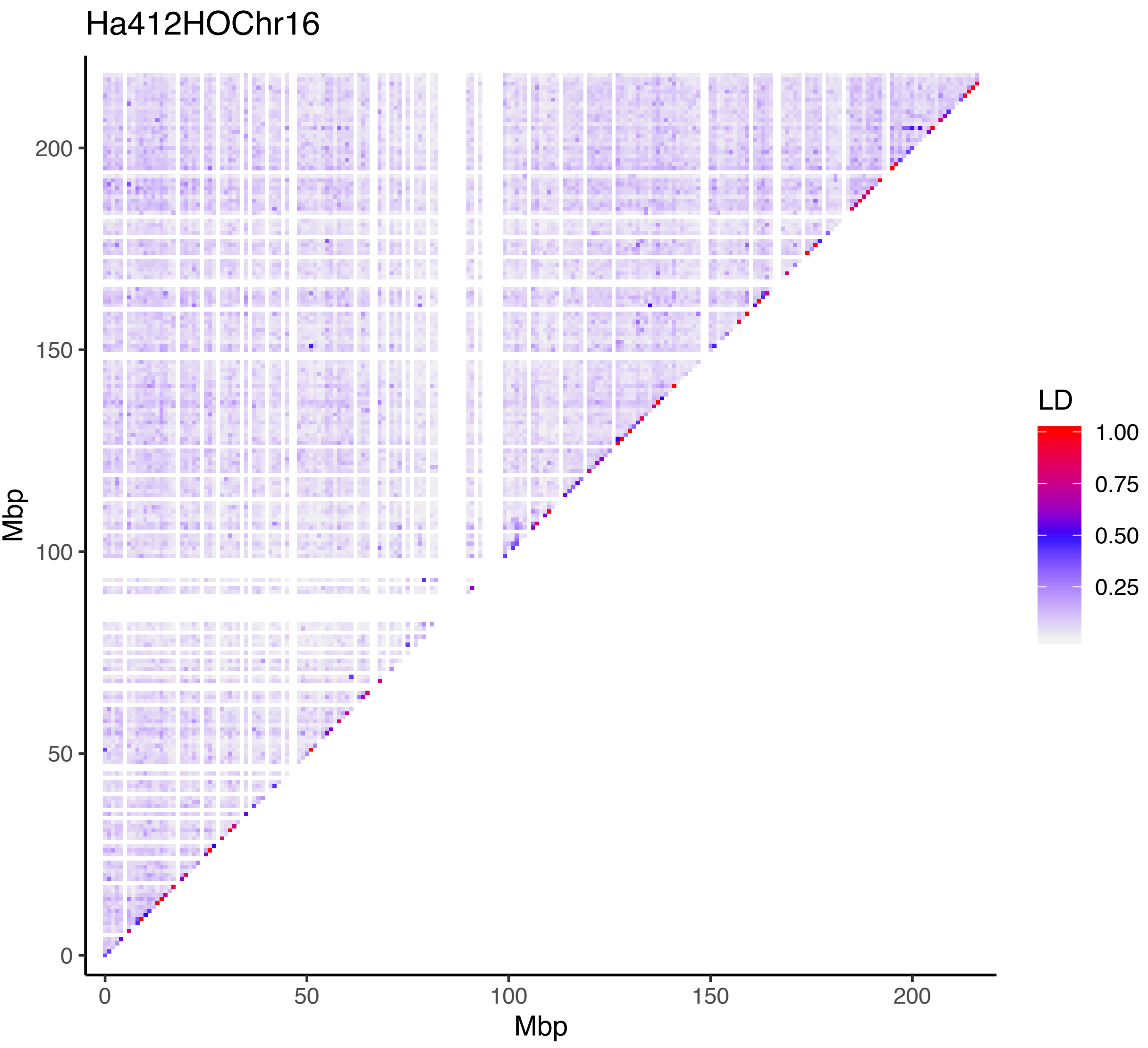
Ha412HOChr16
Mbp
0
50
100
150
200
Mbp
0
50
100
150
200
LD
1.00
0.75
0.50
0.25

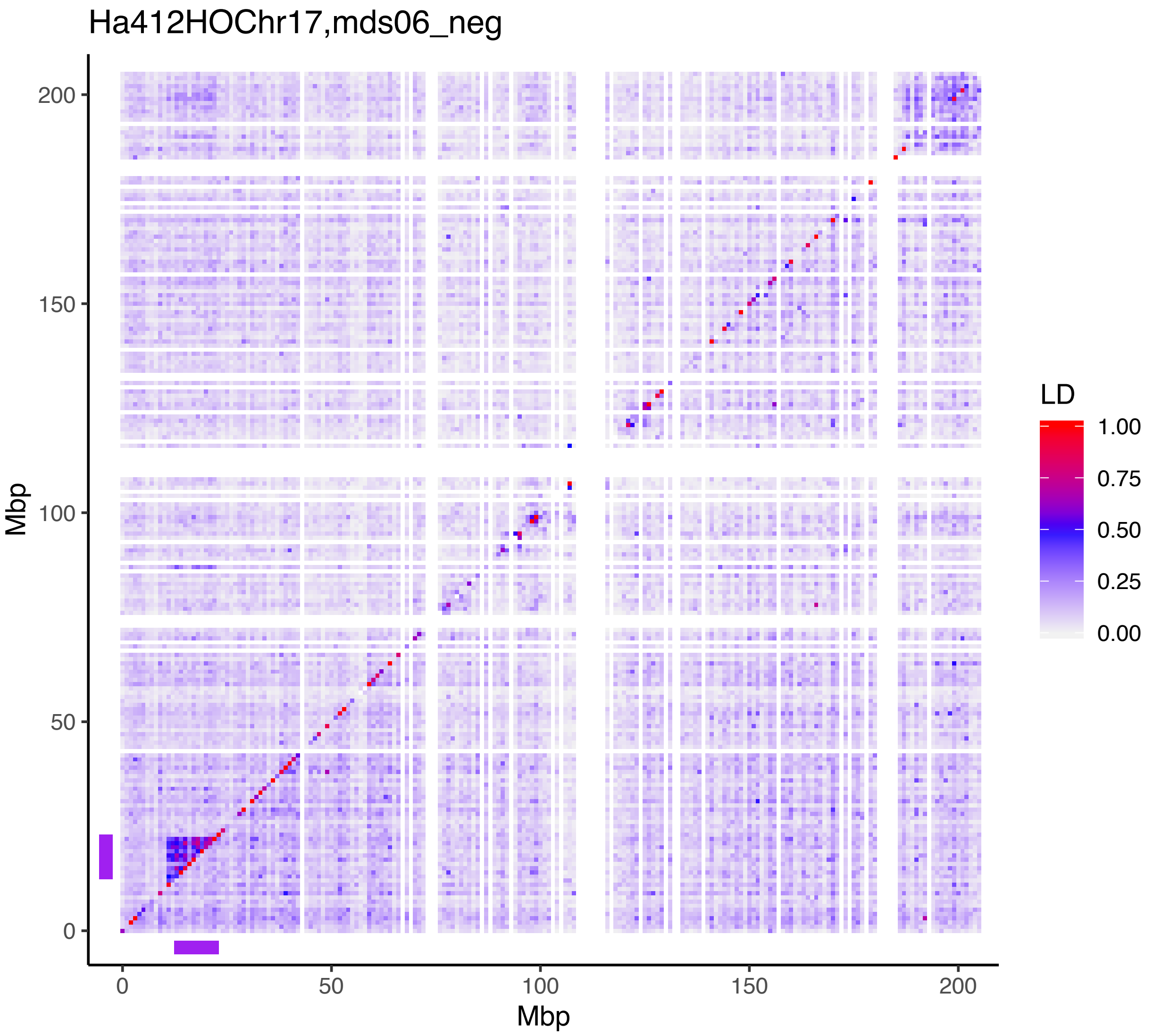

Ha412HOChr17,mds06_neg
Mbp
0
50
100
150
200
LD
1.00
0.75
0.50
0.25
0.00
