## Supporting Information Figure S3 for "Multiple chromosomal inversions contribute to adaptive divergence of a dune sunflower ecotype"

**FIGURE S3** Maps of Great Sand Dunes National Park showing genotype distributions of CHR07.01, CHR17.01 and CHR09.02. Genotypes are based on k-means cluster assignment in PCA. The sand dunes are represented by barren land surrounded by shrubby habitat in the map

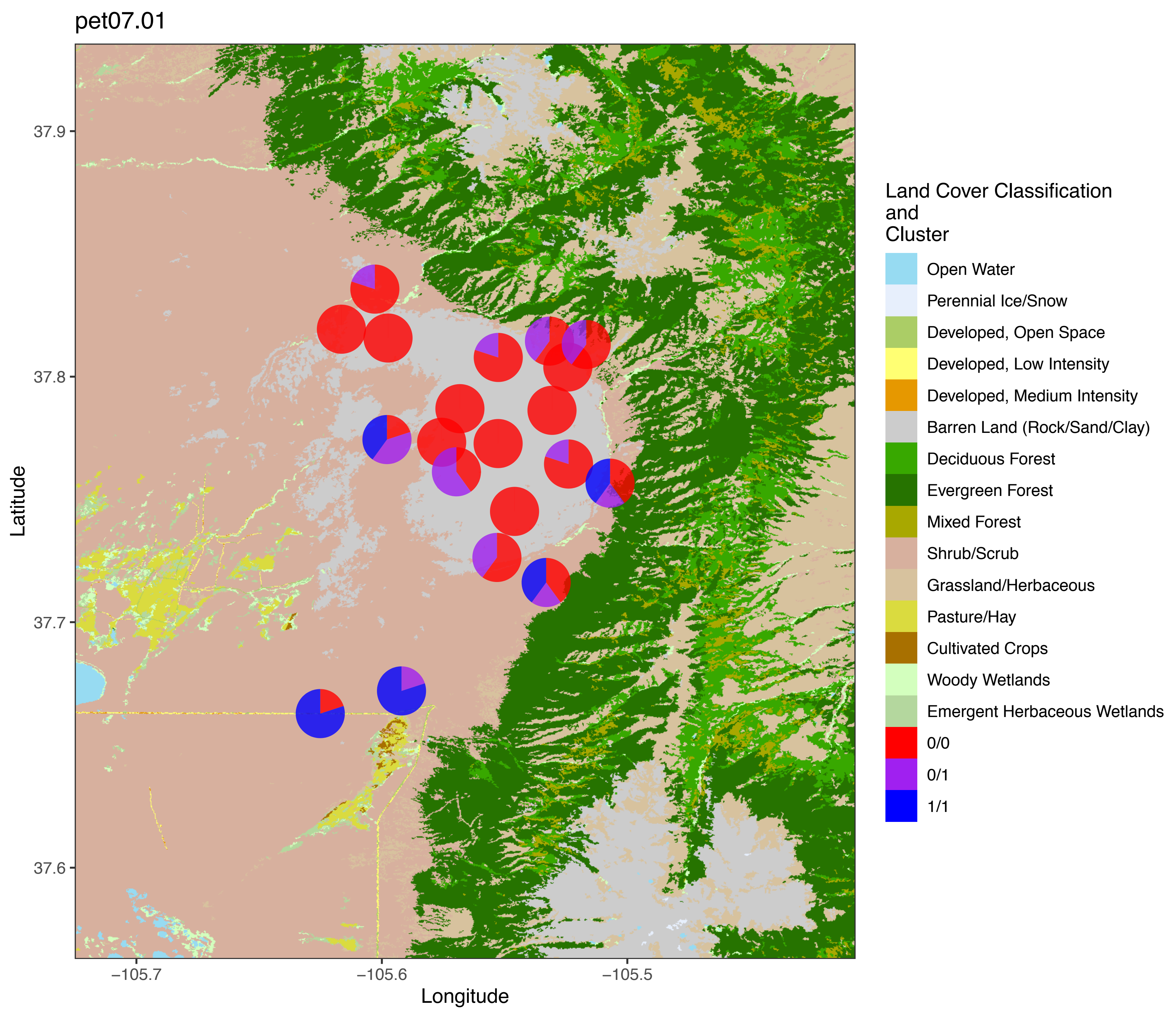
pet07.01
37.9
37.8
37.7
37.6
Latitude
−105.7
−105.6
−105.5
Longitude
Land Cover Classification
and
Cluster
Open Water
Perennial Ice/Snow
Developed, Open Space
Developed, Low Intensity
Developed, Medium Intensity
Barren Land (Rock/Sand/Clay)
Deciduous Forest
Evergreen Forest
Mixed Forest
Shrub/Scrub
Grassland/Herbaceous
Pasture/Hay
Cultivated Crops
Woody Wetlands
Emergent Herbaceous Wetlands
0/0
0/1
1/1

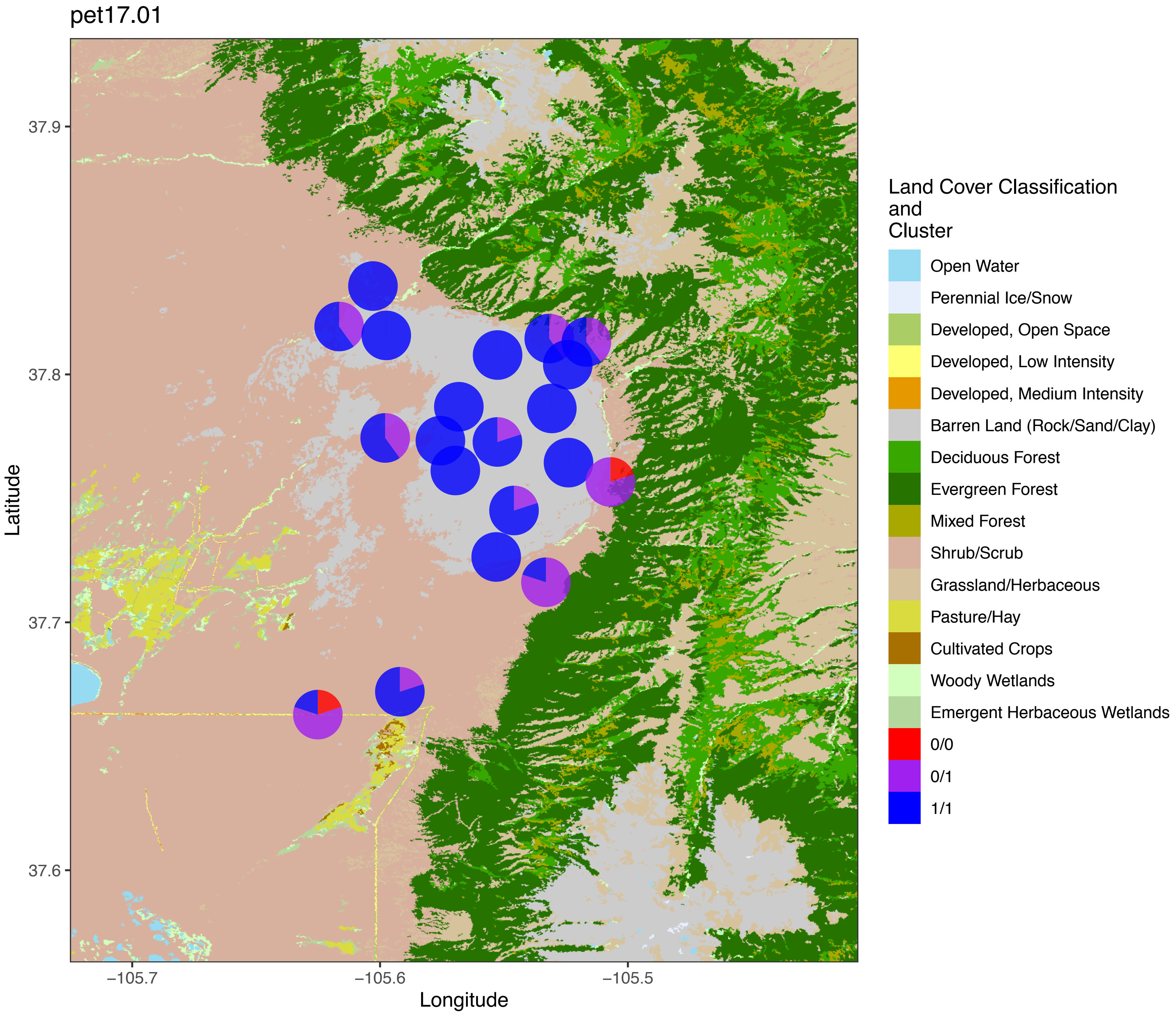

pet17.01
37.9
37.8
37.7
37.6
Latitude
−105.7
−105.6
−105.5
Longitude
Land Cover Classification
and
Cluster
Open Water
Perennial Ice/Snow
Developed, Open Space
Developed, Low Intensity
Developed, Medium Intensity
Barren Land (Rock/Sand/Clay)
Deciduous Forest
Evergreen Forest
Mixed Forest
Shrub/Scrub
Grassland/Herbaceous
Pasture/Hay
Cultivated Crops
Woody Wetlands
Emergent Herbaceous Wetlands
0/0
0/1
1/1

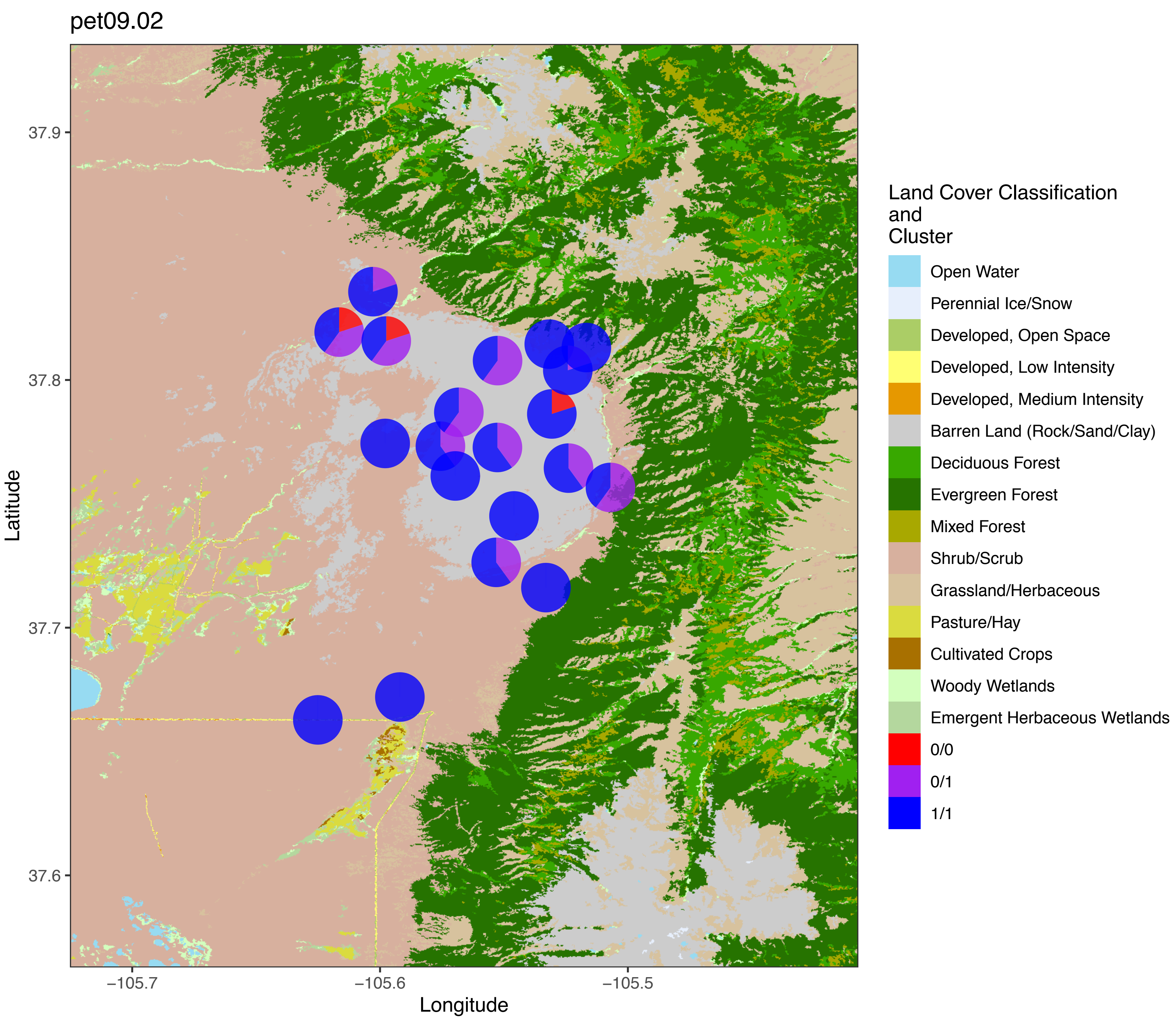
pet09.02
Latitude
37.9
37.8
37.7
37.6
Longitude
−105.7
−105.6
−105.5
Land Cover Classification
and
Cluster
Open Water
Perennial Ice/Snow
Developed, Open Space
Developed, Low Intensity
Developed, Medium Intensity
Barren Land (Rock/Sand/Clay)
Deciduous Forest
Evergreen Forest
Mixed Forest
Shrub/Scrub
Grassland/Herbaceous
Pasture/Hay
Cultivated Crops
Woody Wetlands
Emergent Herbaceous Wetlands
0/0
0/1
1/1
