## Supporting Information Figure S5 for "Multiple chromosomal inversions contribute to adaptive divergence of a dune sunflower ecotype"

**FIGURE S4** Genetic map of dune ecotype compared to the HA412HOv2 reference genome. Marker distances on HA412 are based their physical position, assuming an average recombination rate of 0.5 cM/Mb to make the linkage groups comparable. Linkage groups are named and color-coded based on their the synteny blocks with *Helianthus annuus* chromosomes according to Ostevik et al. (2019). Homologous markers are connected by lines.

### HA412 vs Dune

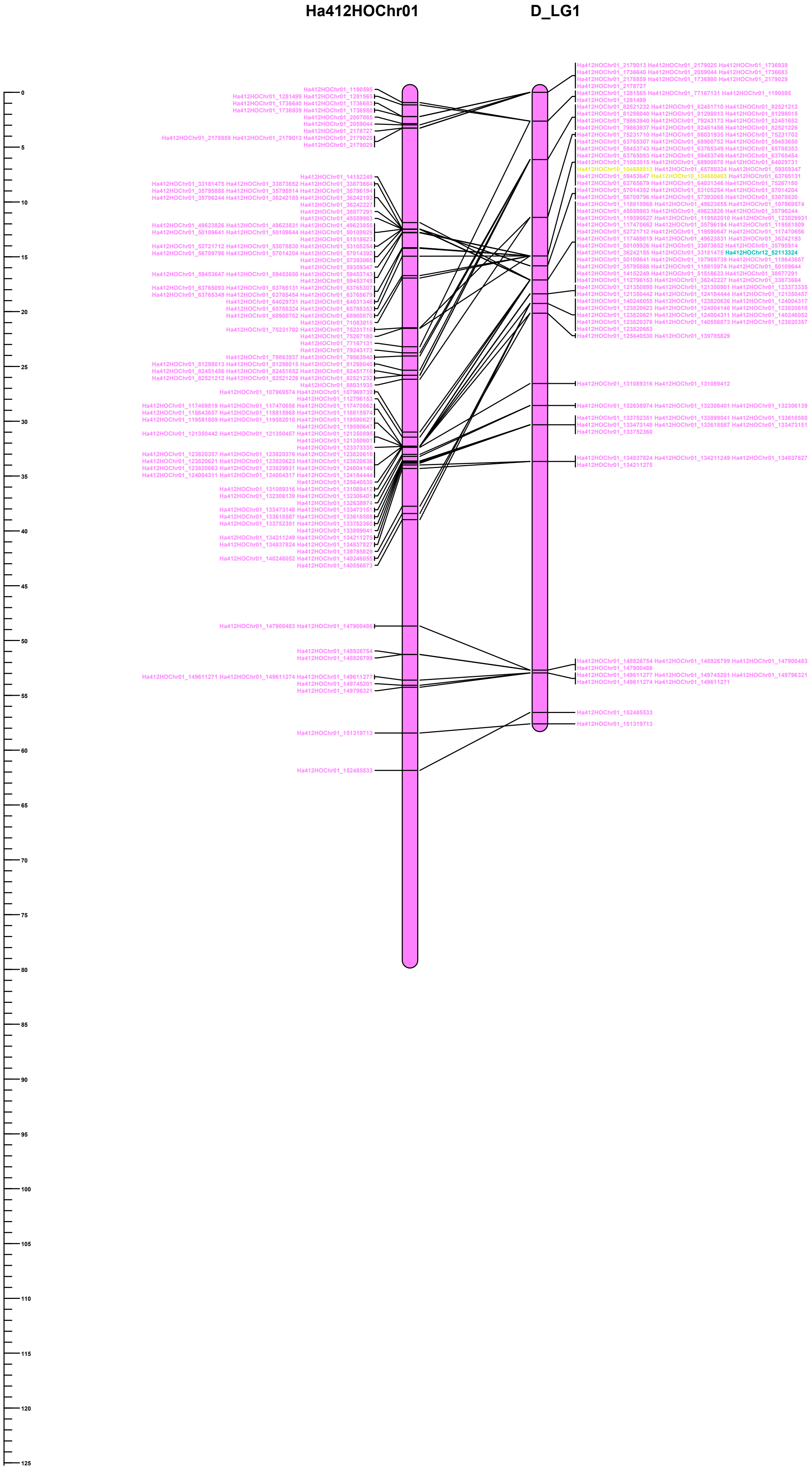

### HA412 vs Dune

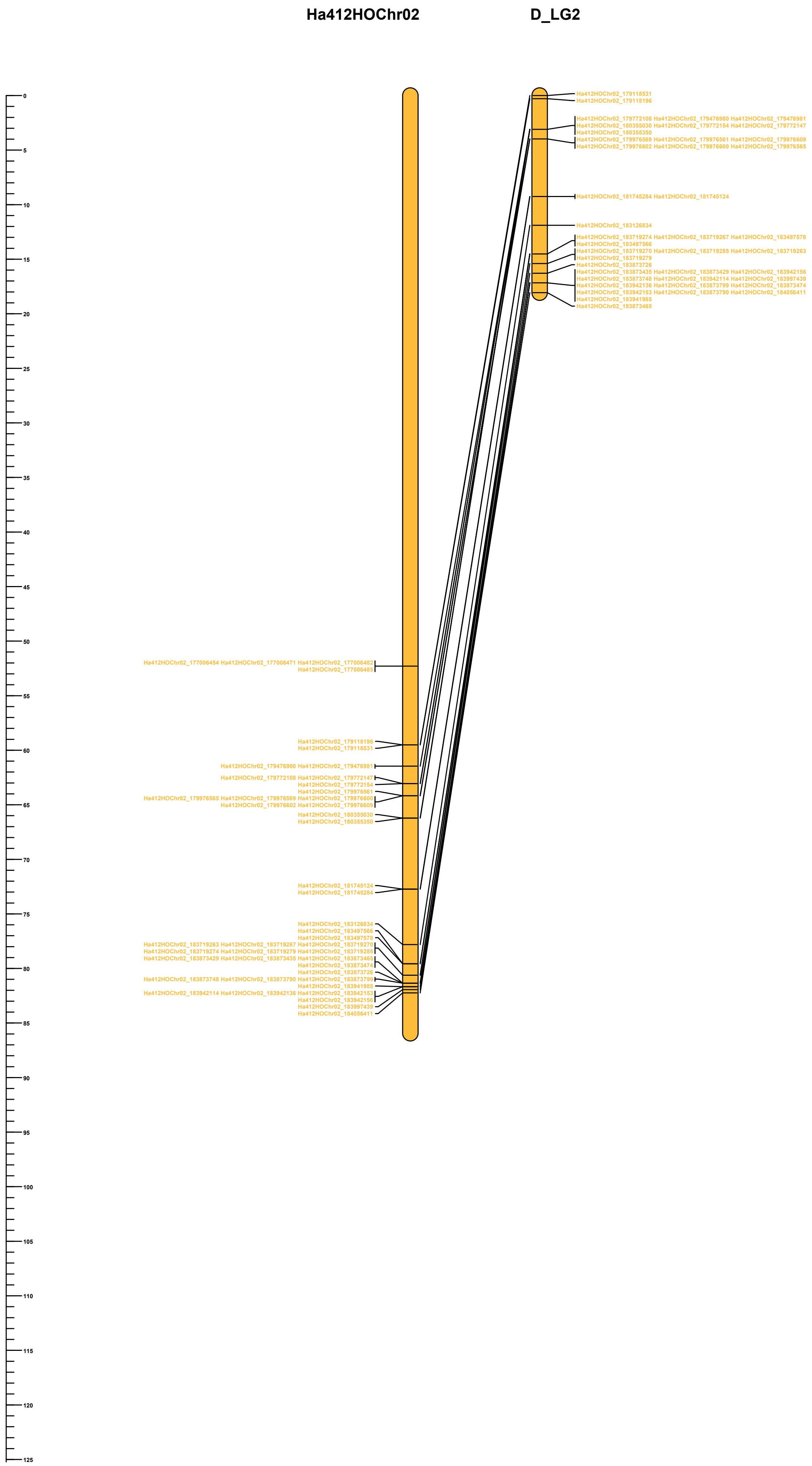

### HA412 vs Dune

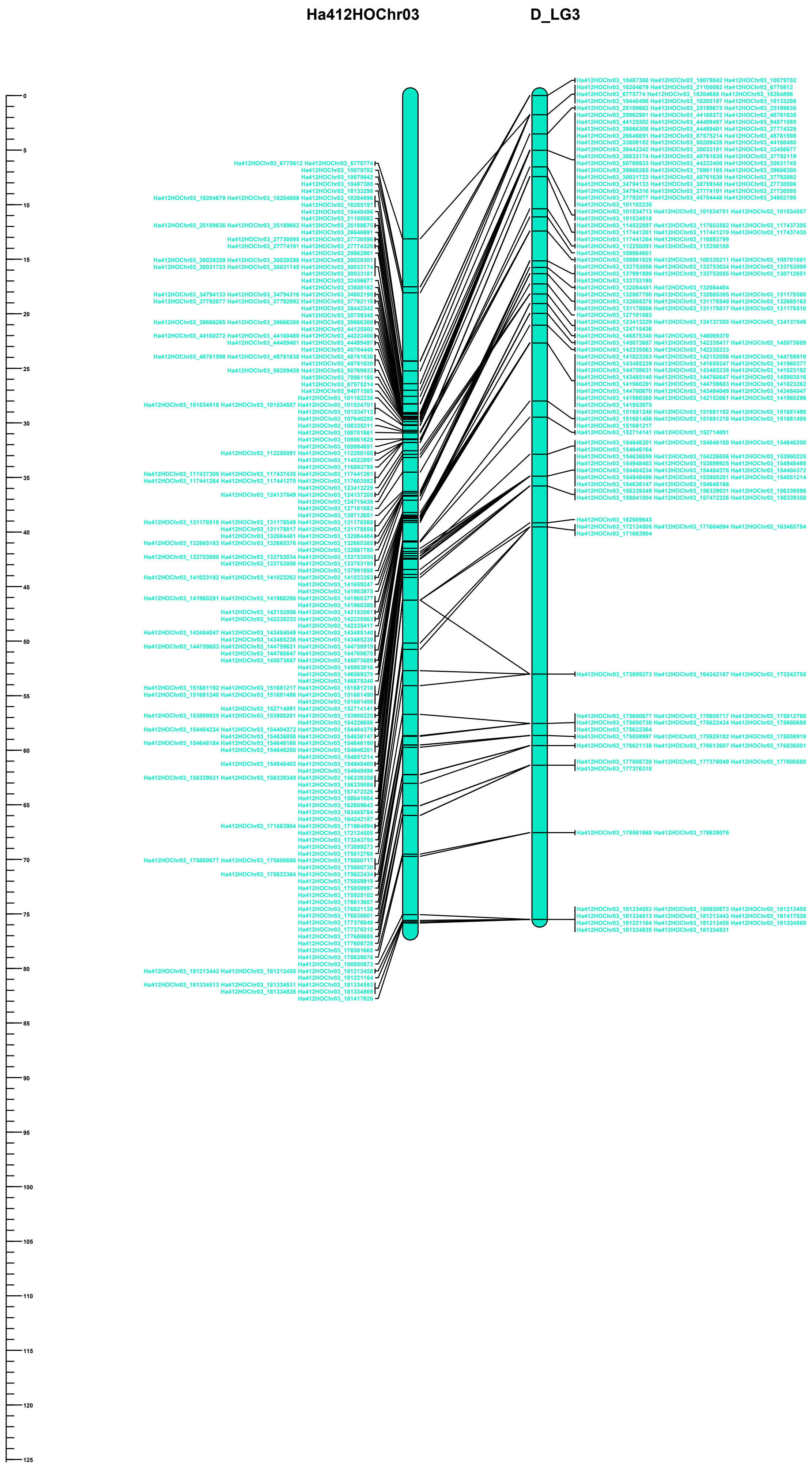

### HA412 vs Dune

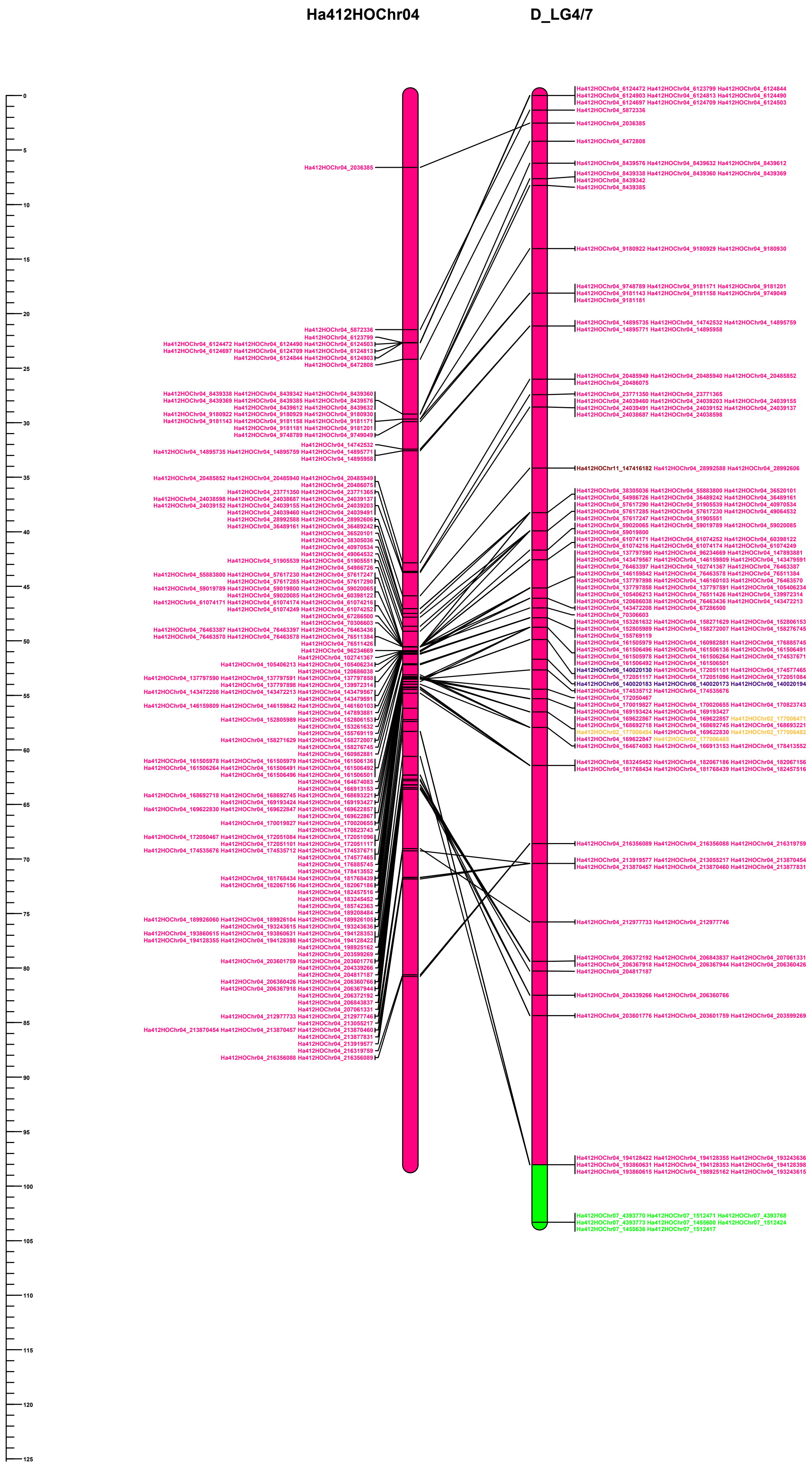

### HA412 vs Dune

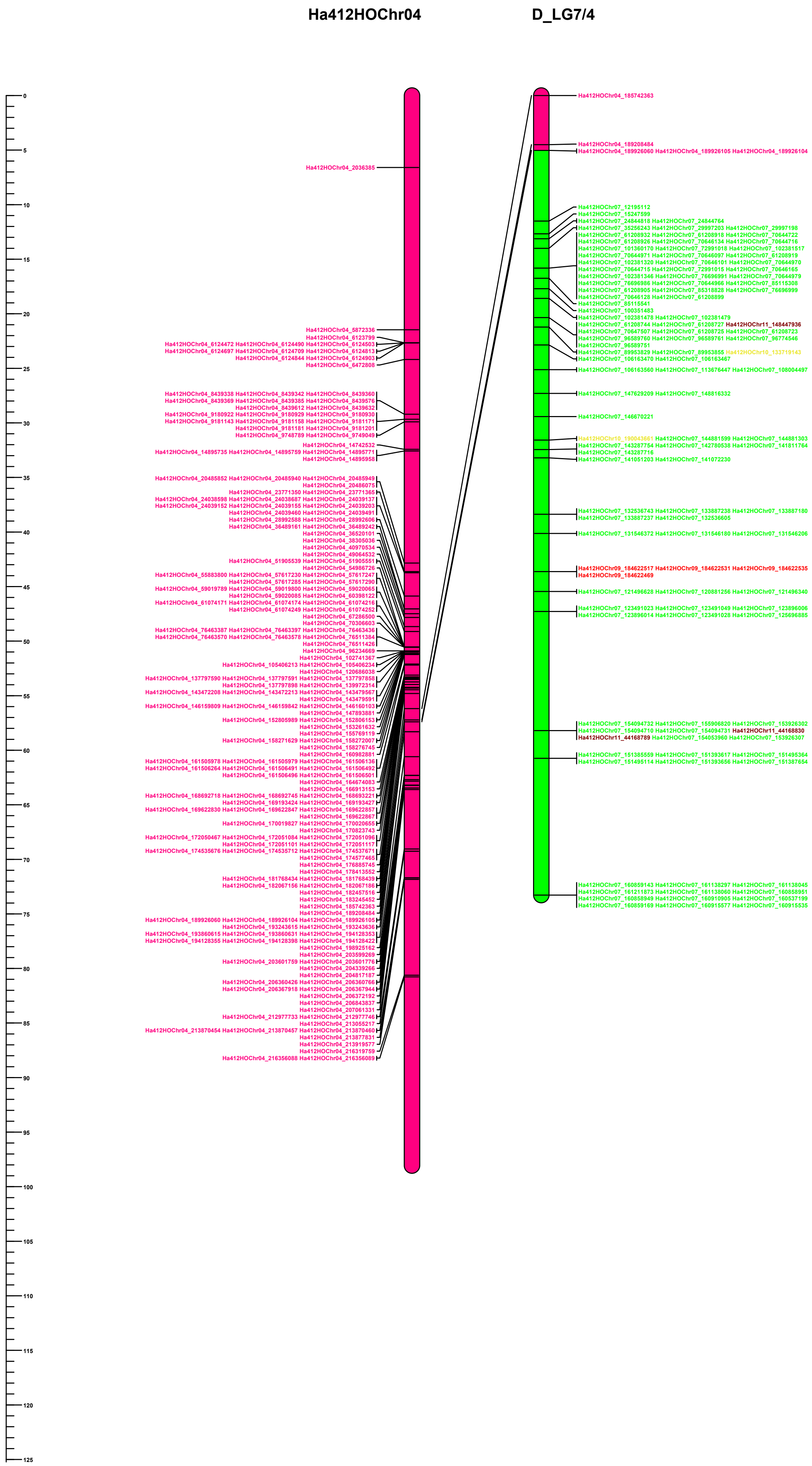

### HA412 vs Dune

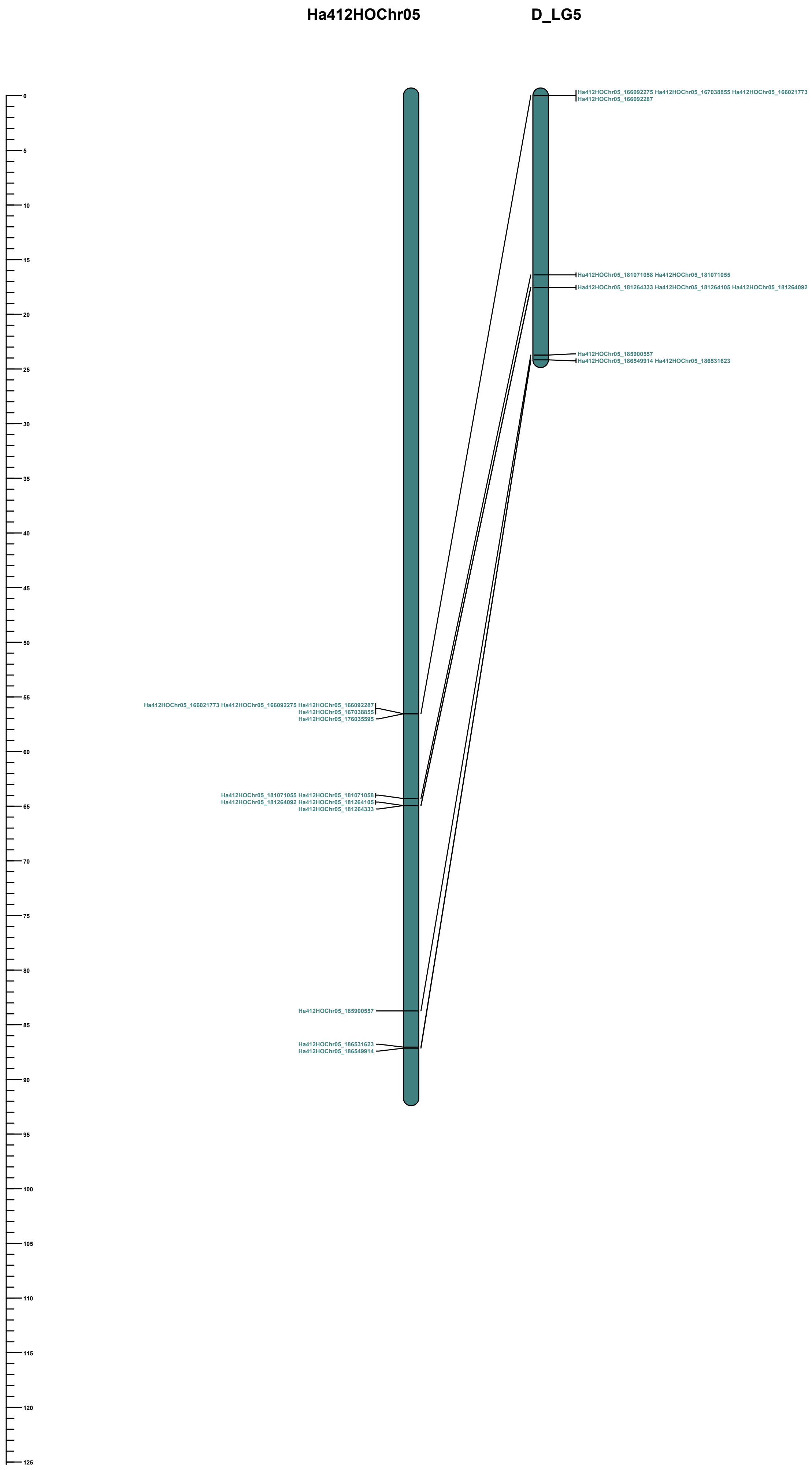

### HA412 vs Dune

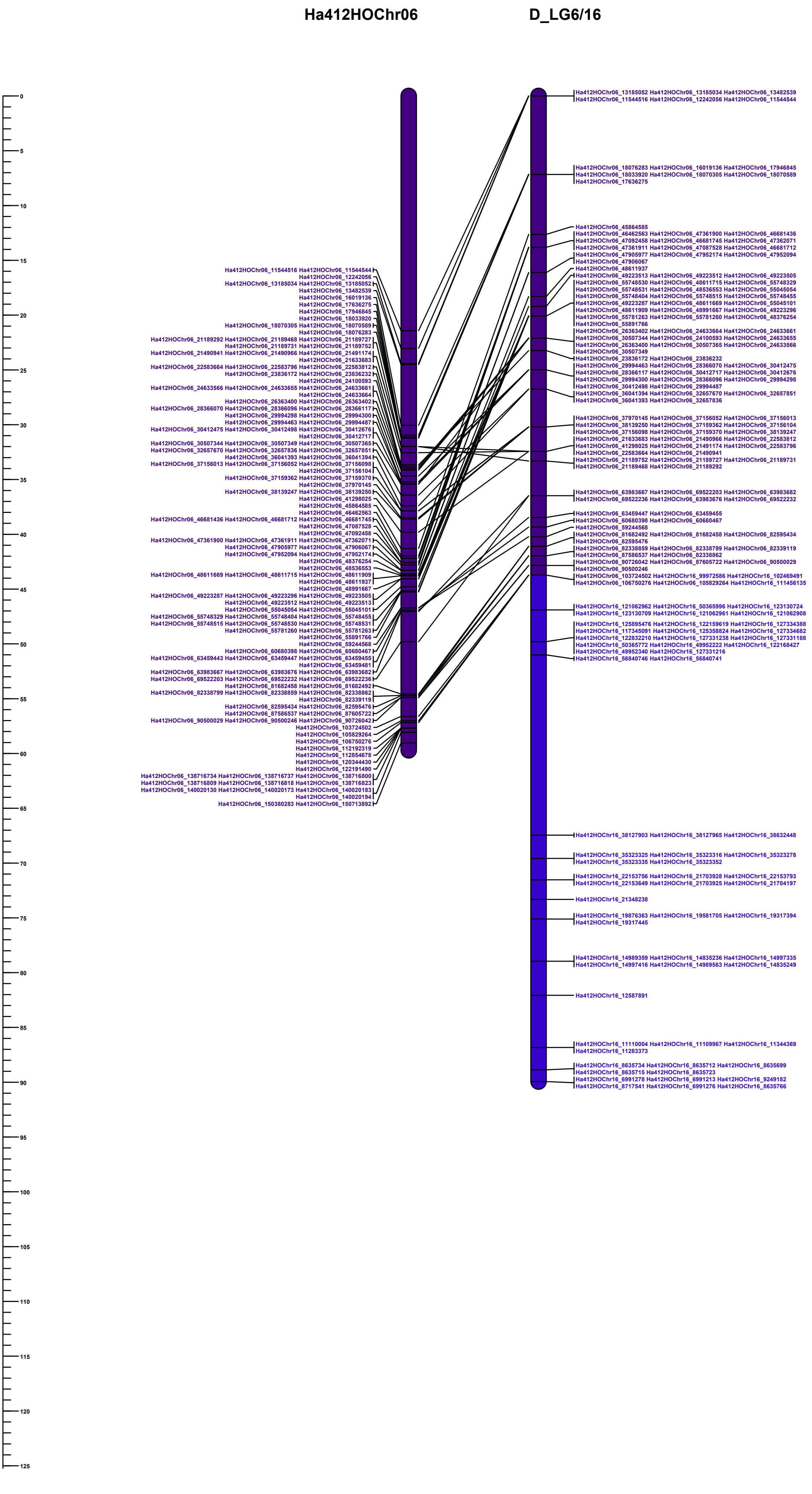

### HA412 vs Dune

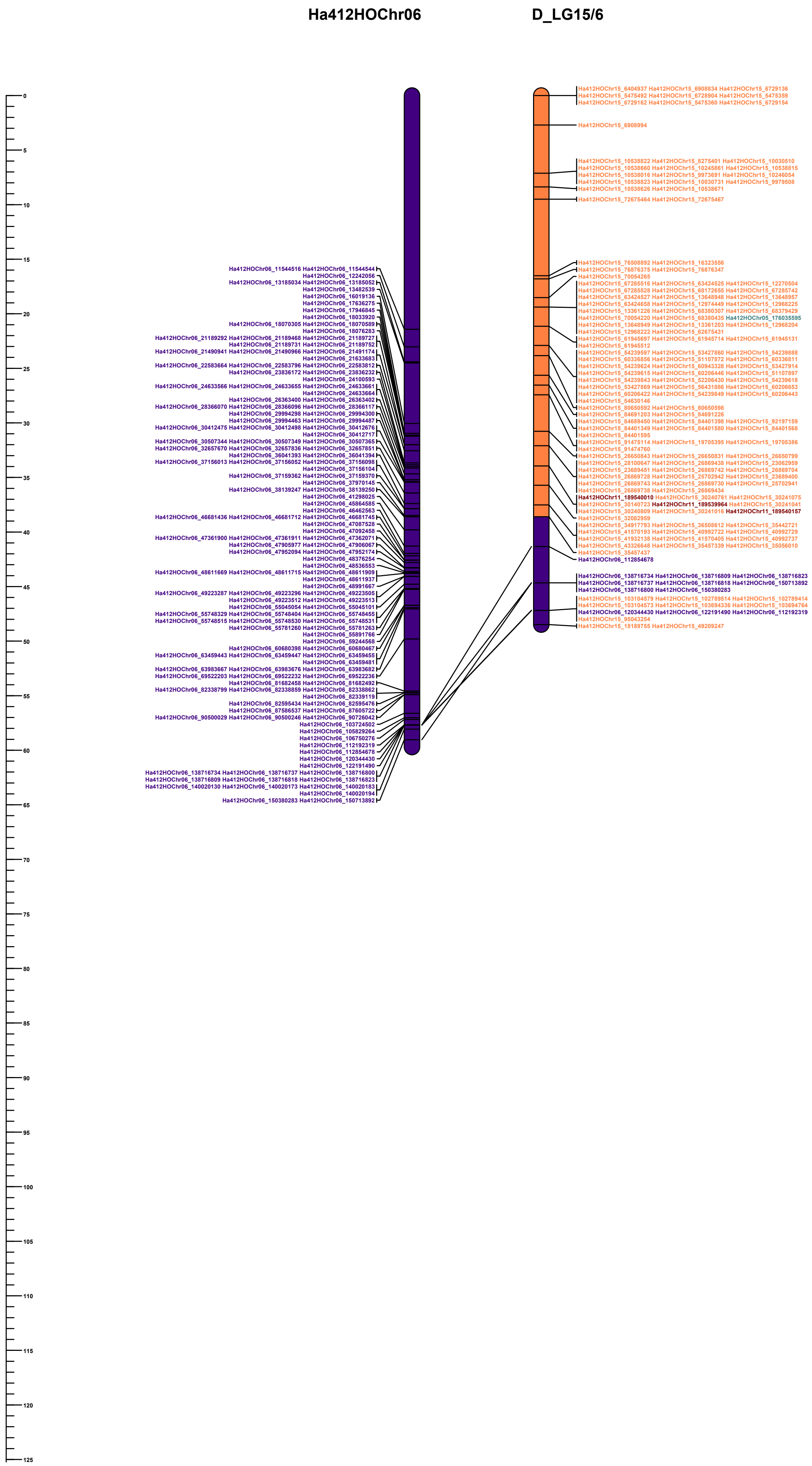

### HA412 vs Dune

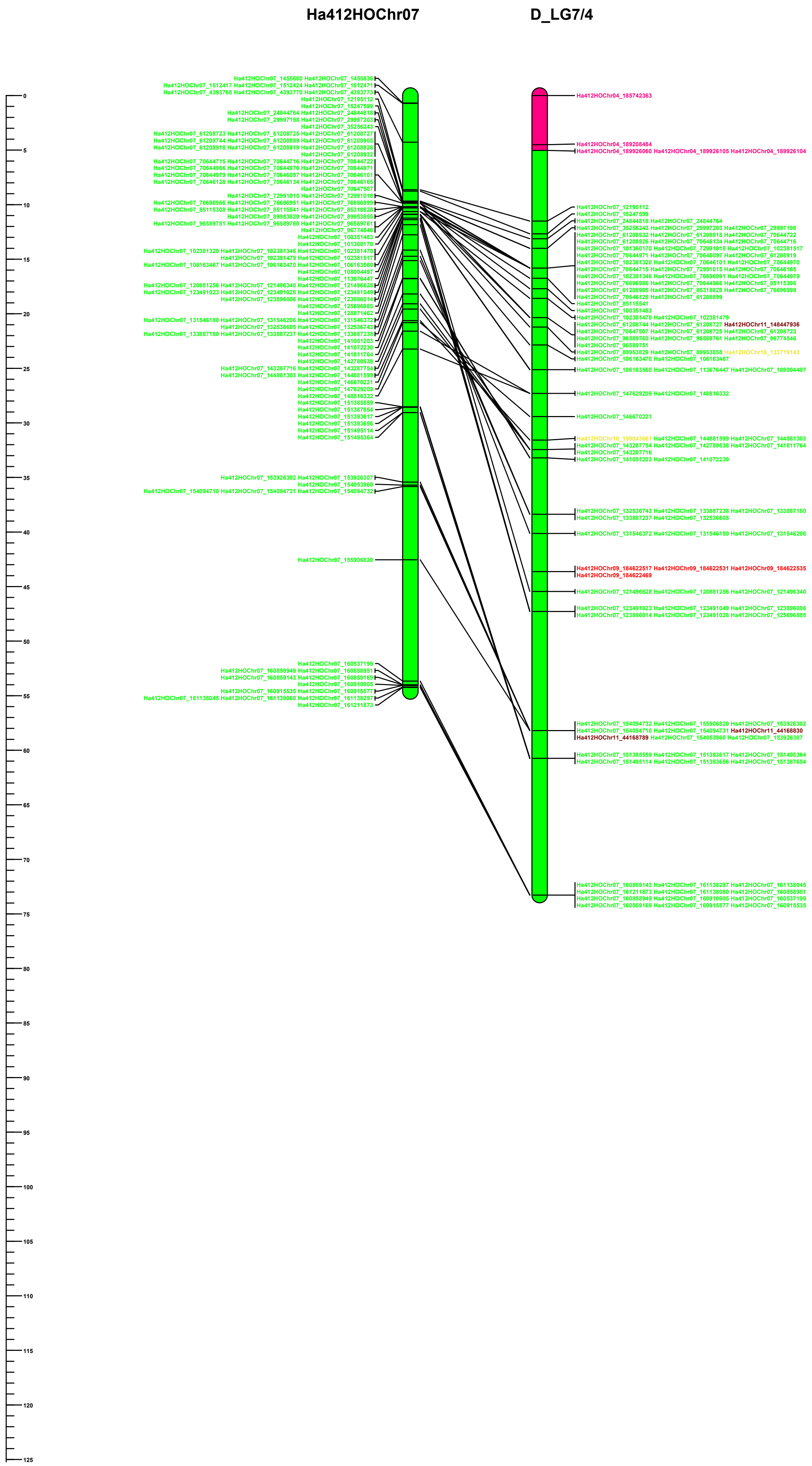

### HA412 vs Dune

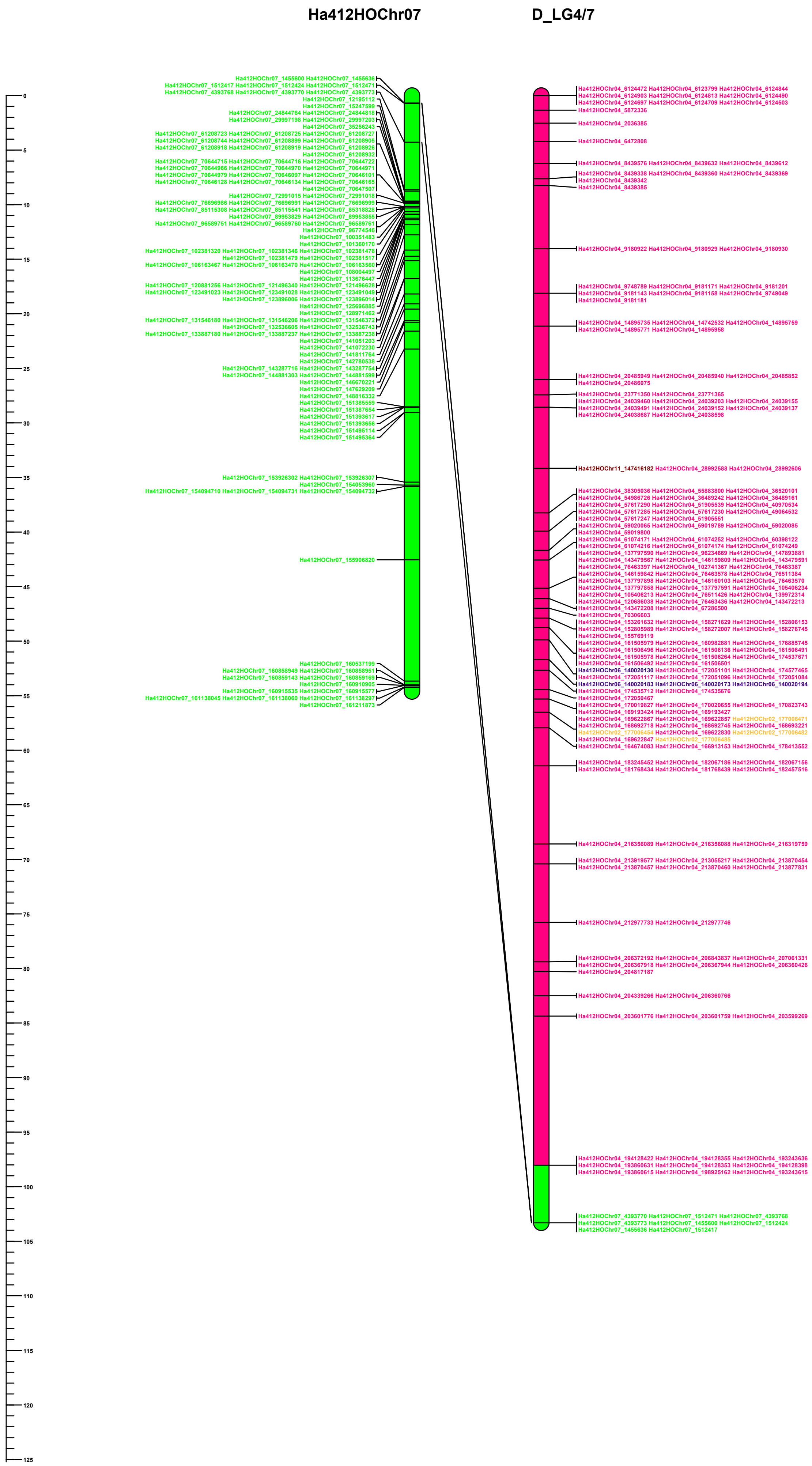

### HA412 vs Dune

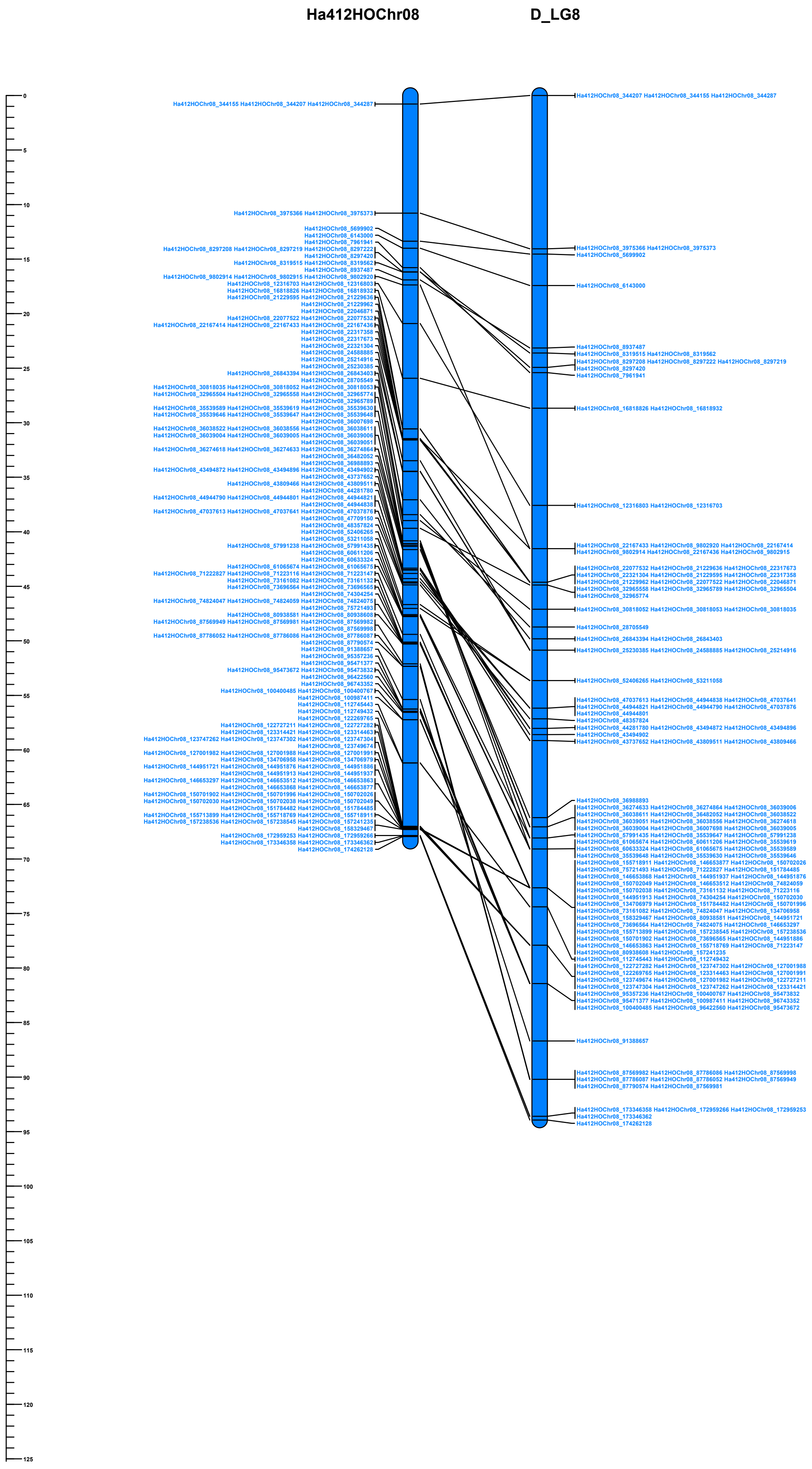

### HA412 vs Dune

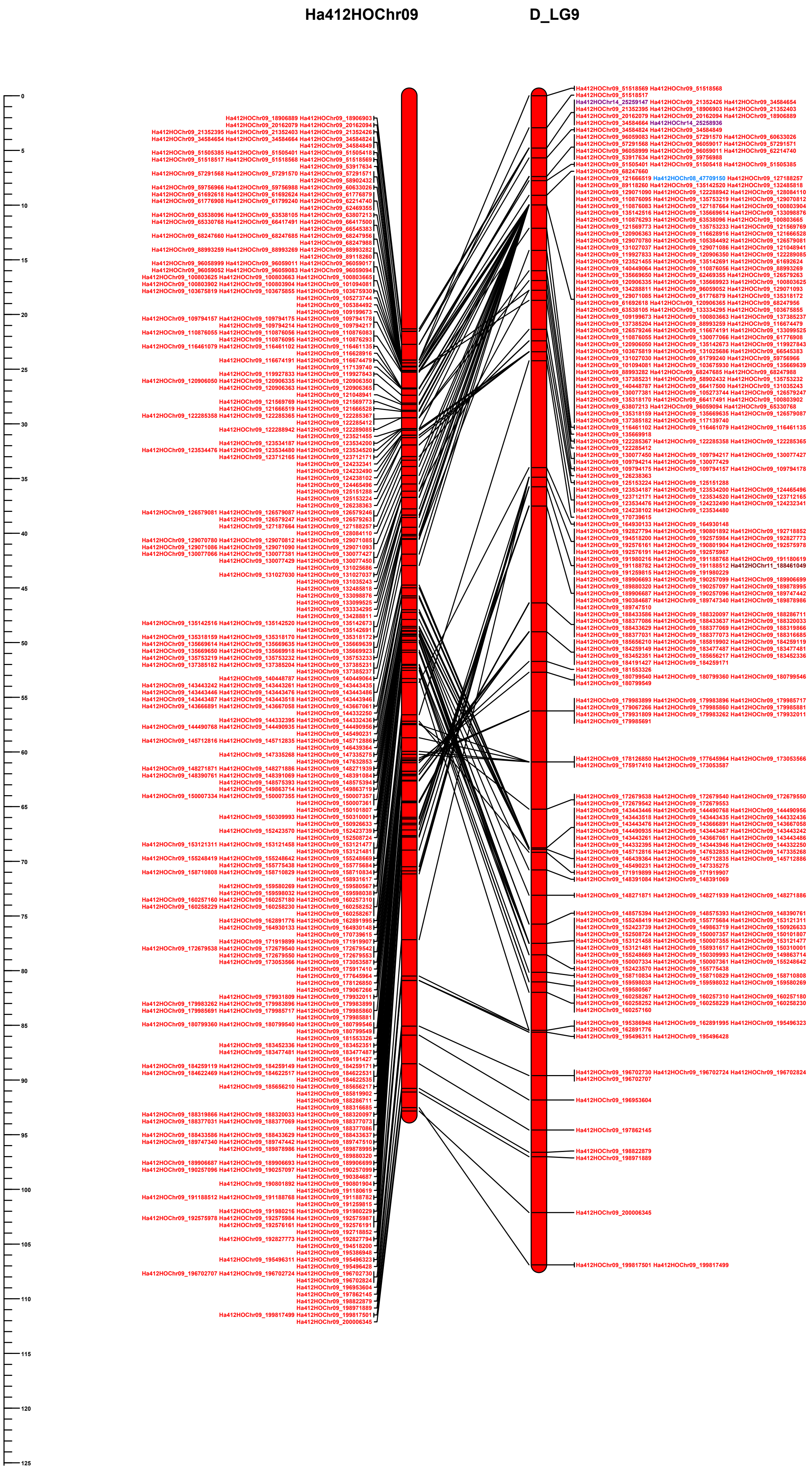

### HA412 vs Dune

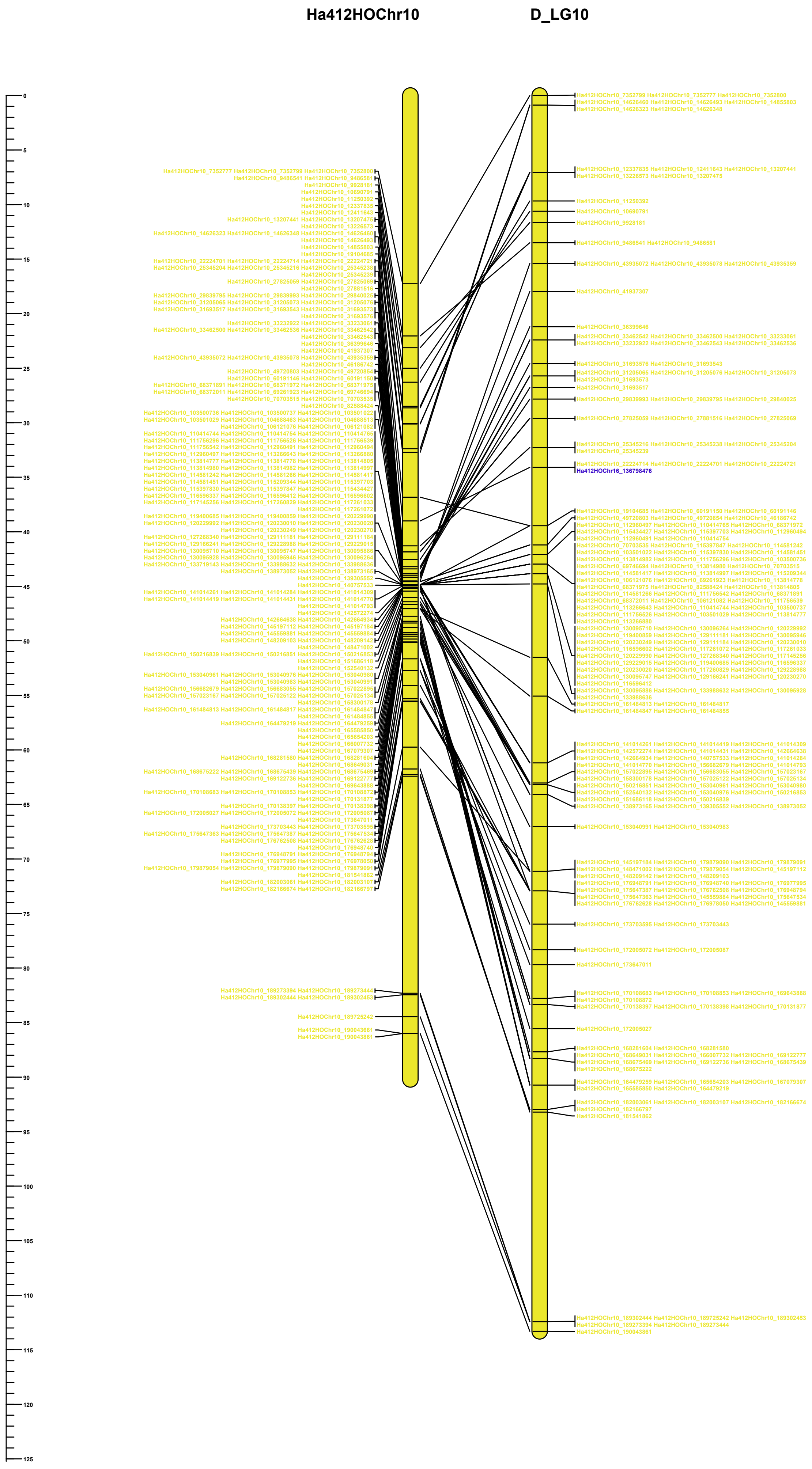

### HA412 vs Dune

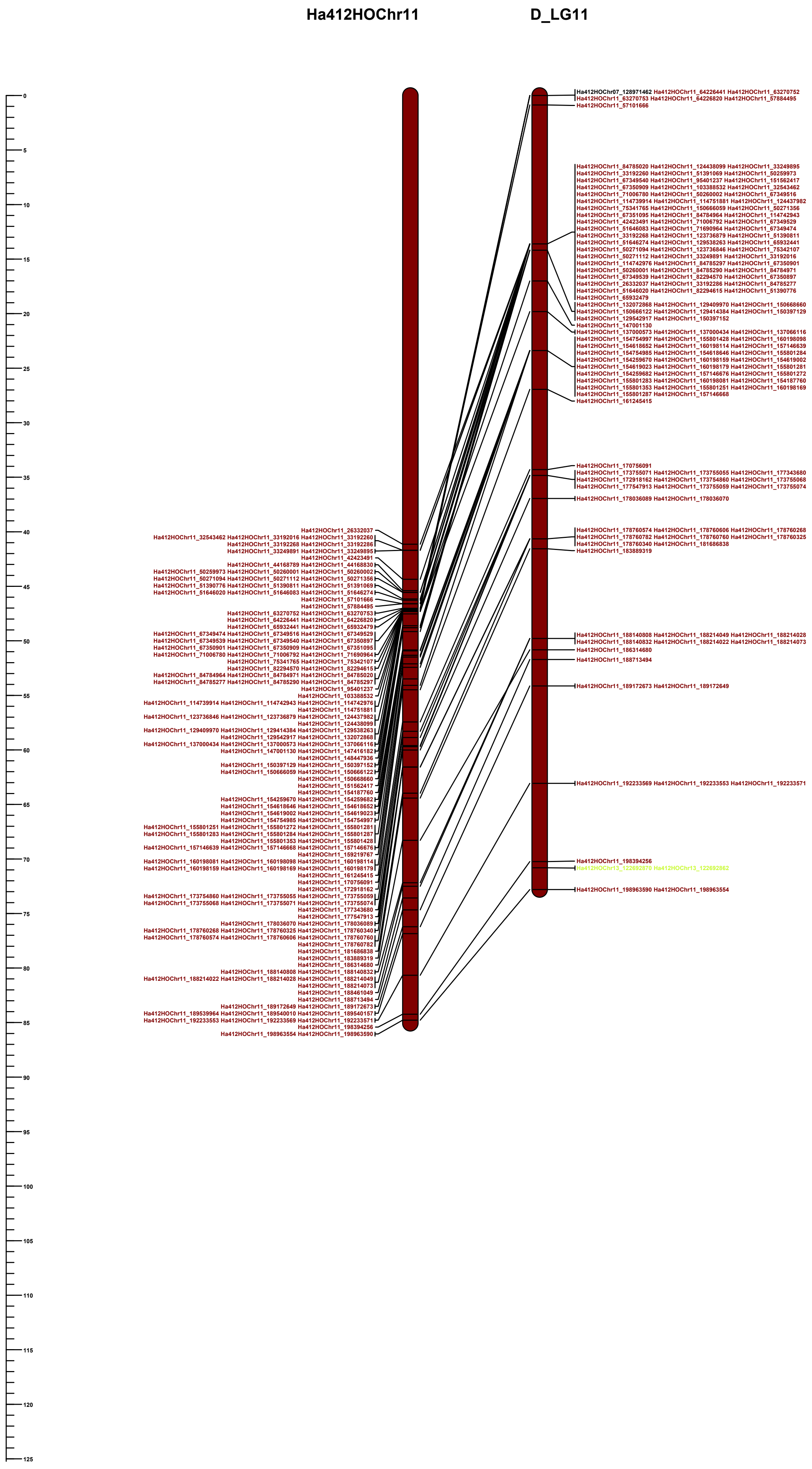

### HA412 vs Dune

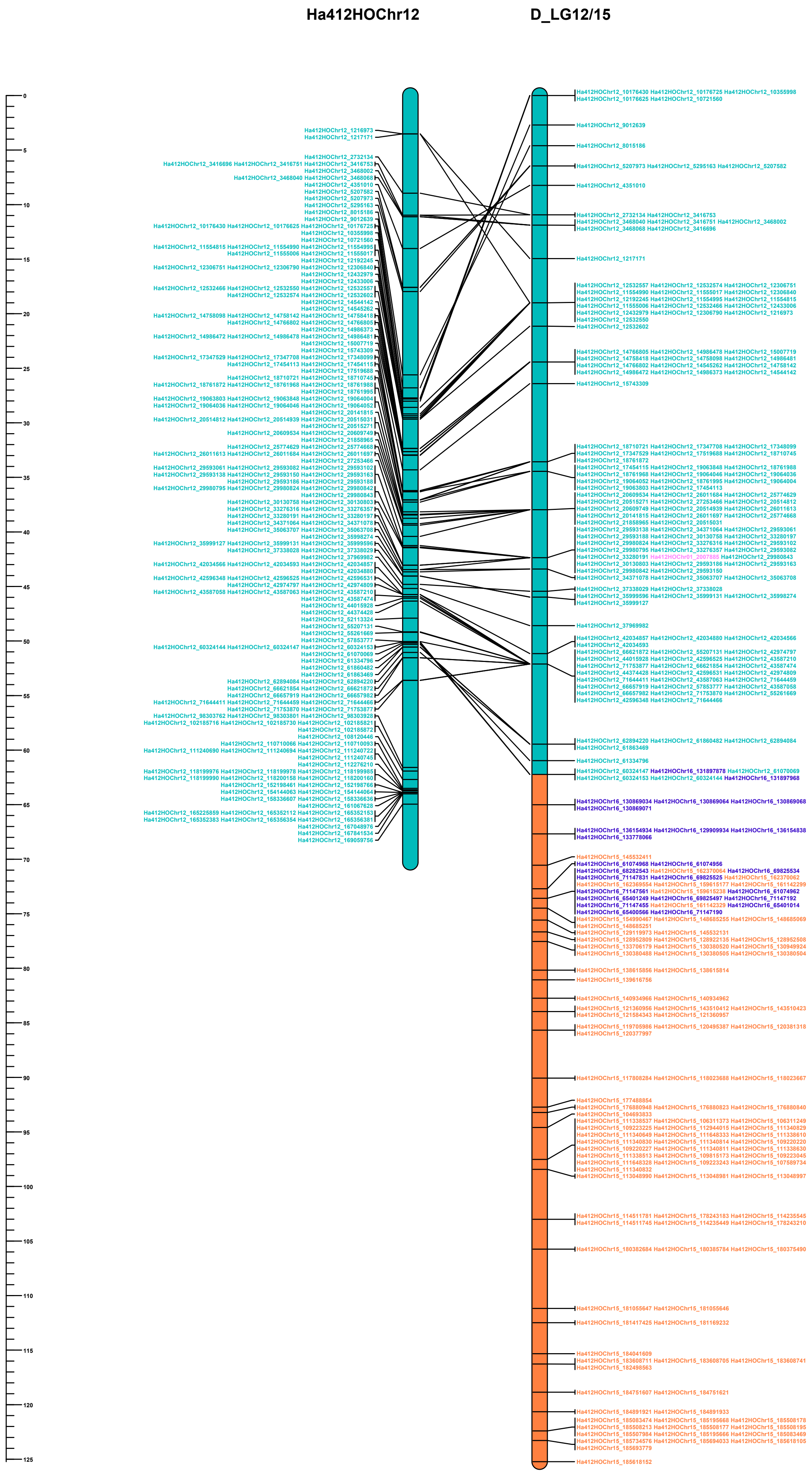

### HA412 vs Dune

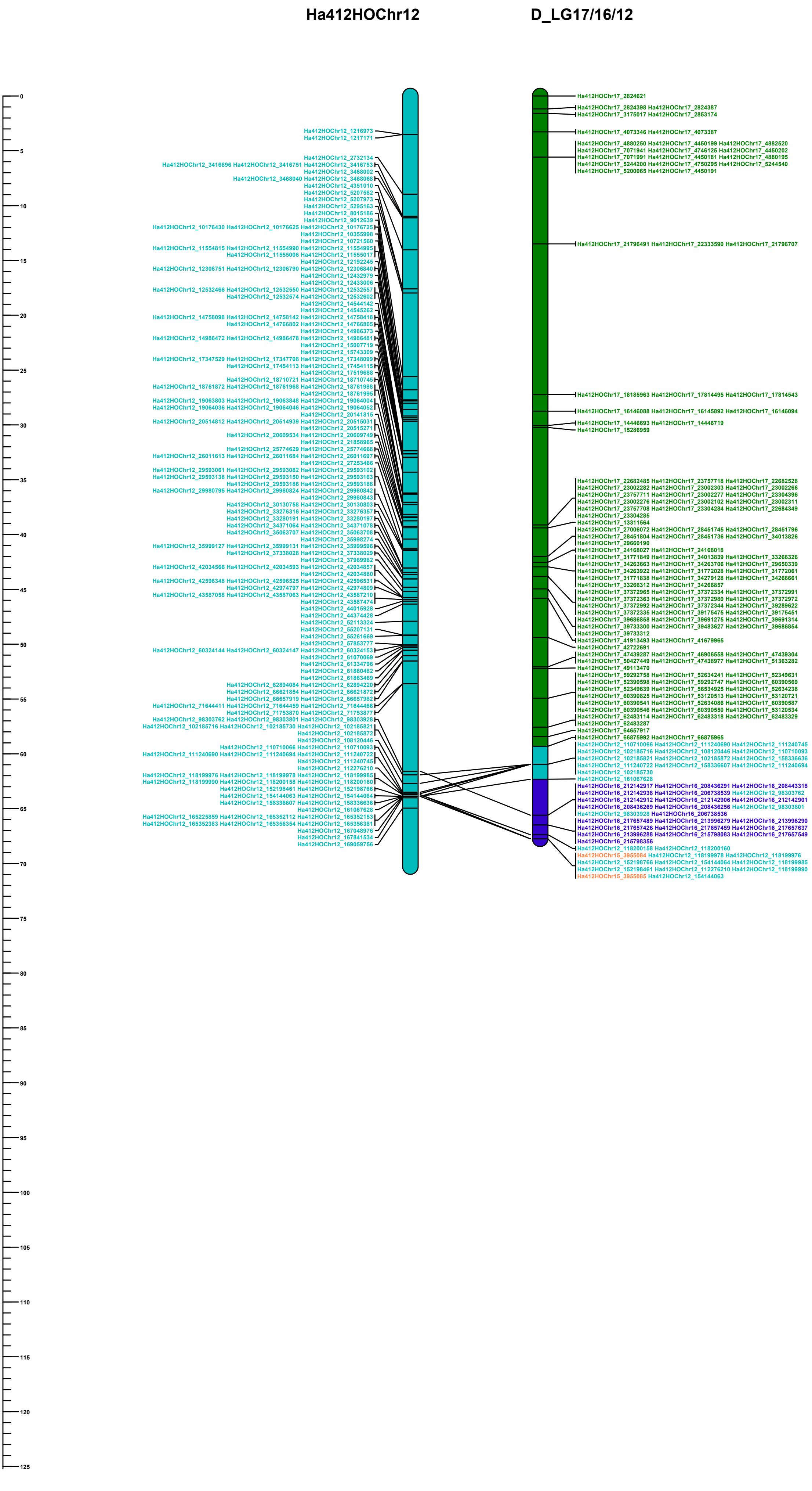

### HA412 vs Dune

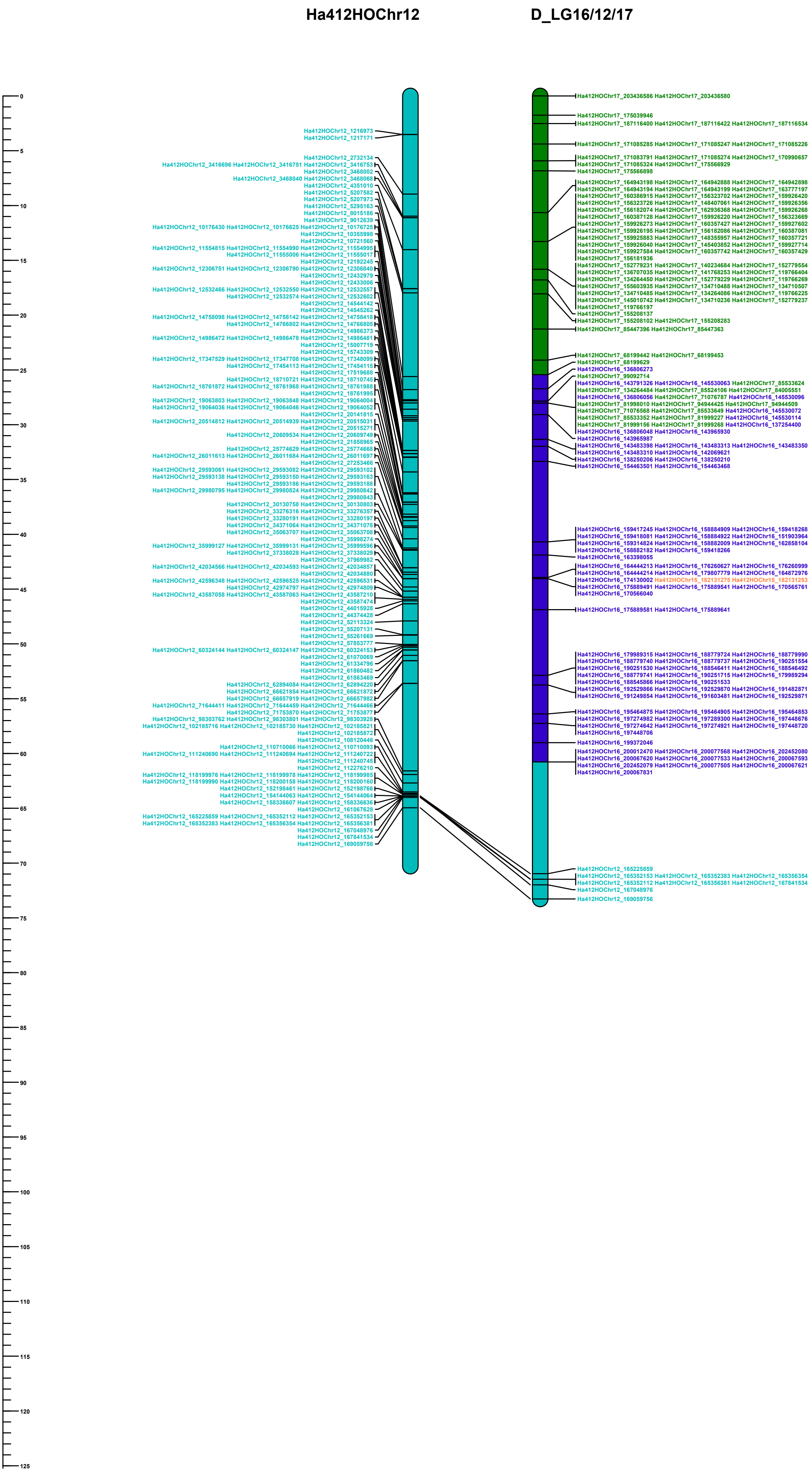

### HA412 vs Dune

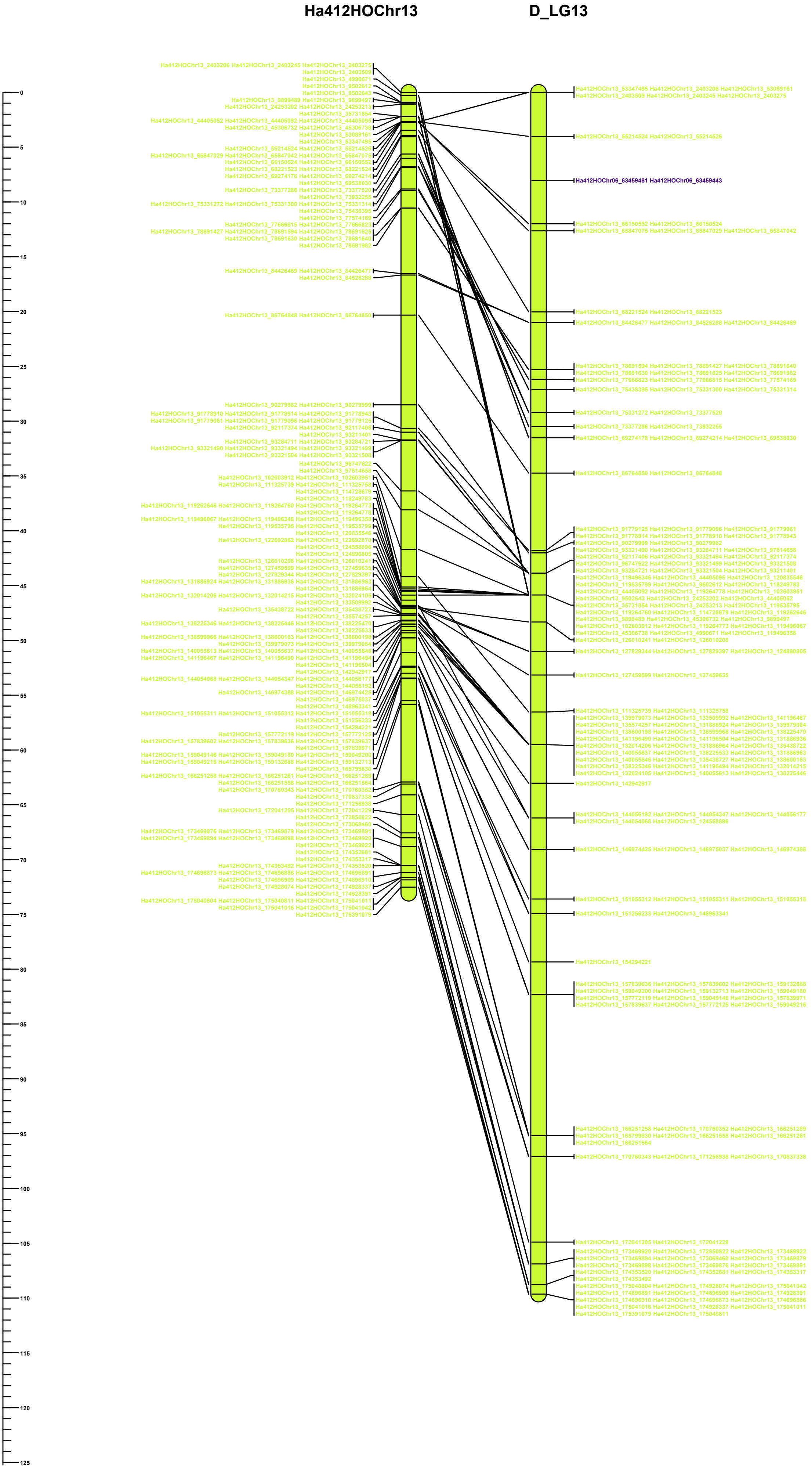

### HA412 vs Dune

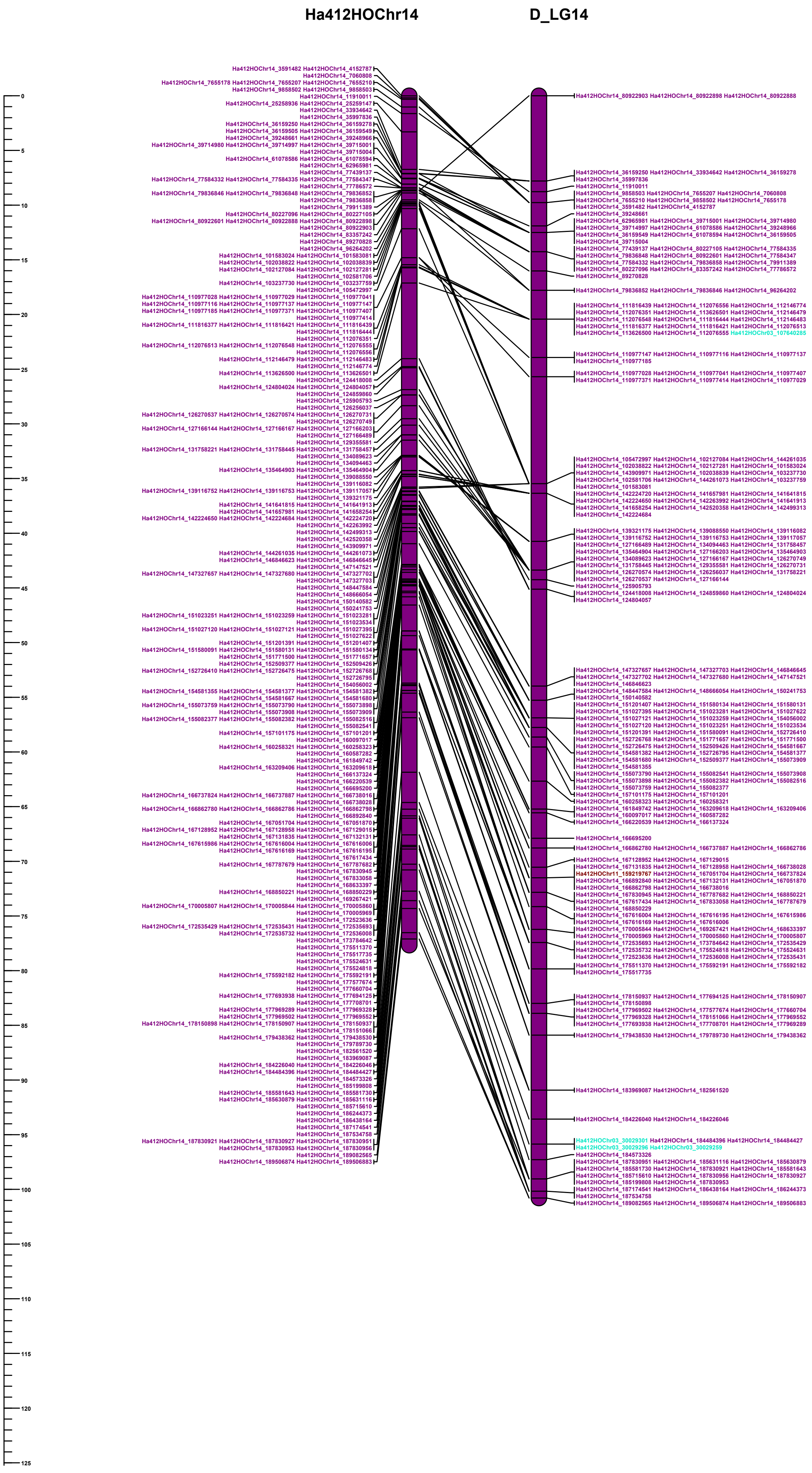

### HA412 vs Dune

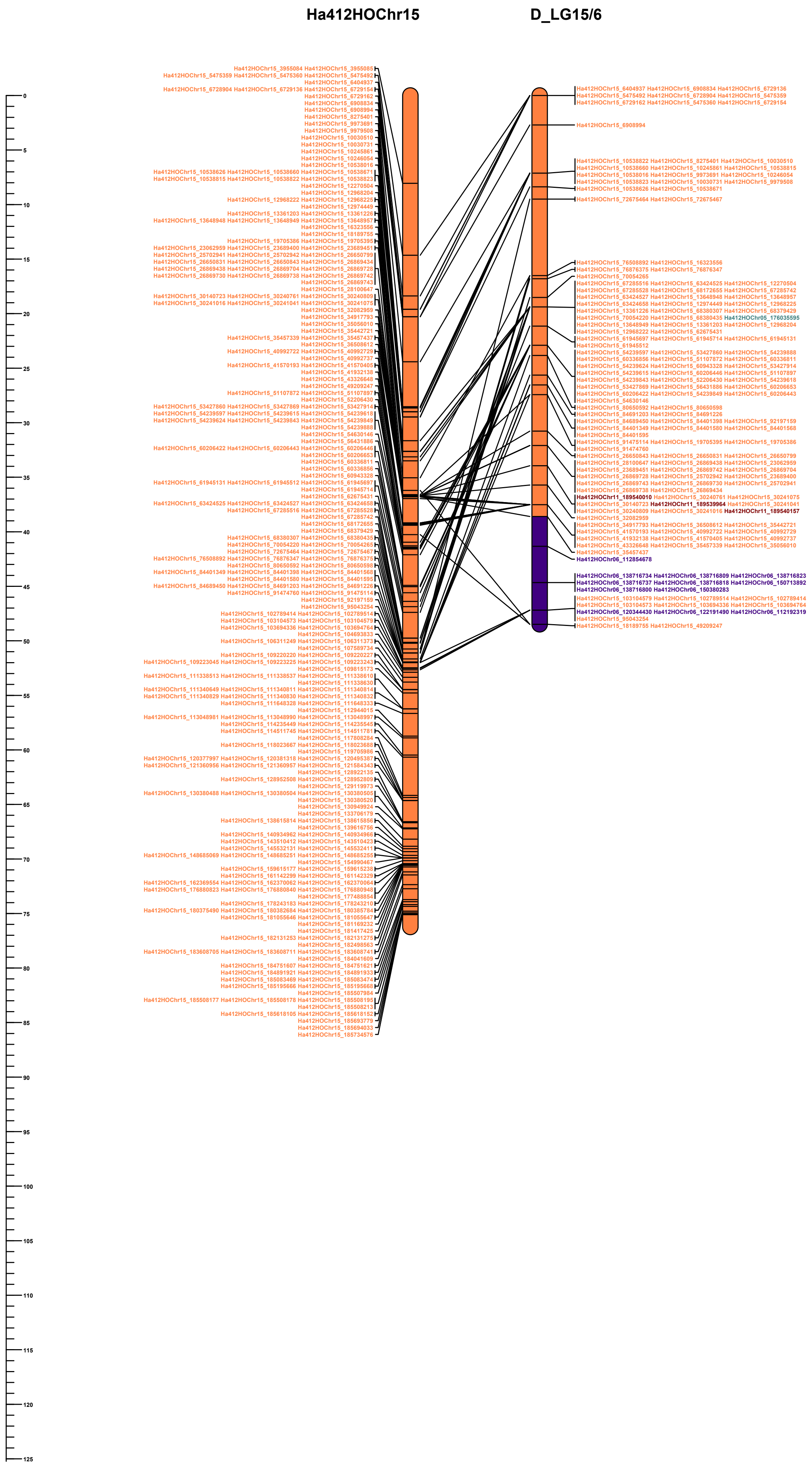

### HA412 vs Dune

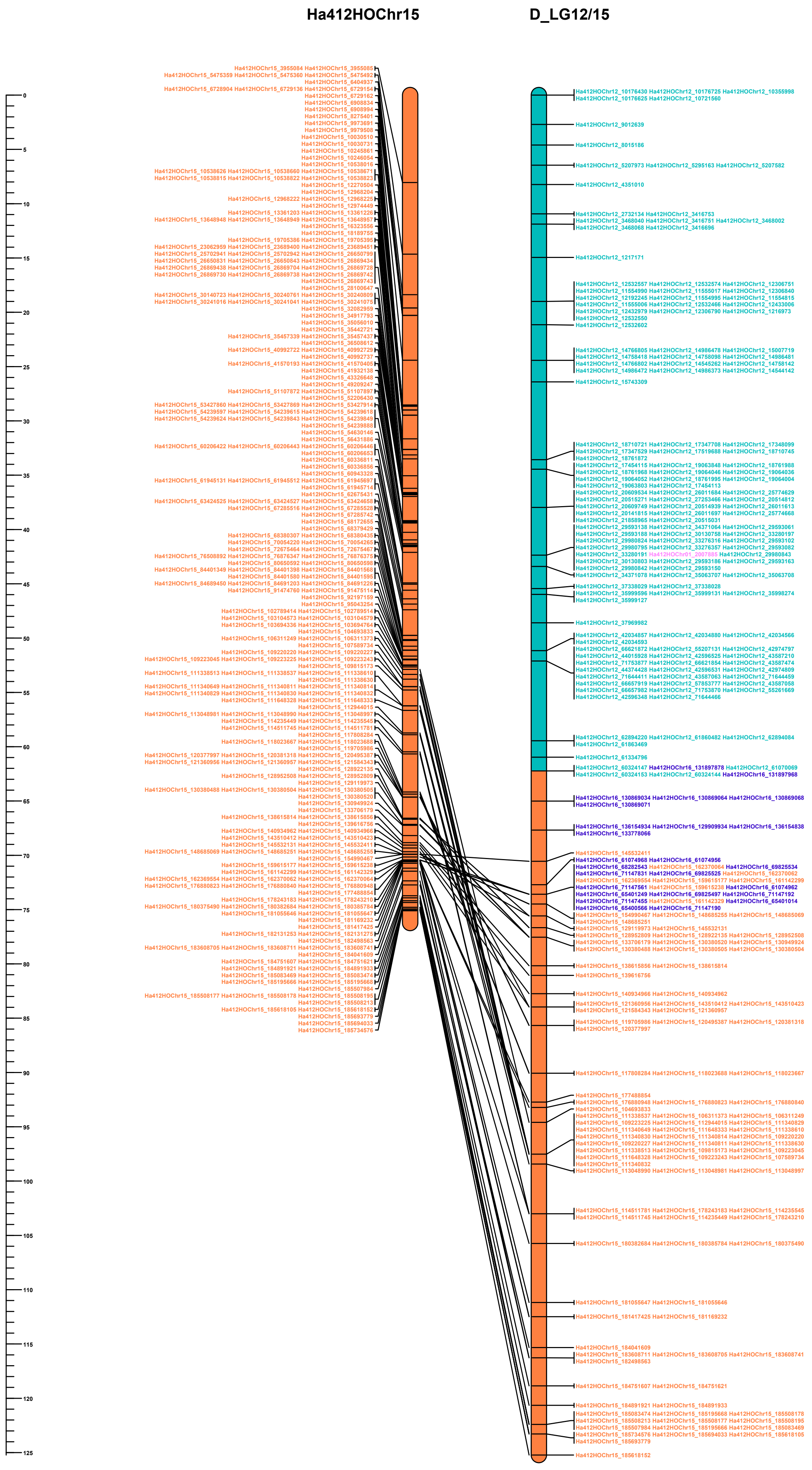

### HA412 vs Dune

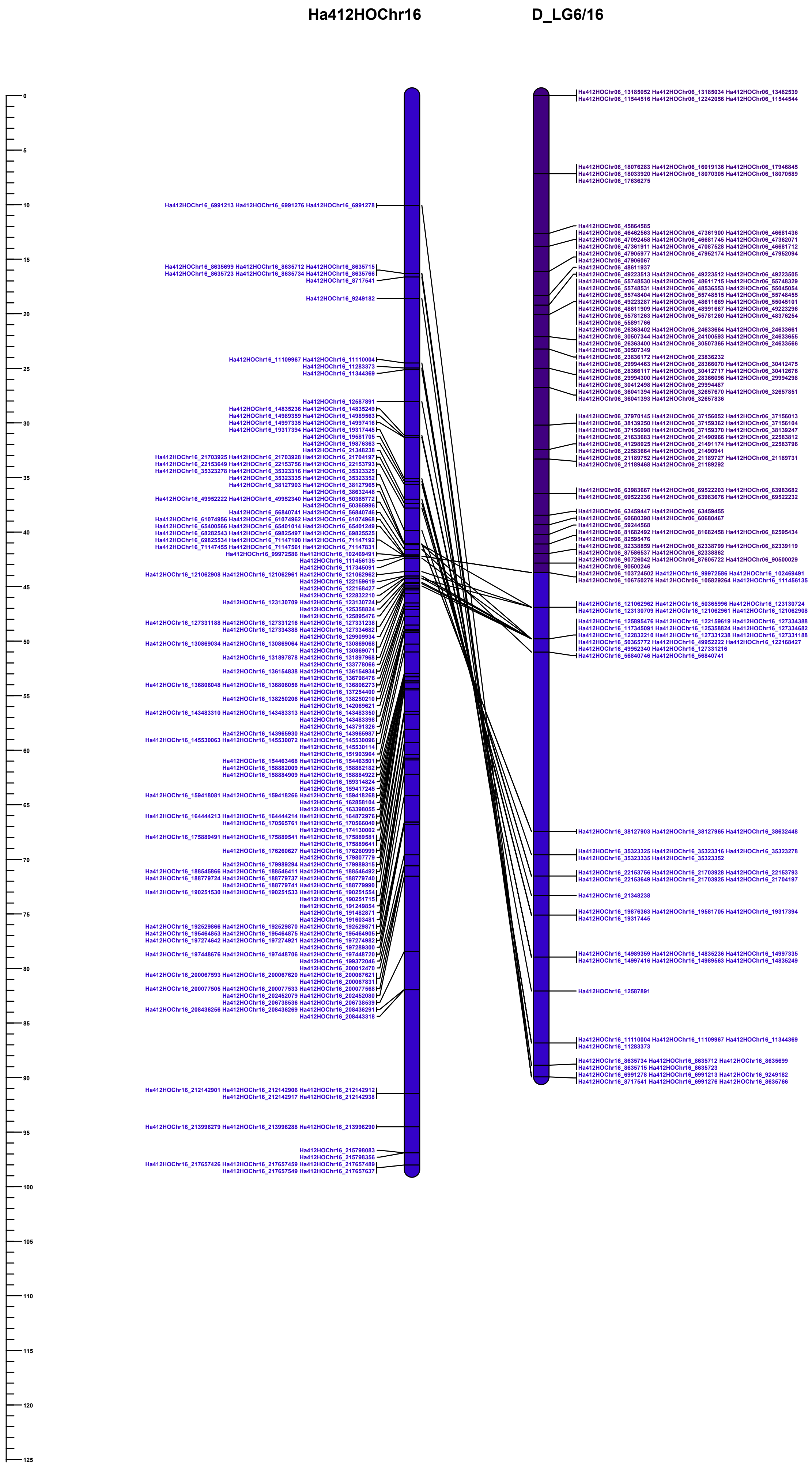

### HA412 vs Dune

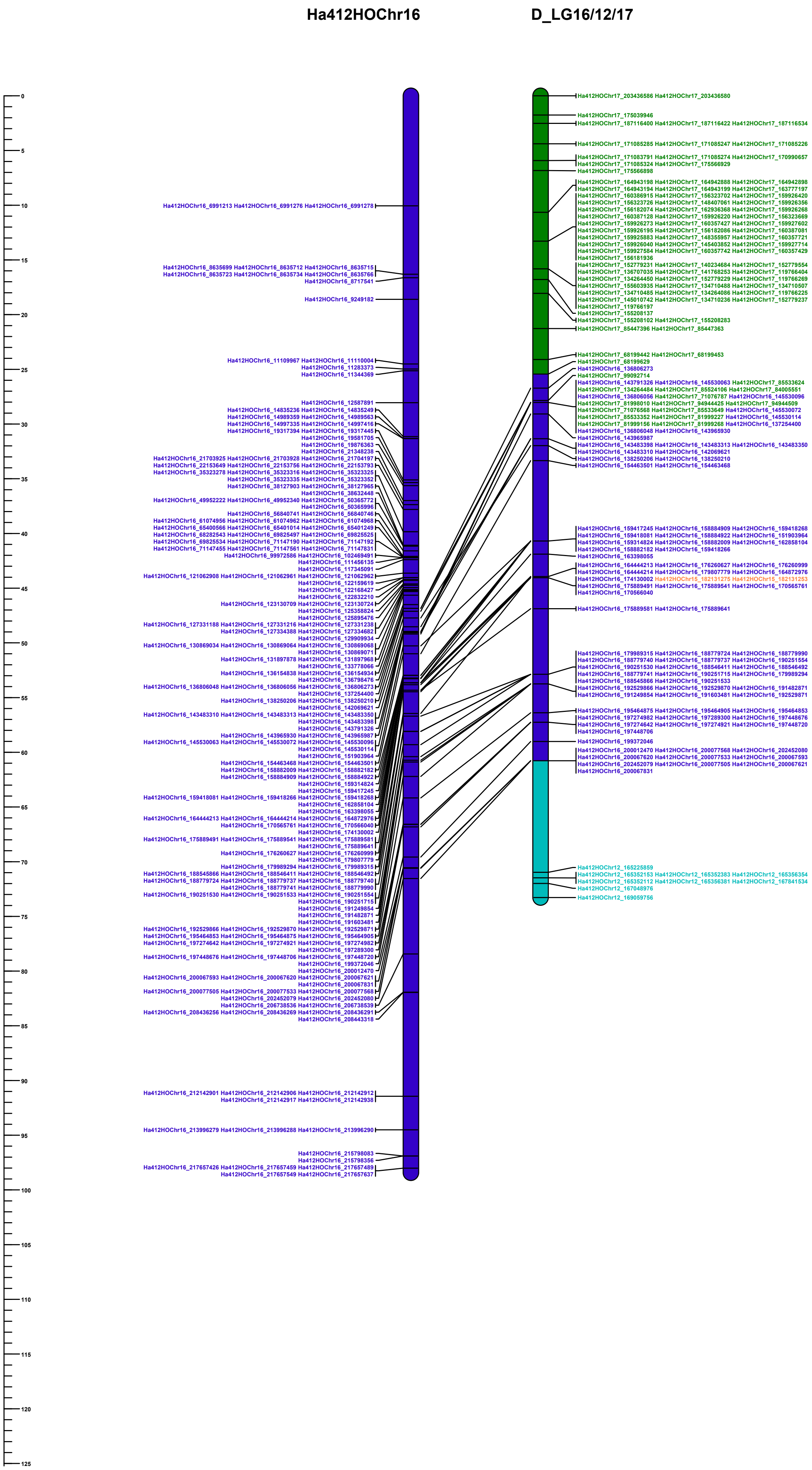

### HA412 vs Dune

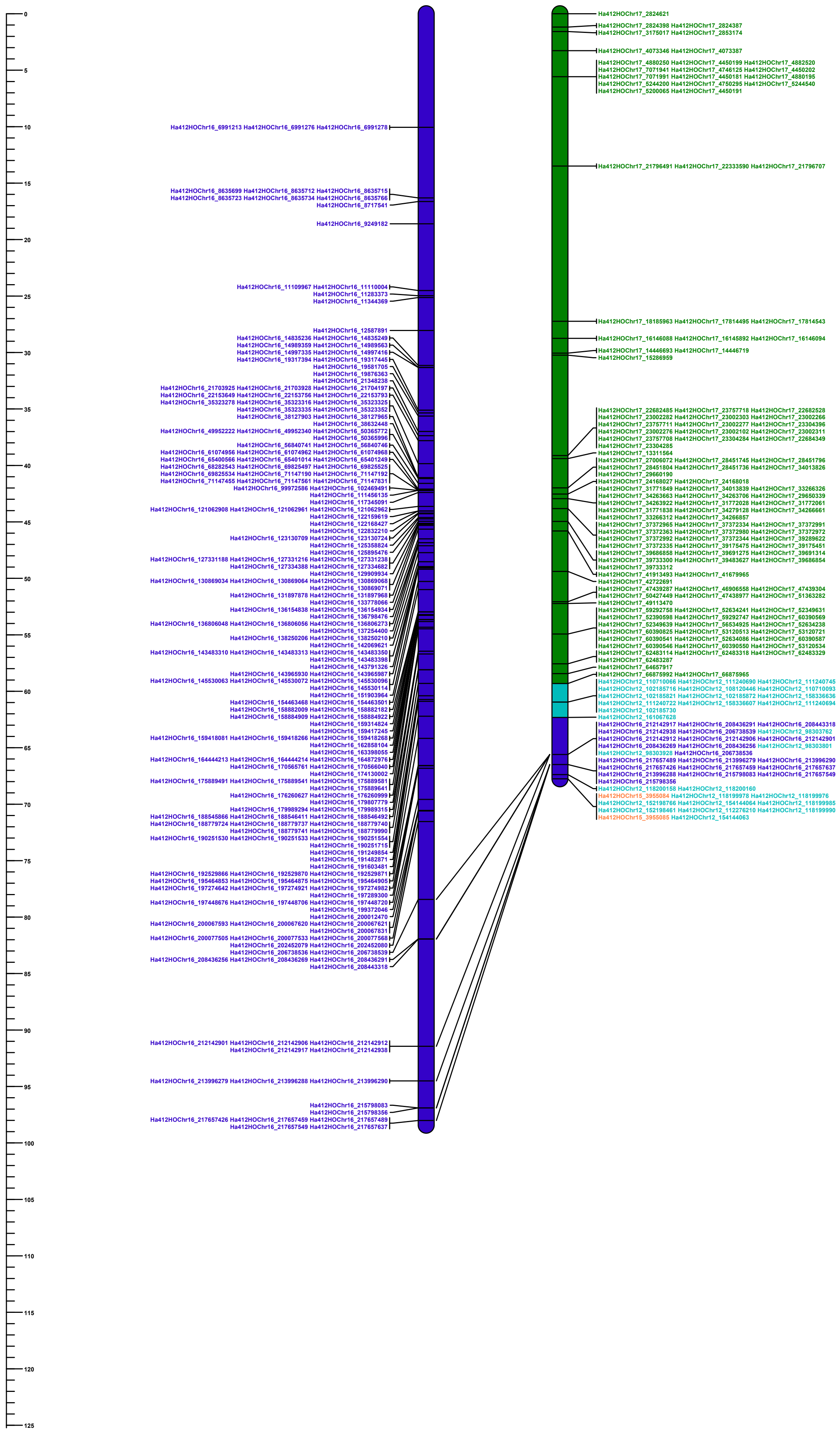

### HA412 vs Dune

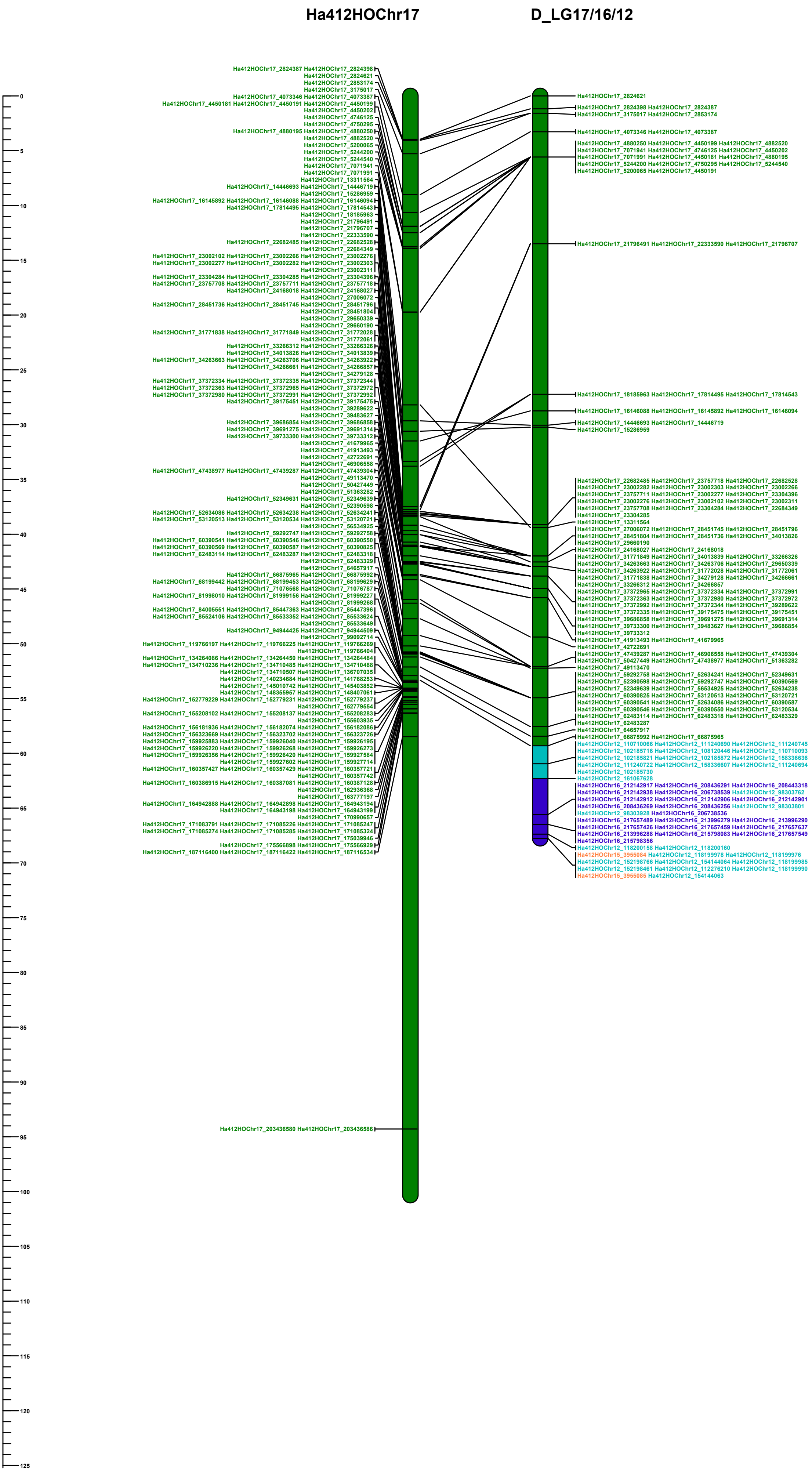

### HA412 vs Dune

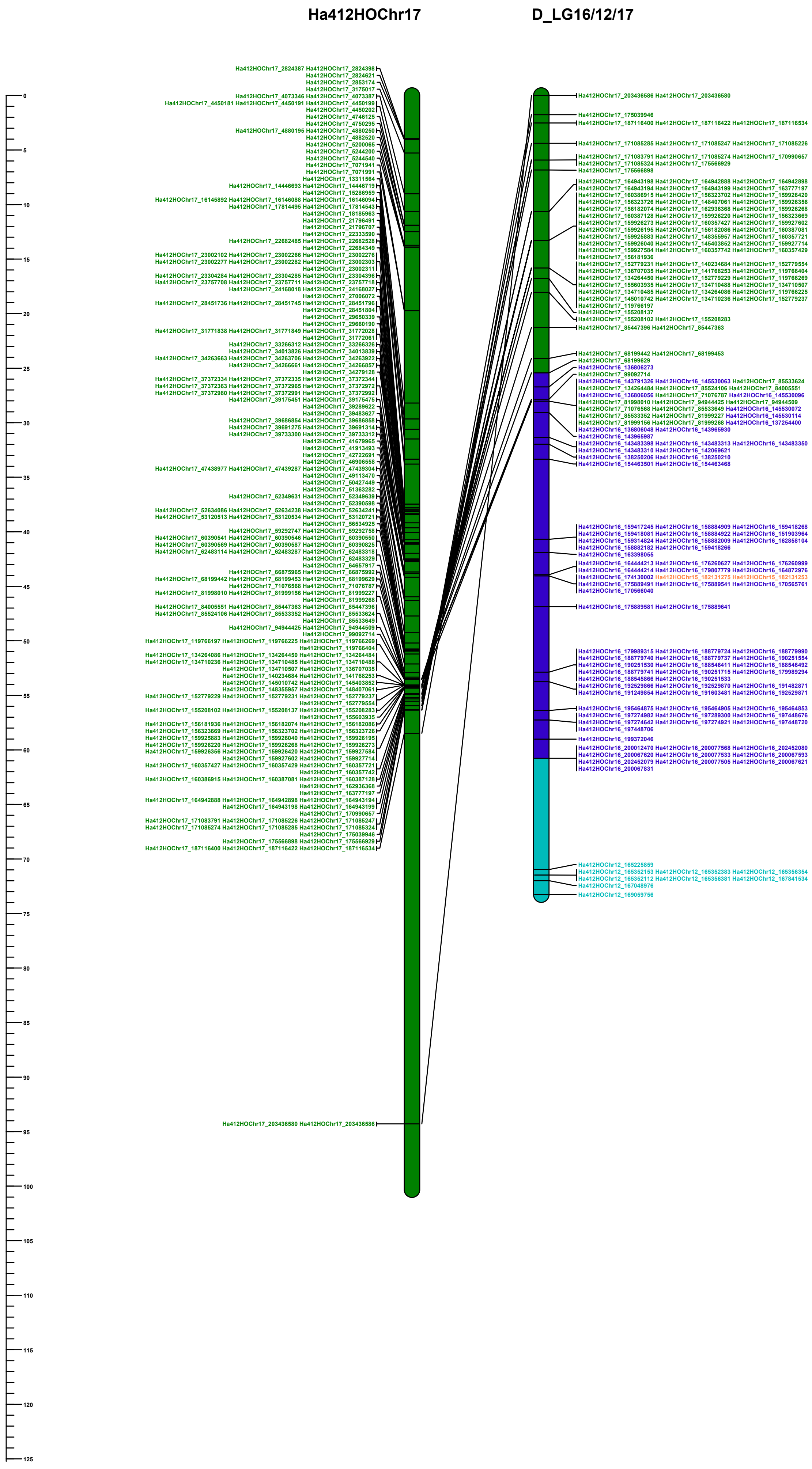
