## Supporting Information Figure S6 for "Multiple chromosomal inversions contribute to adaptive divergence of a dune sunflower ecotype"

**FIGURE S6** Genetic map comparison for CHR07.01, CHR09.02 and CHR11.01. Maps for non-dune (top panels) and dune (bottom panels) are plotted relative to the HA412HOv2 reference genome. Regions identified by lostruct and the markers that fall within them are highlighted in violet. CHR11.01 showed reduced recombination in the non-dune map, but part of this region had markers in reverse order compared to the reference in the map made from dune plant. Marker order is the same between the ecotypes for CHR07.01 and CHR09.02.

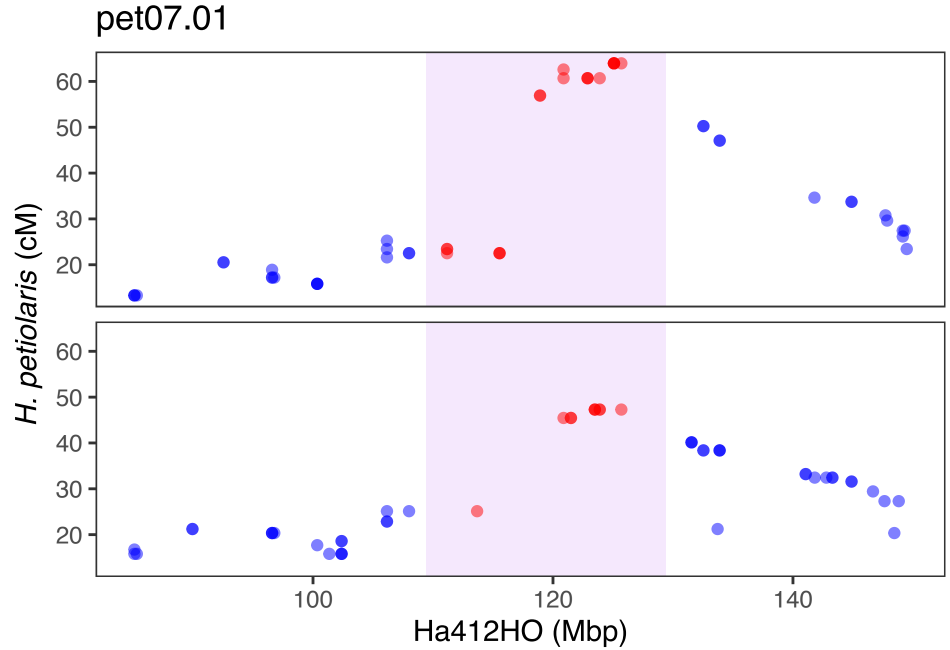
pet07.01
H. petiolaris (cM)
60
50
40
30
20
100
120
140
Ha412HO (Mbp)

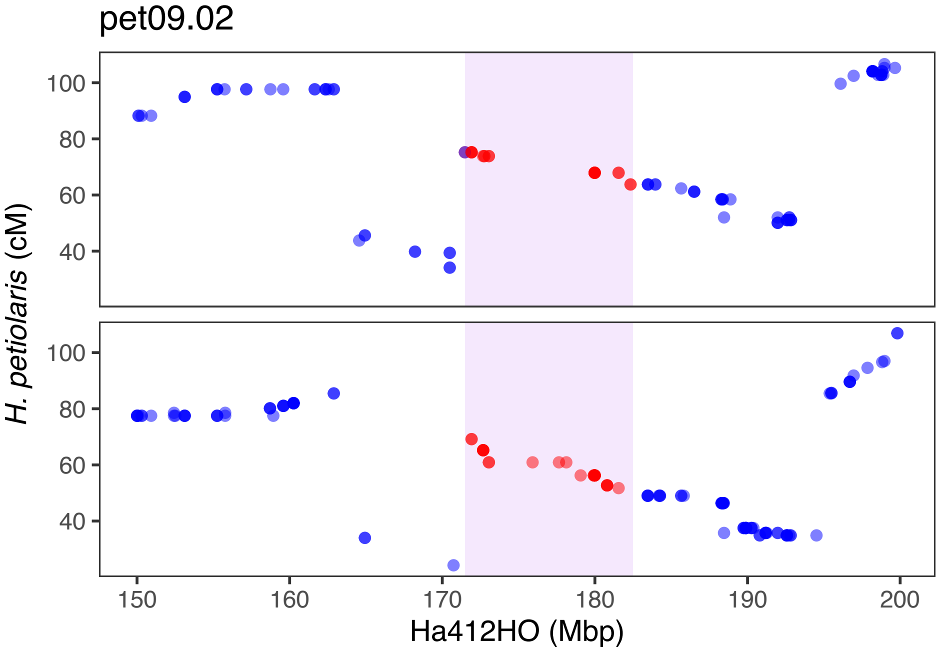

pet09.02
100
80
60
40
H. petiolaris (cM)
150
160
170
180
190
200
Ha412HO (Mbp)

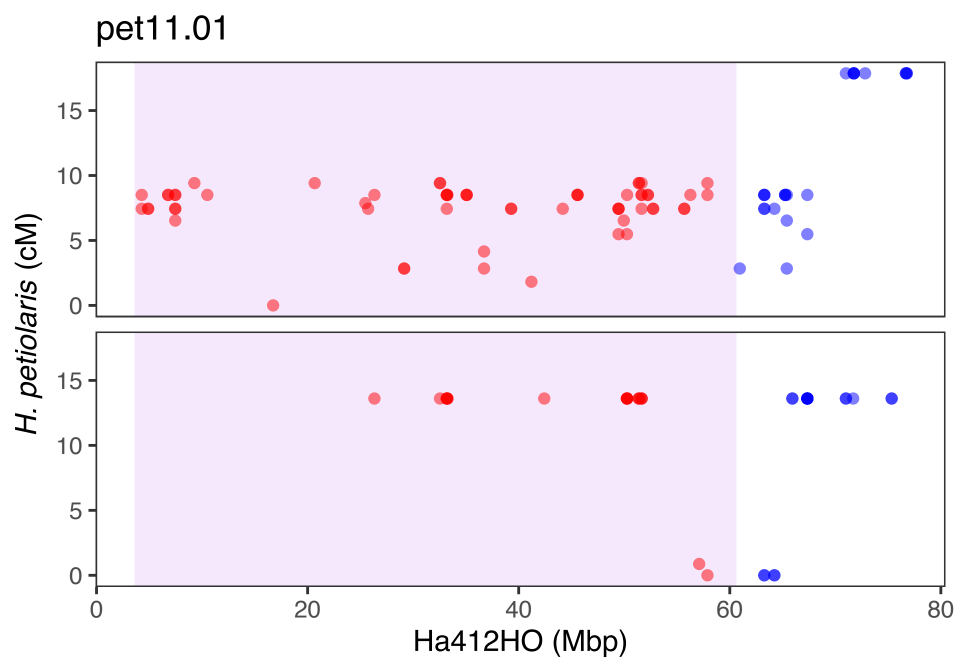
pet11.01
H. petiolaris (cM)
Ha412HO (Mbp)
0
5
10
15
0
20
40
60
80
