## Supporting Information Figure S7 for "Multiple chromosomal inversions contribute to adaptive divergence of a dune sunflower ecotype"

**FIGURE S7** Genome-environment association for coverage variables. Bayes factors ($BF_{is}$, in deciban unit) was estimated using importance sampling estimator approach in BayPass. SNPs on different reference chromosomes are represented in alternate grey colors. The locations and $BF_{is}$ values of 7 putative inversions are indicated by red solid bars. Red horizontal dashed lines represent 1% significance thresholds computed from simulated samples.

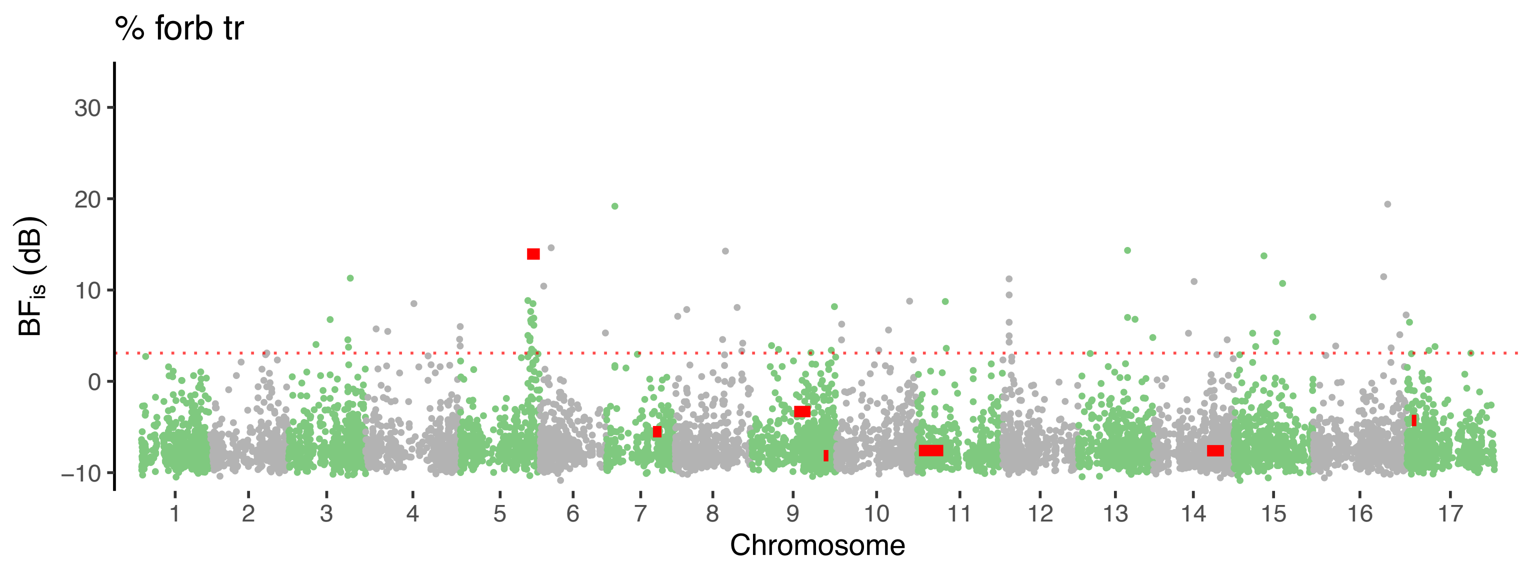

% forb tr
BF_is (dB)
30
20
10
0
−10
Chromosome

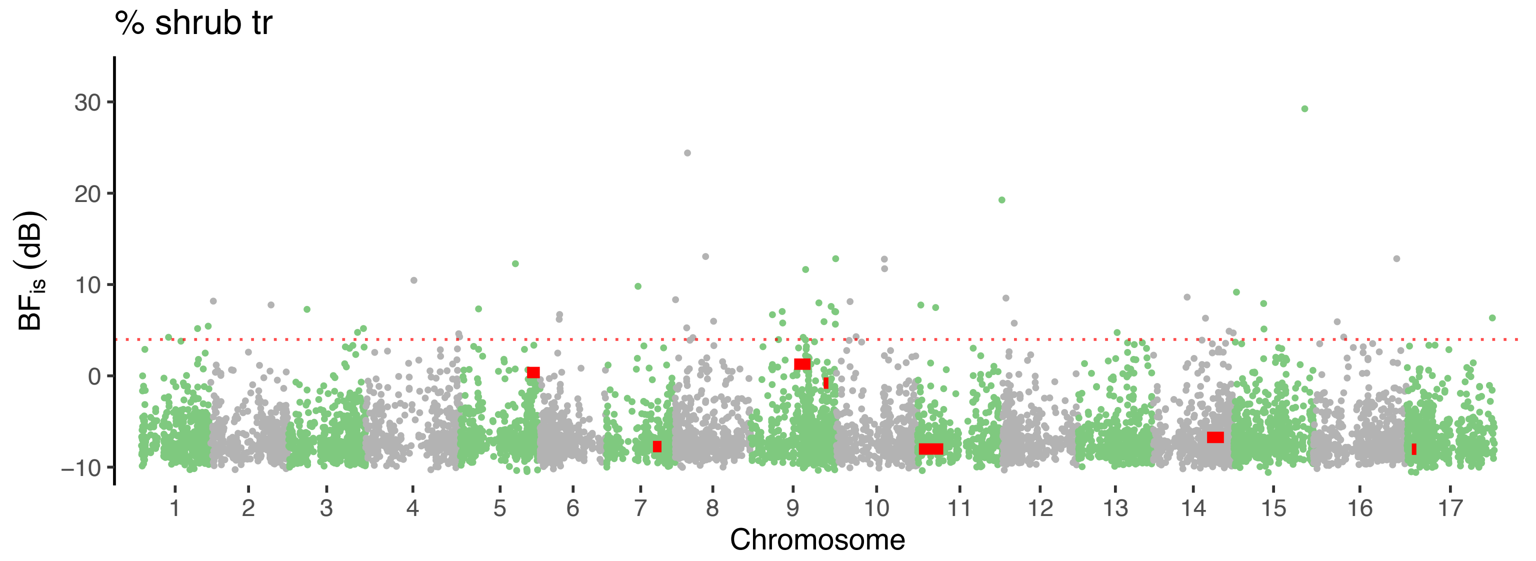

% shrub tr
BF$_{is}$ (dB)
30
20
10
0
−10
1 2 3 4 5 6 7 8 9 10 11 12 13 14 15 16 17
Chromosome

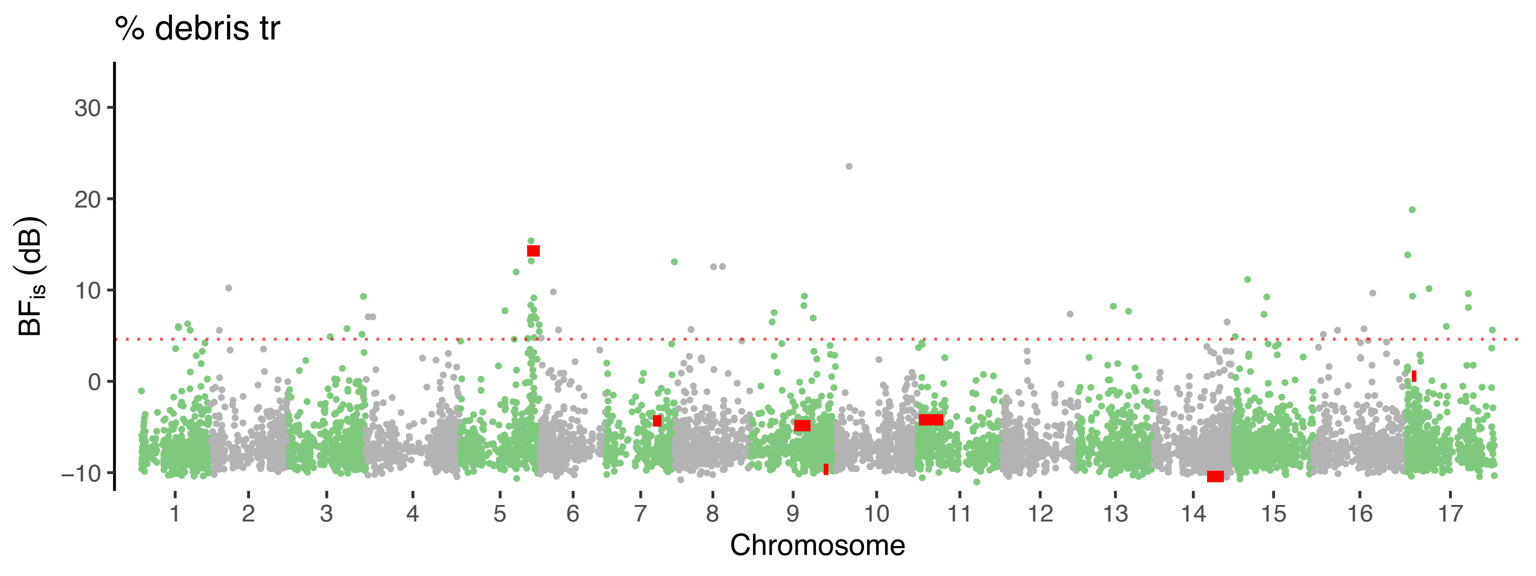

% debris tr
BFis (dB)
30
20
10
0
−10
Chromosome

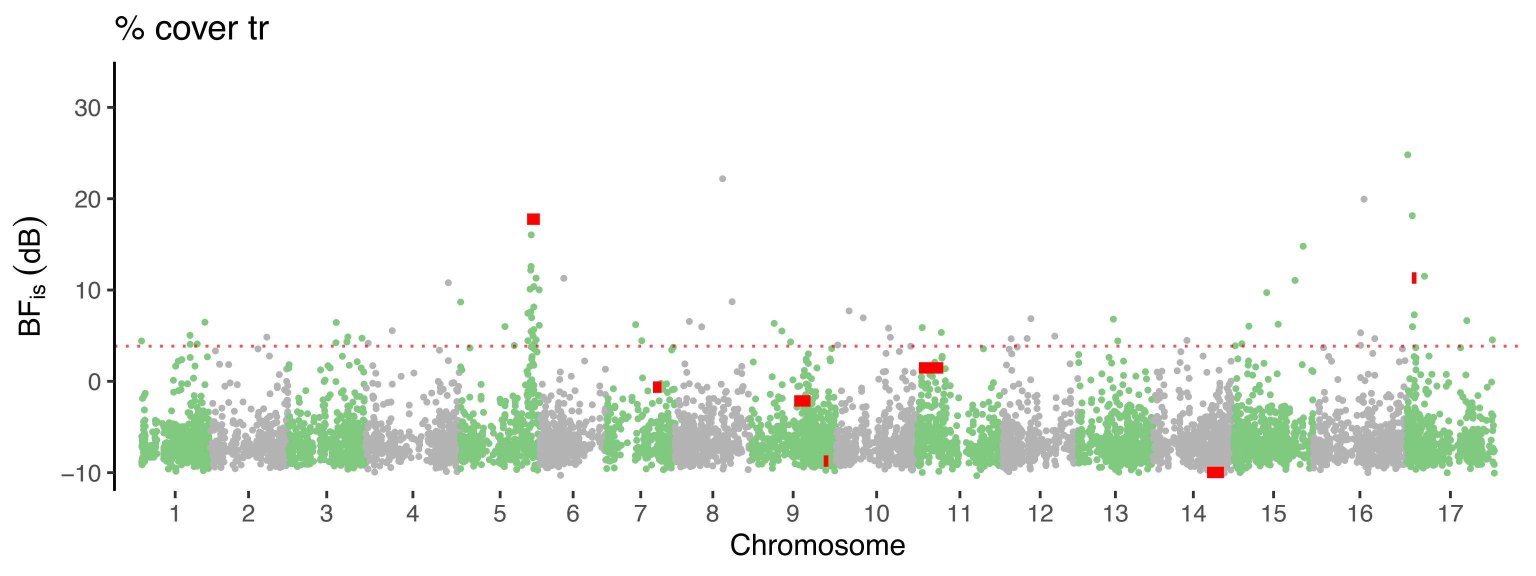
% cover tr
30
20
10
0
−10
BFis (dB)
Chromosome

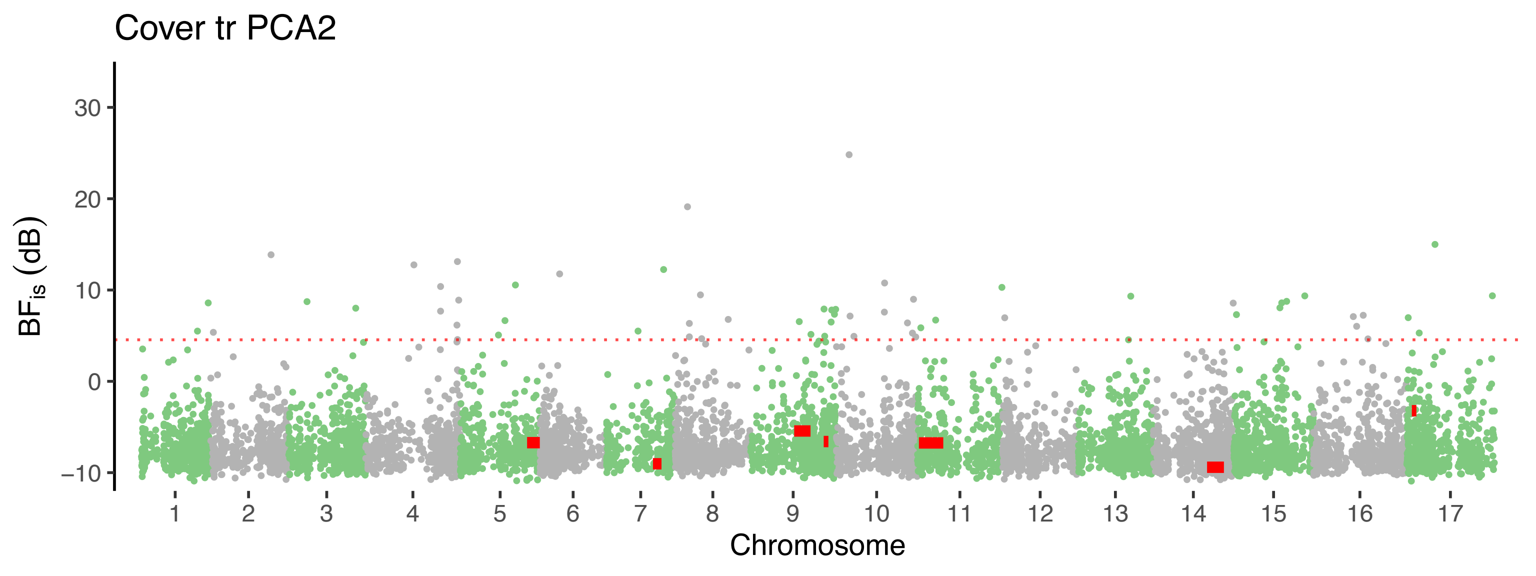
Cover tr PCA2
30
20
10
0
−10
$BF_{is}$ (dB)
Chromosome

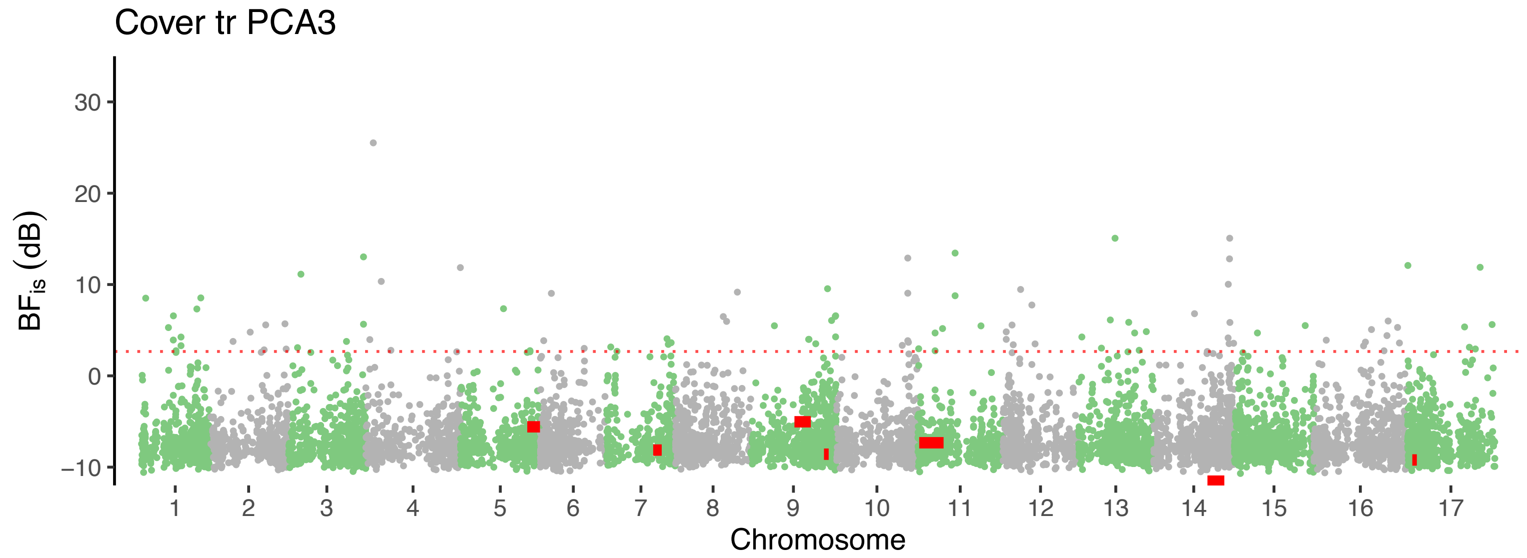

Cover tr PCA3
BFis (dB)
30
20
10
0
−10
Chromosome
