## Supporting Information Figure S8 for "Multiple chromosomal inversions contribute to adaptive divergence of a dune sunflower ecotype"

**FIGURE S8** Genome-environment association for soil variables. Bayes factors ($BF_{is}$, in deciban unit) was estimated using importance sampling estimator approach in BayPass. SNPs on different reference chromosomes are represented in alternate grey colors. The locations and $BF_{is}$ values of 7 putative inversions are indicated by red solid bars. Red horizontal dashed lines represent 1% significance thresholds computed from simulated samples.

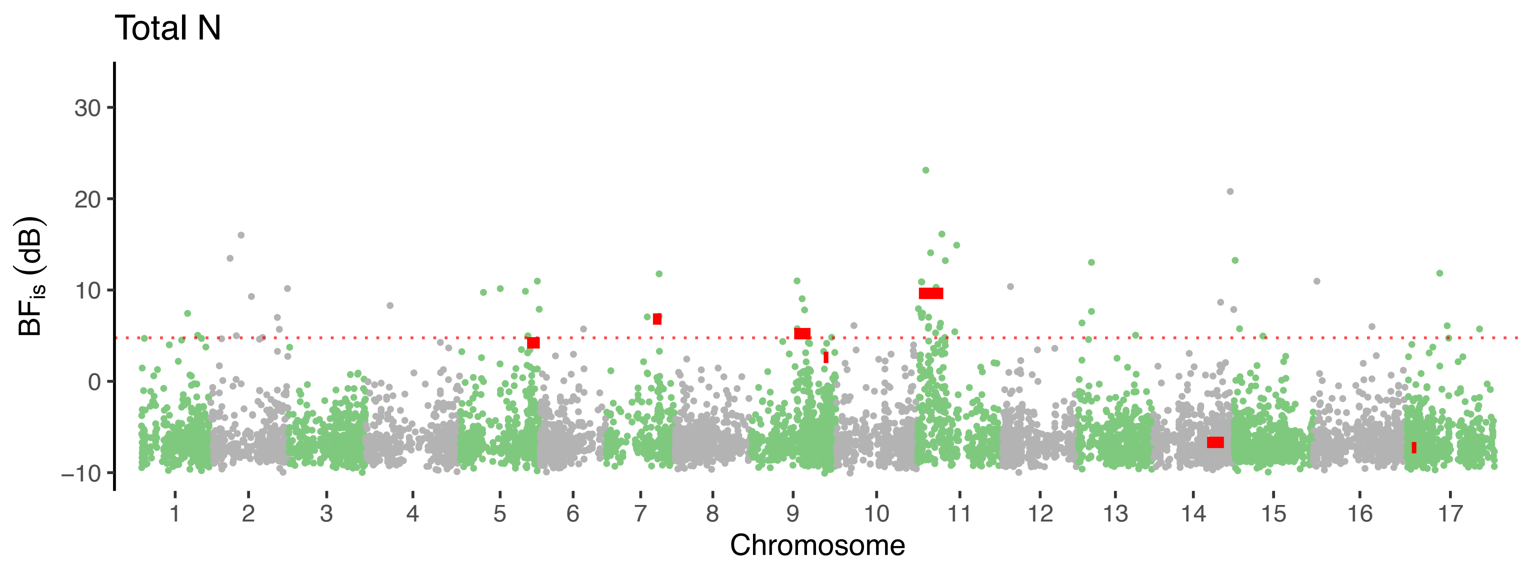
Total N
BFis (dB)
30
20
10
0
−10
Chromosome

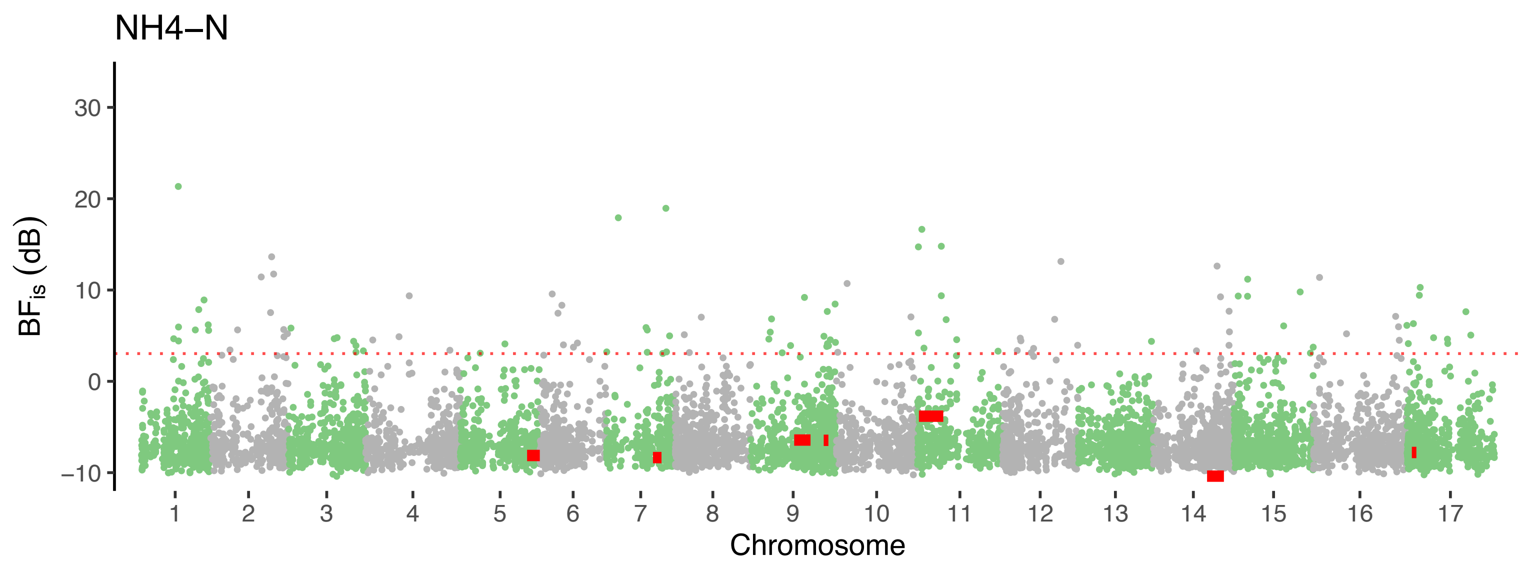

NH4−N
BFis (dB)
30
20
10
0
−10
Chromosome

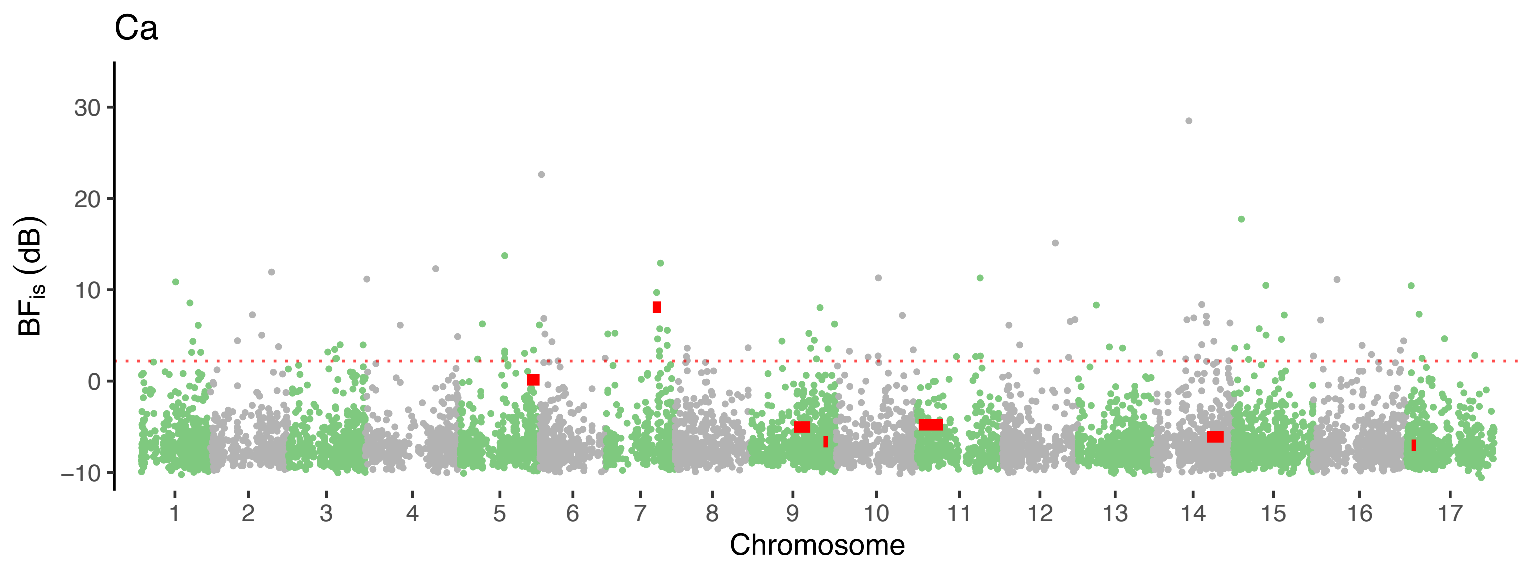

Ca
30
20
10
0
−10
BF_is (dB)
Chromosome

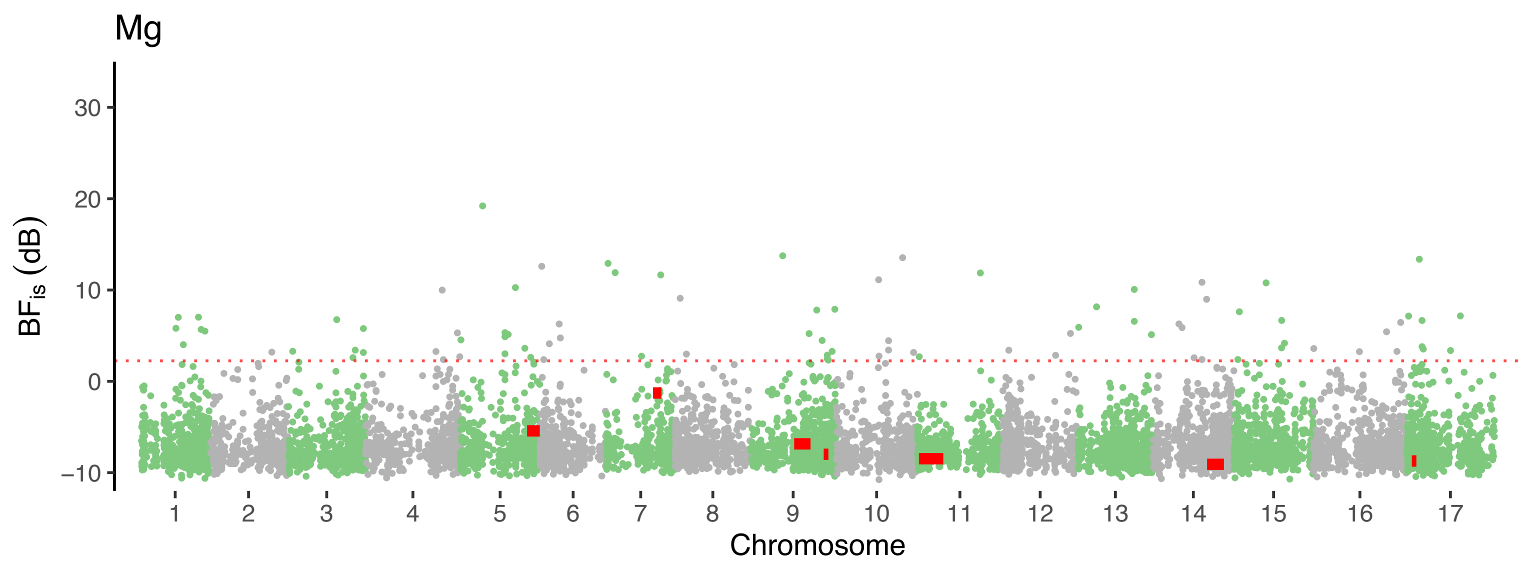

Mg
BF$_{is}$ (dB)
30
20
10
0
−10
1 2 3 4 5 6 7 8 9 10 11 12 13 14 15 16 17
Chromosome

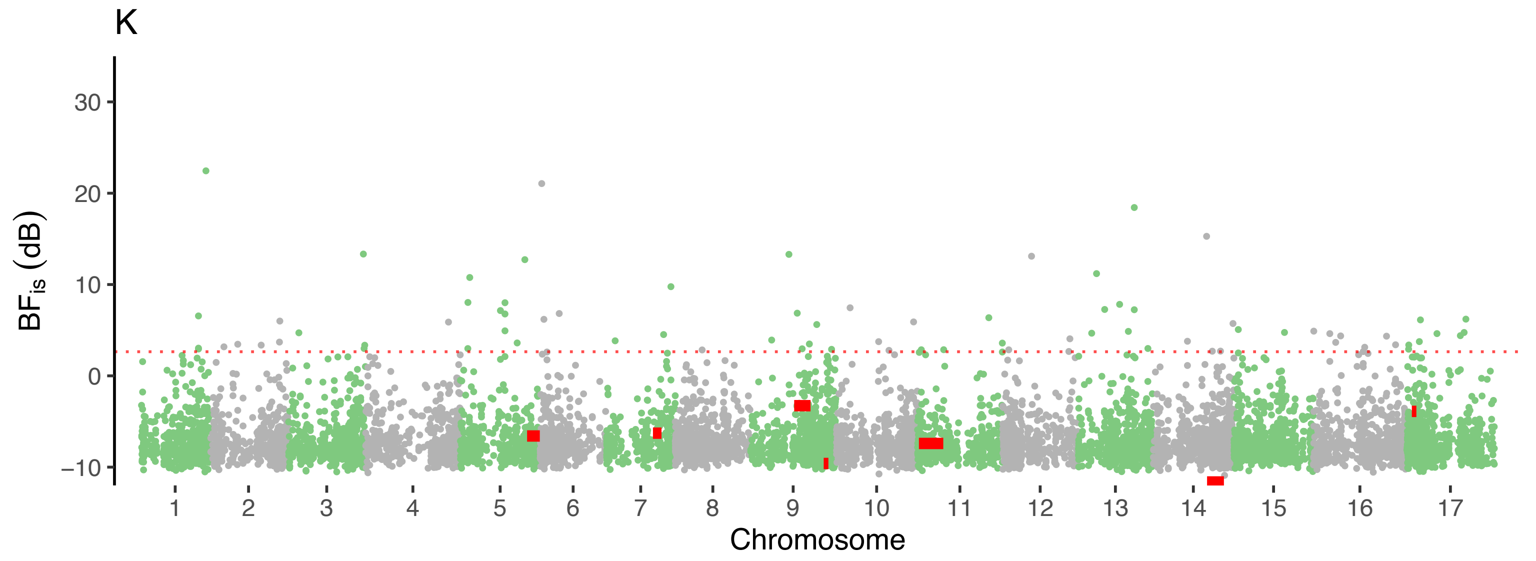
K
30
20
10
0
−10
BFis (dB)
Chromosome

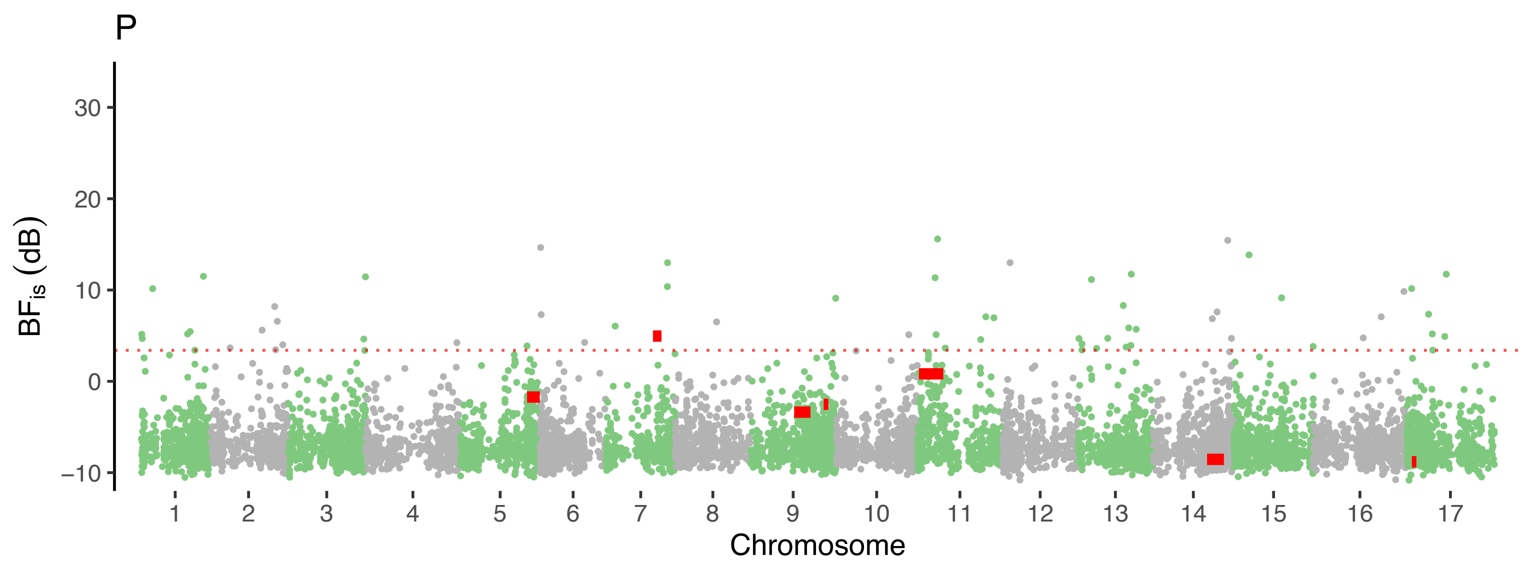
P
30
20
10
0
−10
$BF_{is}$ (dB)
Chromosome

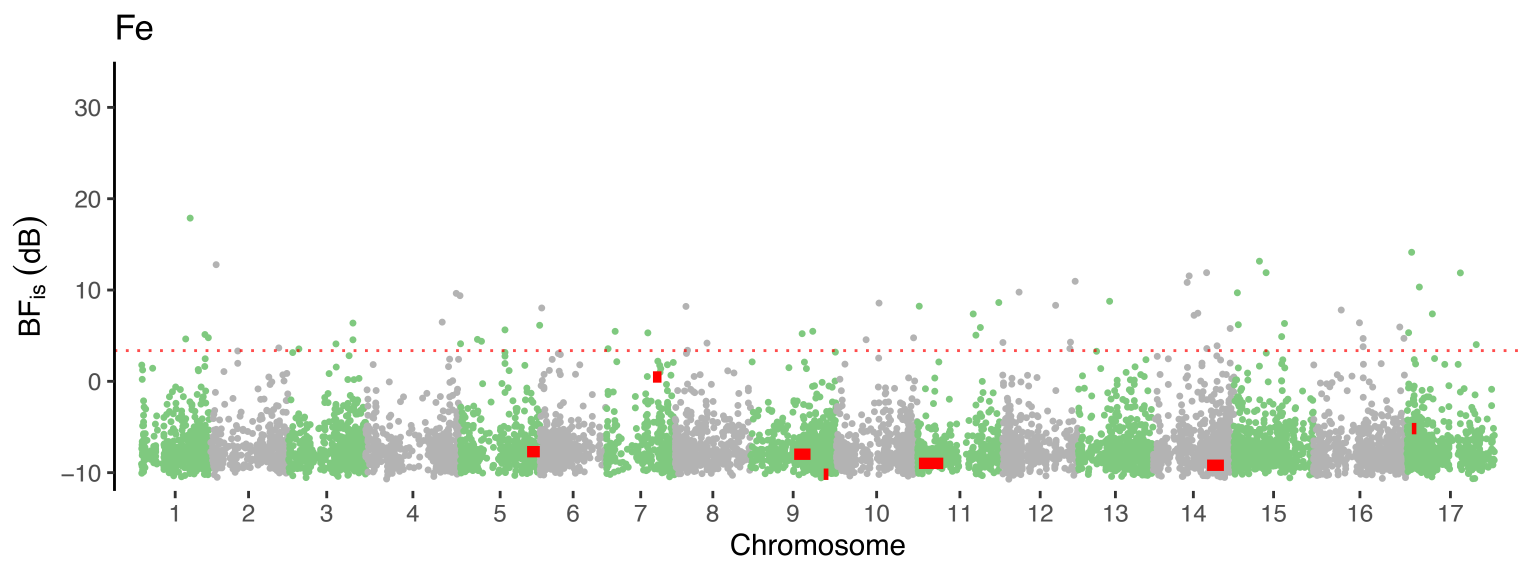

Fe
BF$_{is}$ (dB)
30
20
10
0
−10
Chromosome

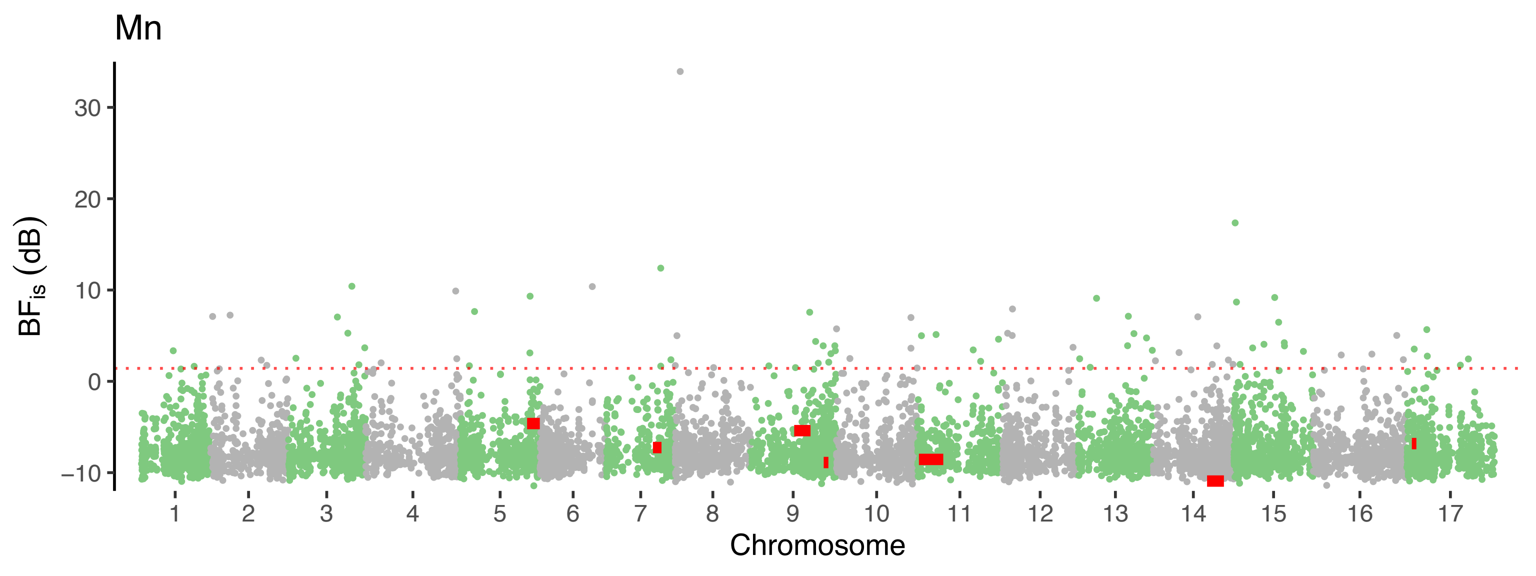

Mn
BFis (dB)
30
20
10
0
−10
Chromosome

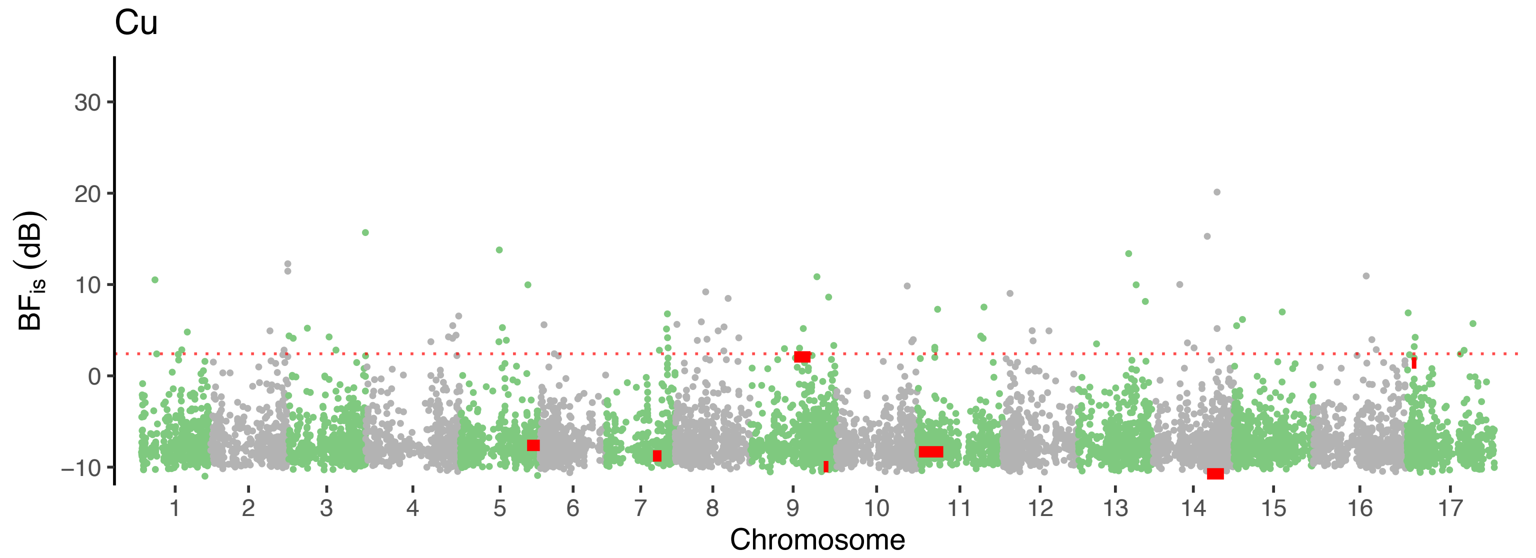
Cu
BFis (dB)
30
20
10
0
−10
Chromosome

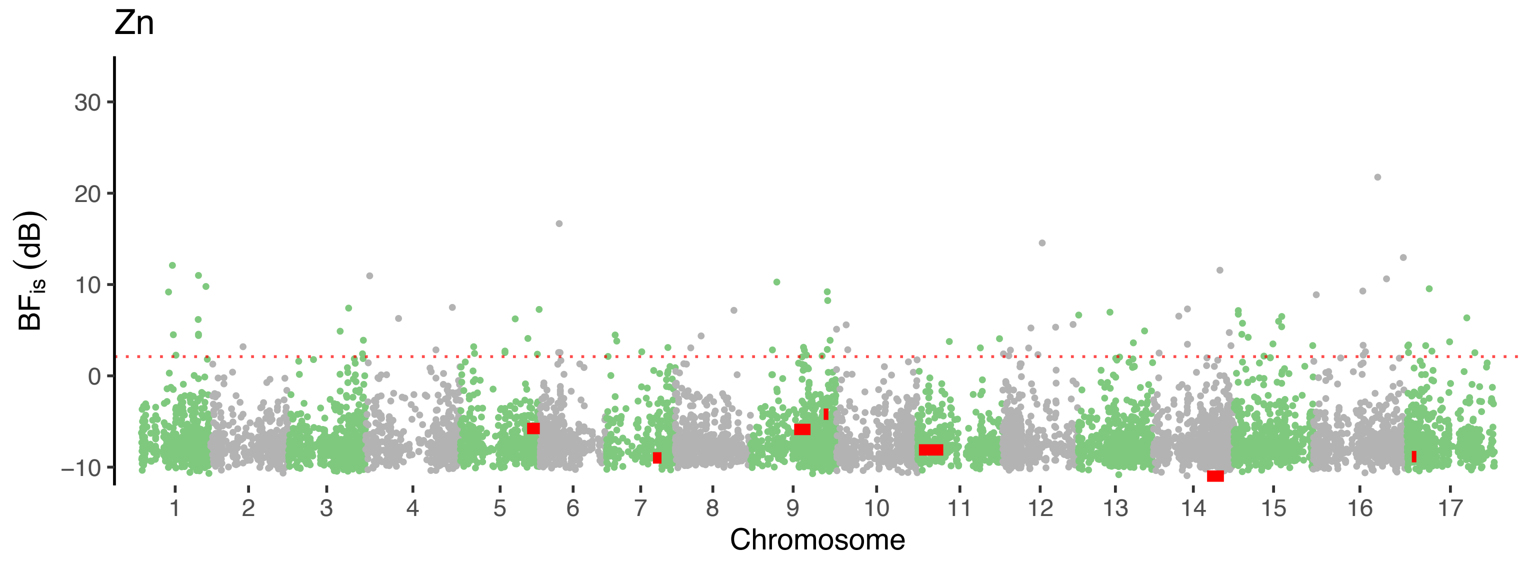
Zn
30
20
10
0
−10
BF_is (dB)
Chromosome

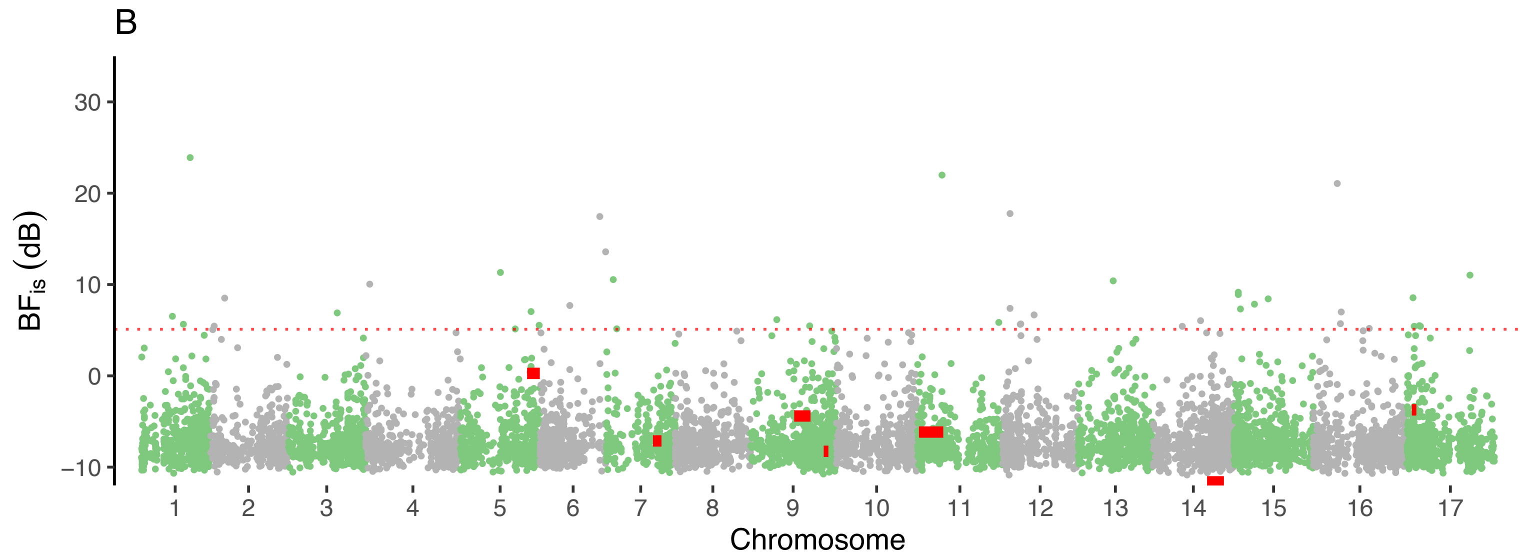

B
$BF_{is}$ (dB)
30
20
10
0
−10
Chromosome

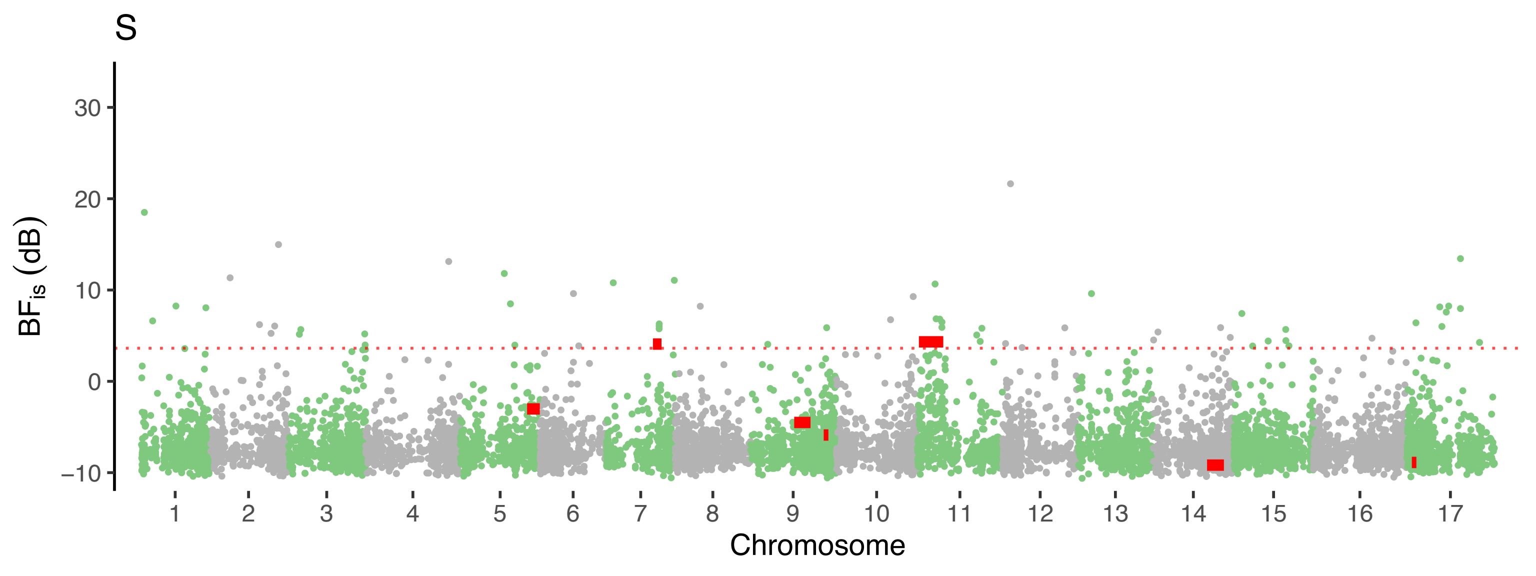

S
BF$_{is}$ (dB)
30
20
10
0
−10
Chromosome

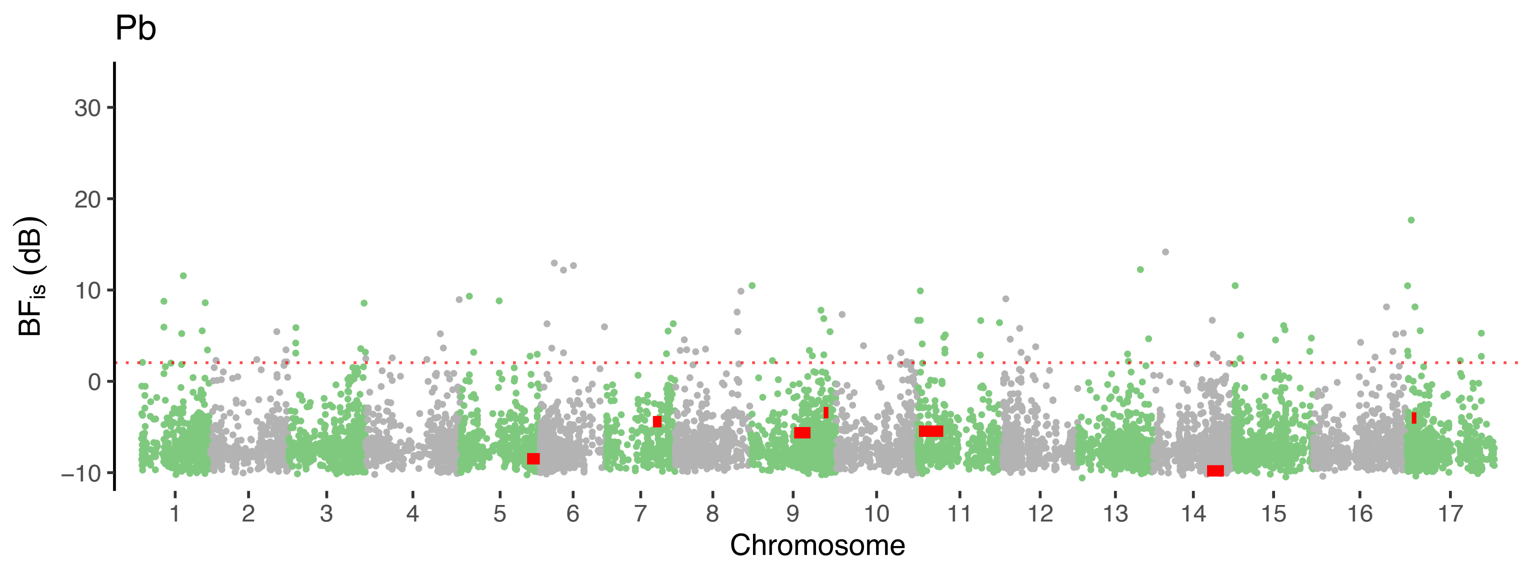

Pb
BF$_{is}$ (dB)
30
20
10
0
−10
Chromosome

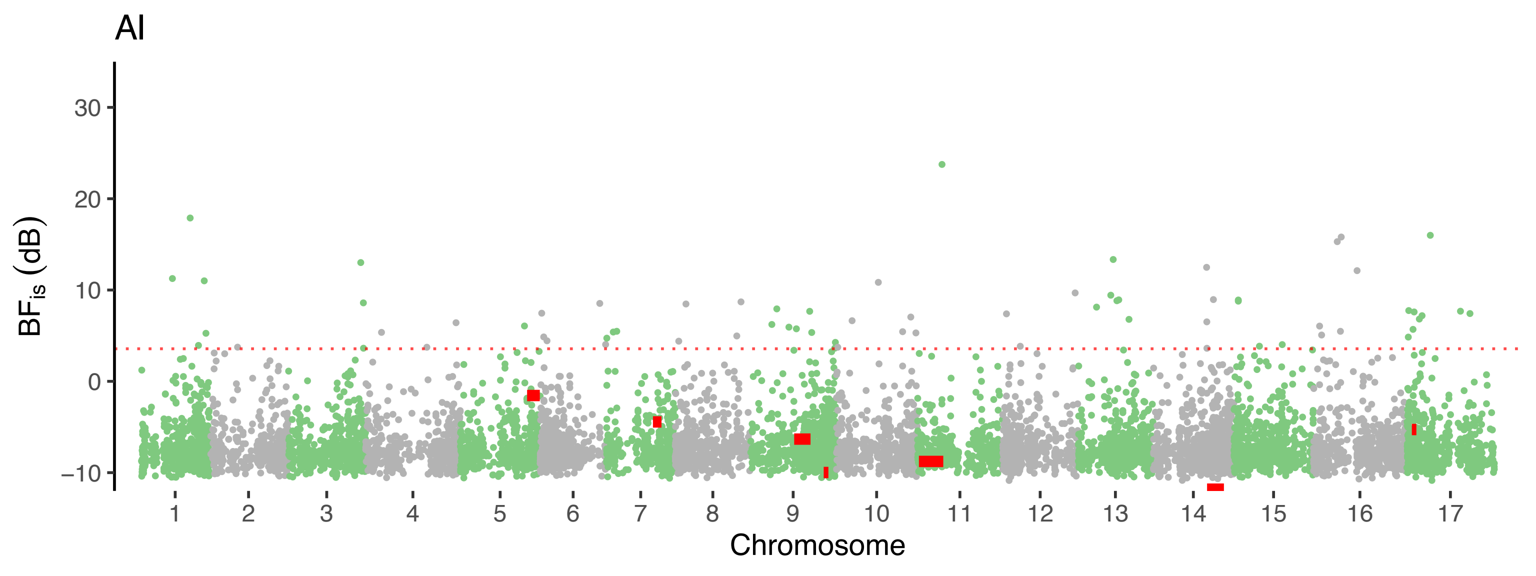
AI
BFis (dB)
30
20
10
0
−10
Chromosome

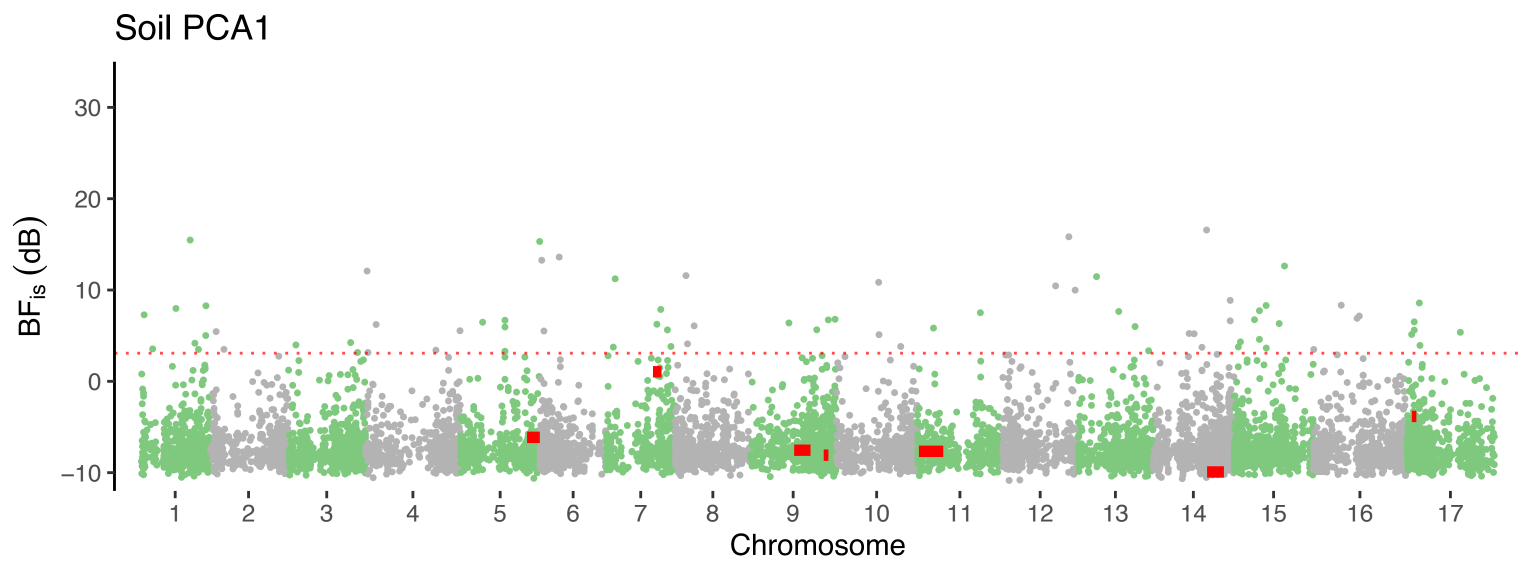

Soil PCA1
BFis (dB)
30
20
10
0
−10
Chromosome

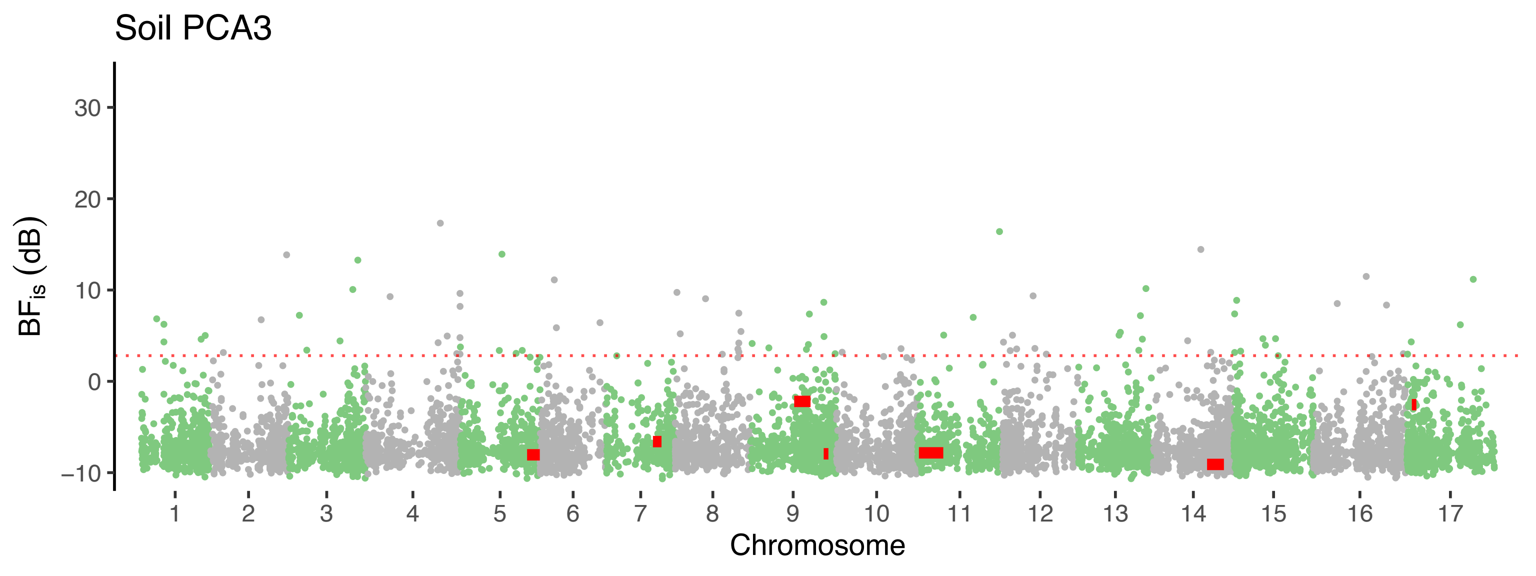

Soil PCA3
BFis (dB)
30
20
10
0
−10
Chromosome
