## Supporting Information Table S1 for "Multiple chromosomal inversions contribute to adaptive divergence of a dune sunflower ecotype"

**TABLE S1** Location and ecotype assignment of each population

| Population | Ecotype† | Latitude | Longitude |
|---|---|---|---|
| 970 | N | 37.663 | -105.625 |
| 1003 | D | 37.773 | -105.576 |
| 1033 | D | 37.787 | -105.568 |
| 1063 | N | 37.774 | -105.598 |
| 1094 | I | 37.819 | -105.616 |
| 1117 | D | 37.816 | -105.597 |
| 1147 | N | 37.836 | -105.603 |
| 1180 | D | 37.773 | -105.553 |
| 1240 | D | 37.786 | -105.531 |
| 1270 | D | 37.764 | -105.524 |
| 1300 | D | 37.745 | -105.546 |
| 1331 | I | 37.726 | -105.553 |
| 1363 | N | 37.716 | -105.533 |
| 1547 | D | 37.761 | -105.570 |
| 1701 | D | 37.804 | -105.524 |
| 1731 | D | 37.808 | -105.553 |
| 1761 | I | 37.814 | -105.532 |
| 1791 | N | 37.813 | -105.517 |
| 2001 | N | 37.757 | -105.507 |
| 2250 | N | 37.672 | -105.592 |

†N: non-dune, D: dune, I: intermediate.
